# Genome-wide screens in accelerated human stem cell-derived neural progenitor cells identify Zika virus host factors and drivers of proliferation

**DOI:** 10.1101/476440

**Authors:** Michael F. Wells, Max R. Salick, Federica Piccioni, Ellen J. Hill, Jana M. Mitchell, Kathleen A. Worringer, Joseph J. Raymond, Sravya Kommineni, Karrie Chan, Daniel Ho, Brant K. Peterson, Marco T. Siekmann, Olli Pietilainen, Ralda Nehme, Ajamete Kaykas, Kevin Eggan

## Abstract

Neural progenitor cells (NPCs) are essential to brain development and their dysfunction is linked to several disorders, including autism, Zika Virus Congenital Syndrome, and cancer. Understanding of these conditions has been improved by advancements with stem cell-derived NPC models. However, current differentiation methods require many days or weeks to generate NPCs and show variability in efficacy among cell lines. Here, we describe human <u>S</u>tem cell-derived <u>N</u>GN2-<u>a</u>ccelerated <u>P</u>rogenitor cells (SNaPs), which are produced in only 48 hours. SNaPs express canonical forebrain NPC protein markers, are proliferative, multipotent, and like other human NPCs, are susceptible to Zika-mediated death. We further demonstrate SNaPs are valuable for large-scale investigations of genetic and environmental influencers of neurodevelopment by deploying them for genome-wide CRISPR-Cas9 screens. Our studies expand knowledge of NPCs by identifying known and novel Zika host factors, as well as new regulators of NPC proliferation validated by re-identification of the autism spectrum gene *PTEN*.

## INTRODUCTION

The higher cognitive abilities of humans emerged in part from the greater neocortical volume and surface area of our brains relative to other species. These enlargements, are in turn, a product of the prolonged expansion of neural progenitor cell (NPCs) at several stages of early neurodevelopment (Homem et al., 2015; Rakic, 2009). Epithelial NPCs, which are the first specified brain cell type in the embryo, line the walls of the neural tube and eventually establish neurogenic regions known as the ventricular and subventricular zones (Alvarez-Buylla et al., 1990). Here, NPCs undergo both symmetric divisions that increase the size of progenitor pools, and asymmetric divisions that initiate neuronal and glial differentiation (Kriegstein and Alvarez-Buylla, 2009; Noctor et al., 2004). Throughout this process, proliferation of a diverse array of intermediate progenitor cell types located in the outer subventricular zone leads to the folded, gyrencephalic architecture of the human brain that differs greatly from the smooth, lissencephalic cortices of rodents and other mammalian species (Lui et al., 2011). Because of the importance of controlled NPC expansion in brain development, dysfunctional NPC proliferation and differentiation can contribute to several disorders including autism spectrum disorders (ASD), medulloblastoma, and Zika Virus (ZIKV) Congenital Syndrome, all of which are poorly understood and lack effective clinical treatments (Hemmati et al., 2003; Marchetto et al., 2017; Tang et al., 2016).

Given that samples of human brain tissue are rare and difficult to manipulate experimentally, the development and optimization of *in vitro* models of human neural cells are needed for scalable and high-throughput investigations of the molecular mechanisms that govern their behaviors, especially as they pertain to disease. Ever improving human pluripotent stem cell (hPSC) culturing methodologies show substantial promise for changing the means by which we investigate such mechanisms. Since the description of the first human embryonic stem cells (hESCs) (Thomson, 1998) and human induced pluripotent stem cells (hiPSCs) (Takahashi et al., 2007), various methods have emerged for their differentiation into NPCs and post-mitotic neurons. One common approach involves the culturing of hPSCs in suspension as embryoid bodies, which are then transferred to an adherent surface and maintained before neural rosettes are manually selected for expansion (Elkabetz et al., 2008; Koch et al., 2009; Zhang et al., 2001). Neural conversion can also be accomplished through the application of small molecule inhibitors of the SMAD and WNT signaling pathways to confluent monolayers of hPSCs (Chambers et al., 2009). These techniques produce neural cells by attempting to mimic events thought to occur *in vivo* and are attractive because of their potential developmental fidelity.

Though transformative and widely adopted, these approaches are limited in their applicability to large-scale experimentation for several reasons. First, existing protocols require between 11 and 50 days to acquire a NPC identity before additional time-consuming steps are needed to produce post-mitotic neurons (Kelava and Lancaster, 2016). Second, these methods show variability in efficacy, in which rounds of differentiation can either fail outright or yield heterogeneous cultures comprised of both neural and non-neural cell types (Muratore et al., 2014). These inconsistencies can significantly impact experiments that involve a single cell line (i.e. run-to-run/batch effects) or multiple lines that are being neuralized simultaneously. In total, these issues can occlude potentially relevant phenotypic differences between cell lines, prevent scaling to sufficient cell numbers for unbiased chemical or genetic screening, and contribute to poor reproducibility of findings across laboratories (Sandoe and Eggan, 2013).

More recently, stem cell differentiation strategies relying on the forced expression of neuralizing transcription factors, such as Neurogenin-2 (NGN2), have emerged (Nehme et al., 2018; Zhang et al., 2013). These NGN2-induced neurons are known to form functional synaptic networks in as little as 4 weeks, and have been proven useful in attempts to probe the disease mechanisms underlying several neurodevelopmental disorders (Merkle and Eggan, 2013; Yi et al., 2016). These “induced-neurons” appear to be more homogenous and offer a level of scalability not afforded by previous methods. However, the relevance of these induced neuronal models to human disease mechanisms has been called into question due in part to the uncertainty regarding the developmental trajectory they follow relative to neurons *in vivo* and whether this renders the resulting neuronal product less physiologically accurate. Specifically, whether or not these cells transition through a functional NPC stage en route to their final neuronal destination is debatable and has thus far not been thoroughly addressed.

Here, we report on a scalable and reliable method for producing stable cultures of human stem cell-derived NPCs through a simple 48-hour induction protocol. These cells, which we have dubbed *<u>S</u>tem cell-derived <u>N</u>GN2-<u>a</u>ccelerated <u>P</u>rogenitor cells* (SNaPs), satisfy the biochemical and functional requirements for NPC classification, thus demonstrating that NGN2-based neuralizing schemes do in fact proceed through a progenitor phase. SNaPs are multipotent cells capable of differentiating into both glia and neurons, as shown by single-cell RNA sequencing (scRNA-seq), immunostaining, and electrophysiology. Clonal assay experiments revealed that SNaPs are proliferative and capable of self-renewal, while SNaPs cultured in suspension demonstrate an ability to self-aggregate into three-dimensional neurospheres. Consistent with the observed cell tropism of ZIKV for NPCs, SNaPs are highly susceptible to ZIKV infection and viral-mediated cell death and can therefore act as a model of flavivirus neuropathogenesis. Importantly, this rapid induction method was successful in a large number of hES and hiPS cell lines, underscoring its reproducibility and broad potential to accelerate future *in vitro* investigations of human brain development. Finally, leveraging the speed and utility of the SNaP system, we conducted a genome-wide CRISPR-Cas9 ZIKV survival screen with a built-in fitness assay that identified new viral host factors as well as novel regulators of NPC proliferation and viability.

## RESULTS

### Brief pulse of NGN2 overexpression in hPSCs produces human NPCs

Overexpression of NGN2 has previously been shown to efficiently induce excitatory glutamatergic neurons from hPSCs (Zhang et al., 2013). We recently showed this method could be made more reliable through the addition of small molecules that further pattern these differentiating cells to more-defined cell fates (Nehme et al., 2018). To develop a sense of whether induced neurons might pass through a progenitor cell phase, we analyzed our recently published RNA sequencing (RNA-seq) data collected from cells harvested at several time points during NGN2 induced differentiation. We discovered that the expression of progenitor cell and pan-neuronal transcript markers increased over the first two days of differentiation, while levels of pluripotency genes quickly diminished relative to human PSCs (Figure 1a, Figure S1a). Transcripts known to be expressed in forebrain progenitor cells peaked at Day 4 before sharply decreasing at Day 14, when cells instead expressed transcripts found in immature post-mitotic neurons. Based on these observations, we concluded that induced neurons might indeed pass through a progenitor-like state in the earliest stages of differentiation. Consistent with this notion, the cellular proliferation marker *MKI67*, which is present in actively cycling cells and also highly expressed in NPCs reached maximal levels 2 days after NGN2 induction, before decreasing over remaining time points (Figure S1a). While these expression data support the notion that differentiating cells might be transiting through a progenitor state, we next sought to functionally substantiate the presence of these putative NPCs by limiting the NGN2 induction to 48 hours and then shifting the resulting cells to standard culture conditions supporting NPC self-renewal.

**Figure 1:**
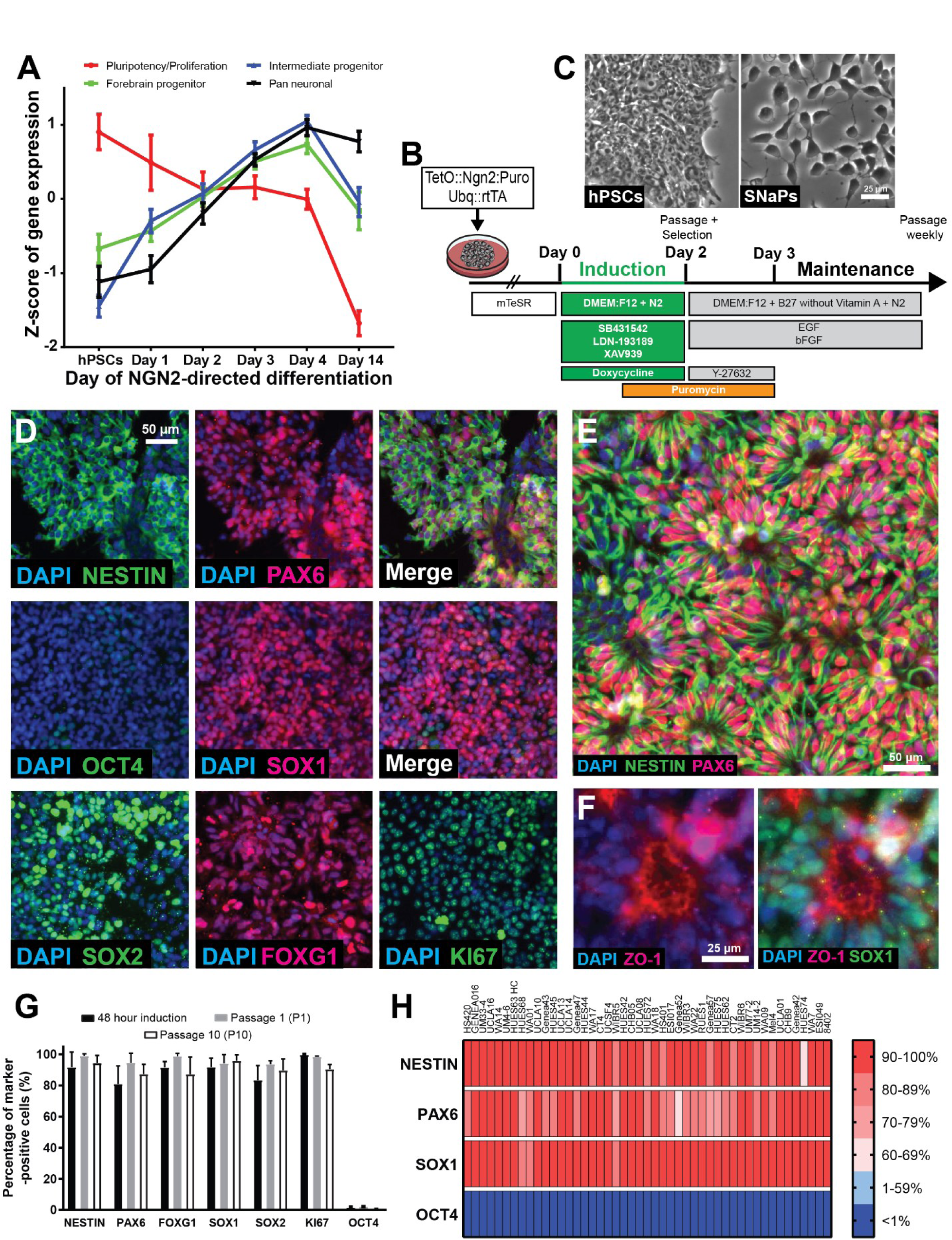
Rapid induction of stem cell-derived human neural progenitor cells. **A**, Normalized expression of pluripotency, progenitor, and neuronal transcript markers over the course of NGN2-directed differentiation of induced neurons (from Nehme et al., 2018). **B**, Schematic describing the SNaP induction and expansion protocol. **C**, Bright field images of SW.1 induced pluripotent stem cells (left) and SNaPs at 48 hours post-induction (right). Scale bar = 25 μm. **D**, SNaP immunostaining of forebrain NPC protein markers at 48 hours post-induction. Scale bar = 50 μm. **E**, SNaPs self-organize into rosette-like structures 2 days after first passage. Scale bar = 50 μm. **F**, Magnified images of a SOX1-positive rosette structure. Scale bar = 25 μm. **G**, Quantification of protein marker expression in SNaPs at 48 hours post-induction, 2 days after first passage, and after passage 10 (n = 8-24 wells from 2-3 differentiations). **H**, Heat map depicting percentage of NPC marker-positive cells in P1-2 SNaPs from 47 different hPSC lines (n = 4 wells). Data are represented as mean ± S.D.

To this end, we transduced hPSCs with lentiviruses encoding TetO::*NGN2:T2A:*PURO and Ubq::rtTA to allow for the DOX-inducible overexpression of NGN2 and the linked puromycin resistance gene (Figure 1b). After expansion of the transduced hPSCs, we initiated NGN2 induction by adding DOX and the small molecule inhibitors of the SMAD (LDN-193189 and SB431542) and WNT (XAV939) signaling pathways, which are known to direct early neural cells to a more dorsalized fate (Hemmati-Brivanlou and Meltont, 1997; Smith et al., 2008). After only 24 hours, puromycin was added to the media to eliminate non-transduced cells. Forty-eight hours post-induction, these cells (hereby referred to as SNaPs) showed a bipolar morphology that is characteristic of human NPCs (Figure 1c) and expressed early forebrain progenitor protein markers PAX6 (80.95%), NESTIN (91.65%), and FOXG1 (91.56%) (Figure 1d-e), which were absent in hPSCs (Figure S1b). We found these SNaPs also expressed SOX1 (91.70%) and SOX2 (83.45%), consistent with a neural stem cell identity, in addition to the aforementioned proliferation marker MKI67 (99.66%). A higher percentage of SNaPs stained positive for these proteins after passage and an additional 24 hours of puromycin selection (PAX6: 94.48%; NESTIN: 99.02%; FOXG1: 98.72%; SOX1: 94.13%; SOX2: 93.7%). Consistent with an exit from pluripotency, we found expression of OCT4 declined precipitously over 48 hours of NGN2 induction as evident by both immunostaining and fluorescence-activated cell sorting (FACS) analyses (Figure 1d, Figure S1c).

To determine if these apparent NPCs were stable and expandable, we passaged the SNaPs into basic fibroblast growth factor (bFGF) and epidermal growth factor (EGF)-containing media supplemented with puromycin and ROCK pathway inhibitor (Y-27632), the latter of which was added to reduce the cell death induced by single-cell passage. The following day, both the ROCK inhibitor and puromycin were removed. At 2 days post-passaging (Passage 1, P1), we observed homogenous cultures of SNaPs that self-organized into neural rosette structures with prototypical apical redistribution of ZO-1 protein localization and expressed many proteins normally found in NPCs as judged by immunostaining (Figure 1f). Furthermore, SNaPs could be serially passaged while maintaining a NPC phenotype (Passage 10, P10; Figure 1g). These data were supported by qPCR analysis of samples harvested from hPSCs and SNaPs at different stages of culture development, which showed that SNaPs exhibited a marked increase in *PAX6*, *FOXG1*, and *SOX1* levels relative to hPSCs at 48 hours post-induction while *OCT4* declined, a pattern that held steady in older cultures (Passages 1 and 6; Figure S1d). Importantly, the induction efficiency of the SNaP method was independent of initial seeding density, which is an improvement upon conventional SMAD inhibition methods that require high hPSC confluency for successful neural conversion (Figure S1e).

As initial experiments were all conducted on the SW.1 hiPS cell line, we wanted to determine whether SNaP induction was effective across a wide range of genetic backgrounds. We generated SNaPs from dozens of different hiPS and hES cell lines (n = 48). Demonstrating the method’s reproducibility, we found that when using ≥75% NESTIN^+^/PAX6^+^/SOX1^+^ and ≤ 0.1% OCT4^+^ cells at Passage 1-2 as our threshold, our approach was successful in 46 of 48 (95.83%) hiPSC and hESC lines on our first attempt (Figure 1h; Table S1).

Next, we sought to compare the cellular identity of SNaPs to hPSC-derived NPCs generated using SMAD/WNT inhibition approaches. The SMAD/WNT NPCs were generated without DOX-induced NGN2 overexpression using 2 weeks of LDN-193189, SB431542, and XAV939 small molecule treatment (*LSX-2w*) or LDN-193189, SB431542, and Retinoic acid treatment for 3 weeks (*LSR-3w*). The transcript profiles of P1 SNaPs and NPCs produced using the LSX-2w and LSR-3w protocols were analogous (Figure S1f), and normalization of SNaP samples to LSX-2w showed convergence of NPC marker expression levels (Figure S1g). This data demonstrates the similarities between SNaPs and conventional stem cell-derived NPCs.

### SNaPs can self-renew and differentiate into neurons and glia

The ability to differentiate into multiple cell types in the neural and glial lineages is a hallmark of NPCs. To assess SNaP multipotency, we plated cells at a low density (15,000 cells/cm^2^) in growth factor-free media (base differentiation media), which is known to stimulate spontaneous differentiation. Two weeks later, we harvested the resulting cells and performed single-cell RNA-seq (scRNA-seq) analysis to assess the expression profiles of 2,167 SNaP-derived cells. We found that SNaPs spontaneously differentiated into several different cell types that could be clustered into 7 groups (Figure 2a, Figure S2a-b). Group 2 cells expressed high levels of *MKI67*, suggesting that they remained highly proliferative (Figure 2b). A subset of cells (Group 5, 12.7%) were PAX6^+^/FOXG1^+^/FEZF2^+^/EMX2^+^, which seemed indicative of phenotype similar to that found in telencephalic regional progenitors (Figure 2c). Another cluster of cells was characterized by expression of *STMN2* and *ELAVL4* (Group 3, 13.7%), suggesting that SNaPs can spontaneously differentiate into post-mitotic neurons. A small percentage of cells (5.8%) did not cluster into any of the 7 groups and were therefore termed “unclassified”.

**Figure 2:**
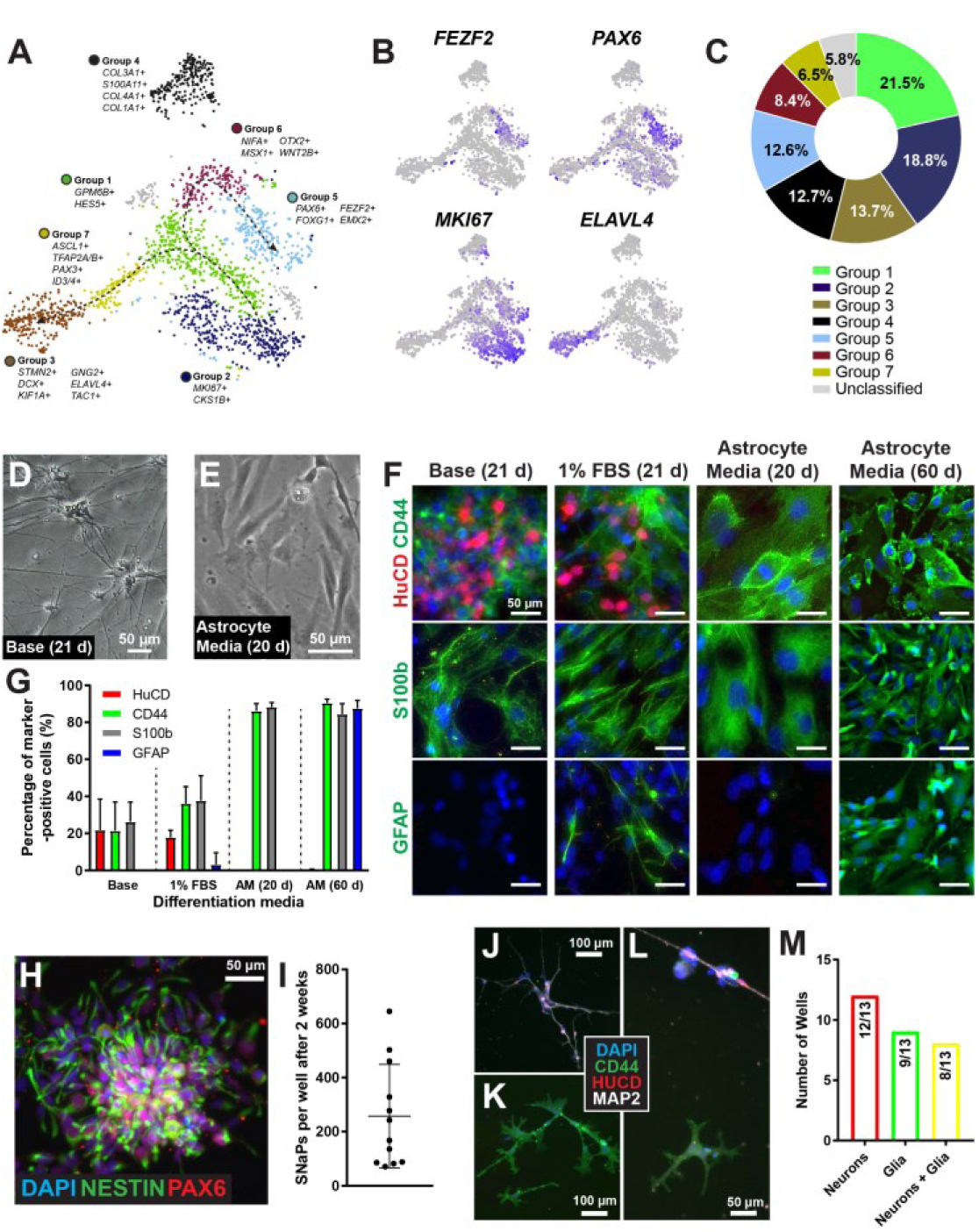
SNaPs are multipotent and capable of self-renewal. **A-B**, 10X single-cell RNA sequencing cluster plot reveals the presence of multiple cell types after 2 weeks of spontaneous differentiation. **C**, Quantification of single-cell RNA sequencing results. **D-E**, Bright field images of (D) SNaP-derived post-mitotic neurons after 3 weeks in base media and (E) glial cells after 20 days in Astrocyte Media (AM). **F**, Immunostaining of neuronal (HuC/D) and glial (CD44, S100b, and GFAP) protein markers in base media or 1% FBS media after 2-3 weeks in culture or after 20-60 days in AM. **G**, Quantification of images from Panel F (n = 4-6 wells per condition). **H-M**, Clonal assay reveals that SNaPs plated as single cells can self-renew. (H) Representative image of a cluster of PAX6/NESTIN-positive cells taken 14 days after plating of a single SNaP (n = 12 wells). (I) Quantification of number of cells per well after two weeks of proliferation. Representative images of single-cell SNaP-derived (J) HuC/D-positive neurons, (K) CD44-positive glia, and (L) neurons+glia co-culture in base media at 14 days post-differentiation. (M) Quantification of number of neuron, glia, and neurons+glia containing wells. Data are represented as mean ± S.D.

To more formally test the potency of SNaPs to differentiate into both astroglial and neuronal lineages, we cultured SNaPs under conditions designed to promote NPC differentiation into each lineage and assessed the resulting derivatives by immunocytochemistry. We found that spontaneous differentiation of these cells in base media for three weeks resulted in a mixture of HuC/D^+^ (the protein product of genes *ELAVL3/4*) neurons and CD44^+^/S100b^+^ glia cells (Figure 2d, f-g). Supplementing the base media with 1% fetal bovine serum (FBS) to promote glial differentiation, yielded both HuC/D^+^ neurons and CD44^+^/S100b^+^ glia, with a small percentage of the glial cells staining positive for the astrocytic marker GFAP (Figure 2f-g). The use of astrocyte media (AM), which was recently shown to effectively convert hPSCs to mature glia (TCW et al., 2017), produced nearly pure cultures of CD44^+^/S-100b^+^ glial cells after 20 days in culture, and CD44^+^/S-100b^+^/GFAP^+^ glial cells after 60 days (Figure 2e-g). Together, these results suggest that SNaPs can differentiate into both neurons and glia, and that the differentiation of these cells can be readily achieved with the extrinsic factors widely used to control NPC differentiation.

To determine if SNaPs are capable of self-renewal, which is defined by the ability of a cell to divide while maintaining a multipotent state and is a characteristic of neural stem cells, we performed single-cell clonal analysis. After plating SNaPs at one per well, we counted an average of 256.58 cells in each well after two weeks in bFGF/EGF-containing media (Figure 2h-i). When these proliferating SNaPs were again re-plated as single cells in individuals wells and cultured in base media for three weeks, we observed spontaneous differentiation into neurons and glia in multiple wells derived from a single clonal progenitor (Figure 2j-m), thus exhibiting the multipotency and self-renewal capabilities of SNaPs.

### SNaP-derived neurons form active networks

Differentiated excitatory neurons *in vitro* are capable of forming synaptic connections and through them displaying network activity. To determine if SNaP-derived neurons could reach this level of functionality, we first stained 30-day-old cultures with protein markers of differentiated superficial and deep layer cortical glutamatergic neurons. We found that a majority of the post-mitotic HuC/D^+^ neurons co-expressed superficial layer markers BRN2 and CUX2 (85.77% and 82.90%, respectively; Figure S3a-b). A small percentage of HuC/D^+^ cells co-expressed the deep layer marker CTIP2 (3.70%). Immunostaining of 50-day-old SNaP-derived neurons co-cultured with primary mouse glia revealed characteristic punctate expression of the synaptic marker Synapsin I (Figure S3c), suggesting that SNaPs can spontaneously differentiate into neurons that display some features found in cortical neurons.

We next proceeded to use multi-electrode arrays (MEA) to assess the ability of SNaP-derived neurons co-cultured with primary mouse glia to produce action potentials and form functional synaptic networks (Figure 3a-b). Network activity was monitored over the course of 14 weeks. The mean number of active electrodes increased over the first several weeks relative to baseline, with activity recorded on all but a small number of electrodes by week 8 (p < 0.001; Figure 3c). The mean firing rate followed a similar time course, reaching a peak frequency of 16.9 Hz by week 12 (p = 0.0492; Figure 3d). Burst analysis of the MEA recordings identified synchronous firing events with a network burst rate that reached statistical significance relative to baseline at five weeks of differentiation (0.138 Hz, p = 0.0058; Figure 3e-f).

**Figure 3:**
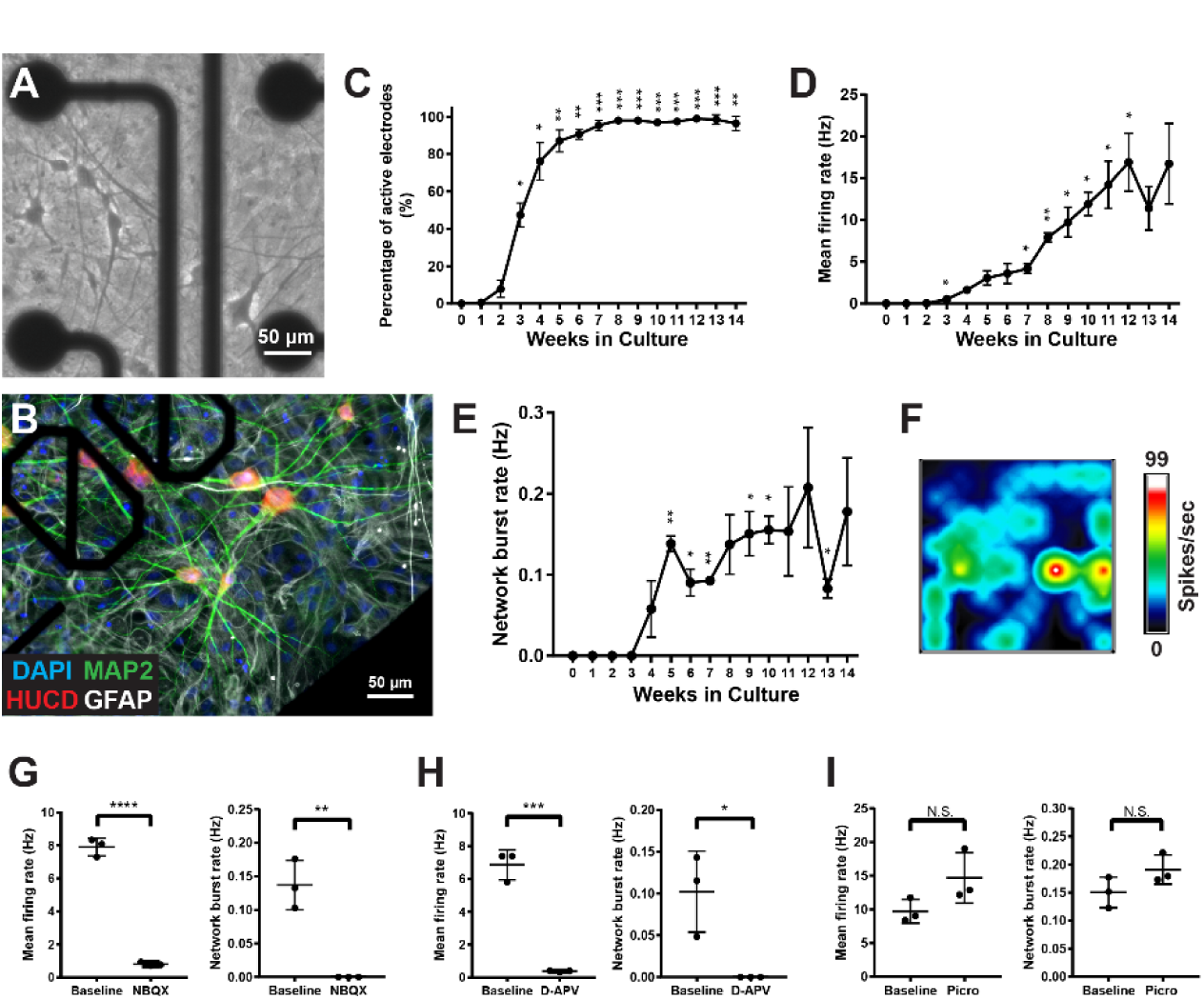
SNaP-derived neurons form synaptic networks. **A**, Bright field image of 24-day-old SNaP-derived neurons co-cultured with primary mouse glia on a muti-electrode (MEA) well. Scale bar = 50 μm. **B**, SNaP-derived neurons co-cultured with mouse glia on a MEA well at 100 days post-plating. **C**, Percentage of active MEA electrodes plateaus after 7-8 weeks in co-culture. **D**, Firing neurons reach a peak mean firing rate after 12-14 weeks. **E-F**, SNaP-derived neurons form functional networks that can fire in synchronous bursts, as shown in (E) the graph depicting increasing network burst firing rates over time and (F) representative heat map of active electrodes. **G-H**, Network activity of SNaP-derived neuronal cultures is affected by (G) AMPA receptor antagonist NBQX (10 μM) or (H) NMDA receptor antagonist D-APV (50 μM) at Day 50 post-plating. I, GABA_A_R antagonist picrotoxin (50 μM) has no effect. Repeated measure one-way ANOVA with Dunnett’s tests for multiple comparisons (C-E) and Student’s t-test (G-I) used for statistical analysis. For (C-E), statistical comparisons were made with baseline recordings (Week 1). N = 3 wells for all experiments. Data are represented as mean ± S.D. N.S. = not significant, *p<0.05, **p<0.01, ***p<0.001, ****p<0.0001.

We employed a pharmacological approach to better understand whether synaptic components contributed to the network activity we observed. The addition of the AMPA receptor (AMPAR) competitive antagonist NBQX (10 μM) at Day 56 significantly decreased the mean firing (p < 0.0001) and mean bursting (p = 0.0029) rates compared to vehicle-treated controls (Figure 3g), suggesting that network activity is driven in part by ionotropic glutamate receptor-mediated excitatory synaptic transmission. Similarly, the NMDA receptor (NMDAR) selective competitive antagonist D-AP5 (50 μM) significantly reduced network activity (firing: p = 0.0003; bursts: p = 0.0219; Figure 3h). Treatment with GABA_A_ receptor antagonist picrotoxin (50 μM) did not alter mean firing or bursting rates (firing: p = 0.1065; bursts: p = 0.1362; Figure 3i), indicating that the activity of inhibitory neurons did not significantly impact the network dynamics of the SNaP-derived cultures that appear to be predominantly populated with excitatory neurons. Together, these results demonstrate the ability of SNaPs to spontaneous differentiate into functional neurons, and in doing so, further supports the classification of SNaPs as human NPCs.

### SNaPs self-aggregate into neurospheres in suspension culture

Neural progenitor cells self-aggregate in suspension culture into three-dimensional structures knowns as neurospheres (Reynolds and Weiss, 1992). Early studies found that NPCs within neurospheres were proliferative and capable of differentiating into post-mitotic neural cell types (Reynolds and Weiss, 1992). This pioneering work spawned the development of assays that measure self-renewal, neuronal migration, and neurite outgrowth (Brennand et al., 2015). To determine if SNaPs can form neurospheres, we cultured dissociated SNaPs in low-attachment 96-well U-shaped plates with bFGF/EGF-containing maintenance media and high concentrations of ROCK inhibitor (Figure 4a) (Salick et al., 2017). Starting at Day 3 of the procedure, we exchanged media every other day and then recorded the two-dimensional area of the SNaP-derived neurospheres (“SNaPspheres”; Figure 4b). We found that SNaPspheres from SW.1 hiPSCs grew to an average size of 1.74 mm^2^ over the course of 14 days and reached 3.70 mm^2^ by Day 36 post-aggregation. SNaPspheres from the H9 hESC line showed similar growth dynamics (Figure 4c-d). Lightsheet imaging of intact day 7 SNaPspheres immunostained with the NPC protein marker phospho-Vimentin (phVim) and the neurite marker MAP2 showed that both progenitors and early neuronal cells were present (Figure 4e).

**Figure 4:**
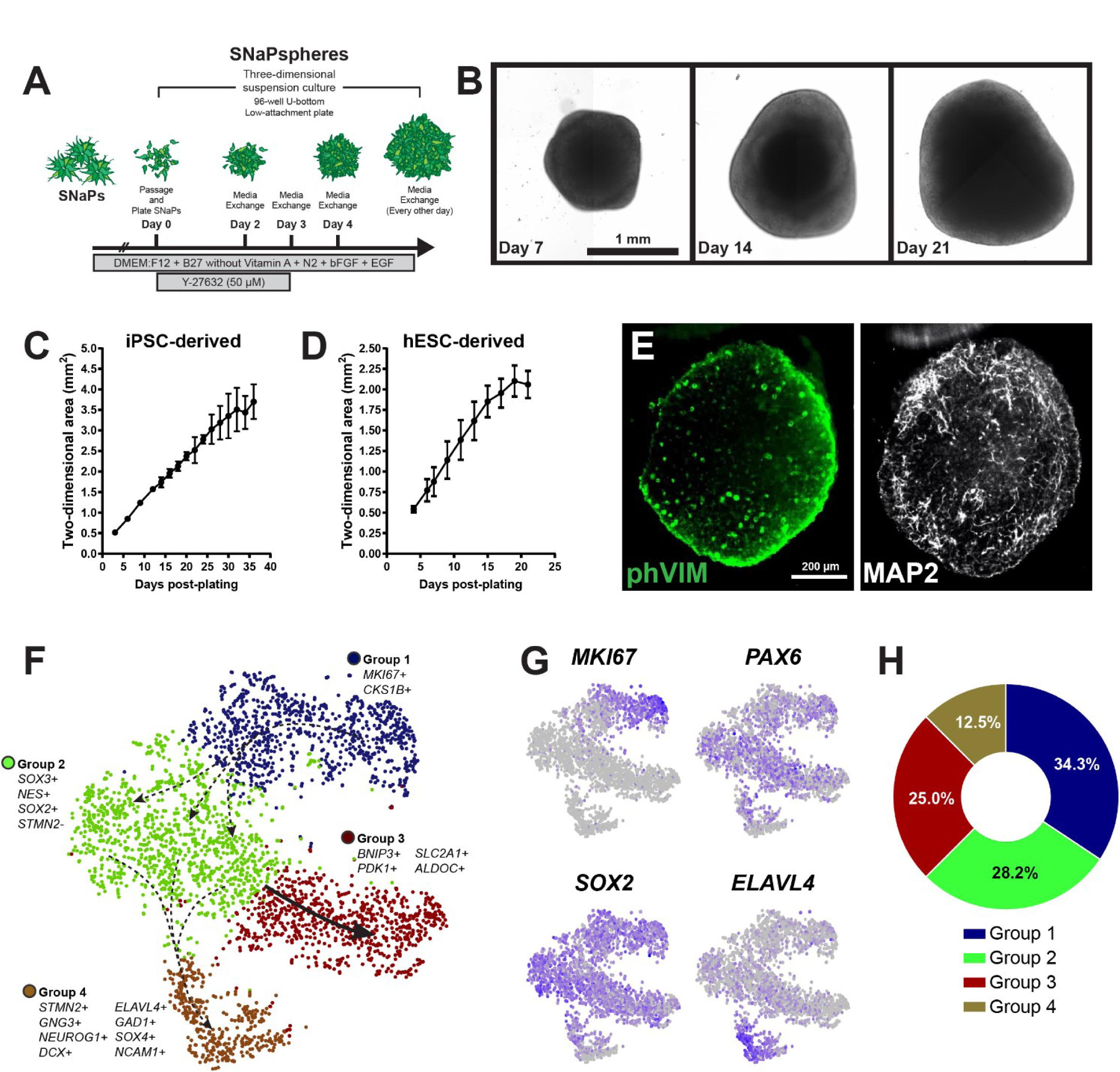
SNaPs self-aggregate into three-dimensional neurospheres. **A**, Schematic describing the production of three-dimensional SNaPspheres from two-dimensional SNaP cultures. **B**, Bright field image of SNaPspheres at days 1, 7, and 21 post-plating. Scale bar = 1 mm. **C-D**, Two-dimensional area of SNaPspheres derived from (C) SW.1 hiPSC SNaPs (n = 12 spheres) or (D) H9 hESC SNaPs (n = 8 spheres). **E**, Lightsheet imaging of a Day 7 SNaP-neurosphere stained with phospho-Vimentin (NPC marker) and MAP2 (neurite marker). **F-G**, Groups of cells within 7-day-old SNaPspheres as revealed by 10X single-cell RNA sequencing cluster analysis. **H**, Quantification of Panel F. Data are represented as mean ± S.D.

Our conclusion that both self-renewal and differentiation were occurring in these neurospheres was supported by scRNA-seq analysis, which found that approximately 12.5% of the cells expressed transcripts enriched in post-mitotic neurons such as *ELAVL4* and *STMN2* (Group 4) while the remaining were either *MKI67^+^* mitotic cells (Group 1, 34.2%) or quiescent progenitor cells that expressed *PAX6* and *SOX2* (Group 2, 28.3%) (Figure 4f-h, Figure S4a-b). A subset of the quiescent state progenitors (Group 3, 25.0%) expressed genes that are typically upregulated under hypoxic conditions, such as *BNIP3*, *PDK1*, and *SLC2A1*. It is likely that these cells were located in the core of the SNaPsphere where oxygen and other nutrients are less accessible than in perimeter regions. Together, these findings highlight the utility of SNaPspheres for three-dimensional modeling of NPC behaviors.

### Application of SNaPs to study Zika virus neuropathogenesis

ZIKV is a positive-strand RNA virus that is in the same flaviviridae family as West Nile (WNV), Dengue (DENV), and Yellow Fever viruses (YFV). Transmitted primarily by mosquitoes, ZIKV is a growing and likely recurrent global health concern punctuated by outbreaks in French Polynesia (2013-2014), the Americas (2015-2016), and most recently India (October 2018) (Rossi et al., 2018; Saxena et al., 2018). Prenatal ZIKV infection has been linked to several congenital brain abnormalities including microcephaly, intracranial calcifications, and fetal demise (Brasil et al., 2016; Mlakar et al., 2016). Longer-term monitoring of children prenatally exposed to ZIKV showed that 1 in 7 suffered from likely co-morbidities that ranged from sensory and motor impairments to seizures (Rice et al., 2018). ZIKV differs from other flaviviruses in that it preferentially infects NPCs and glial cell types while mostly sparing hPSCs and post-mitotic neurons (Tang et al., 2016). Based on these observations, we used susceptibility to ZIKV infection as an independent validator of SNaP cellular identity. We used two strains of ZIKV, the African MR-766 strain isolated from a sentinel monkey in Uganda in 1947 (ZIKV-Ug) and a more recent American strain harvested from a patient in Puerto Rico (ZIKV-PR). Immunostaining for ZIKV envelope protein (4G2) conducted 54 hours post-infection (hpi) showed that SNaP monolayers could be infected with ZIKV-Ug (Figure 5a), a finding that was also evident in PAX6^+^ SNaP rosettes (Figure 5b). SNaPs were vulnerable to both the African and American strains of ZIKV, with 79.6% and 38.5% of these cells expressing detectable levels of 4G2 when infected with ZIKV-Ug or ZIKV-PR, respectively, at a multiplicity of infection (MOI) of 10 (Figure 5c-d). ZIKV infection led to a dose-dependent decrease in SNaP viability by 96 hpi. Notably, ZIKV-Ug infection resulted in significantly greater levels of cell death compared to ZIKV-PR at all MOIs (p < 0.0001; Figure 5e), which is consistent with other reports detailing the increased virulence of this African strain relative to modern American and Asian ZIKV isolates in both culture and mice (Simonin et al., 2016, 2017). Furthermore, quantification of ZIKV RNA levels in the conditioned media of infected SNaPs revealed a time-dependent increase in viral titer, suggesting that SNaPs were capable of supporting the replication of ZIKV particles (ZIKV-Ug: p = 0.0007; ZIKV-PR: p = 0.0005; Figure 5f). Immunostaining of ZIKV-sensitive Vero cells 24 hours after exposure to the conditioned media from infected SNaPs confirmed that the ZIKV particles propagated in SNaPs were infectious (Figure 5g).

**Figure 5:**
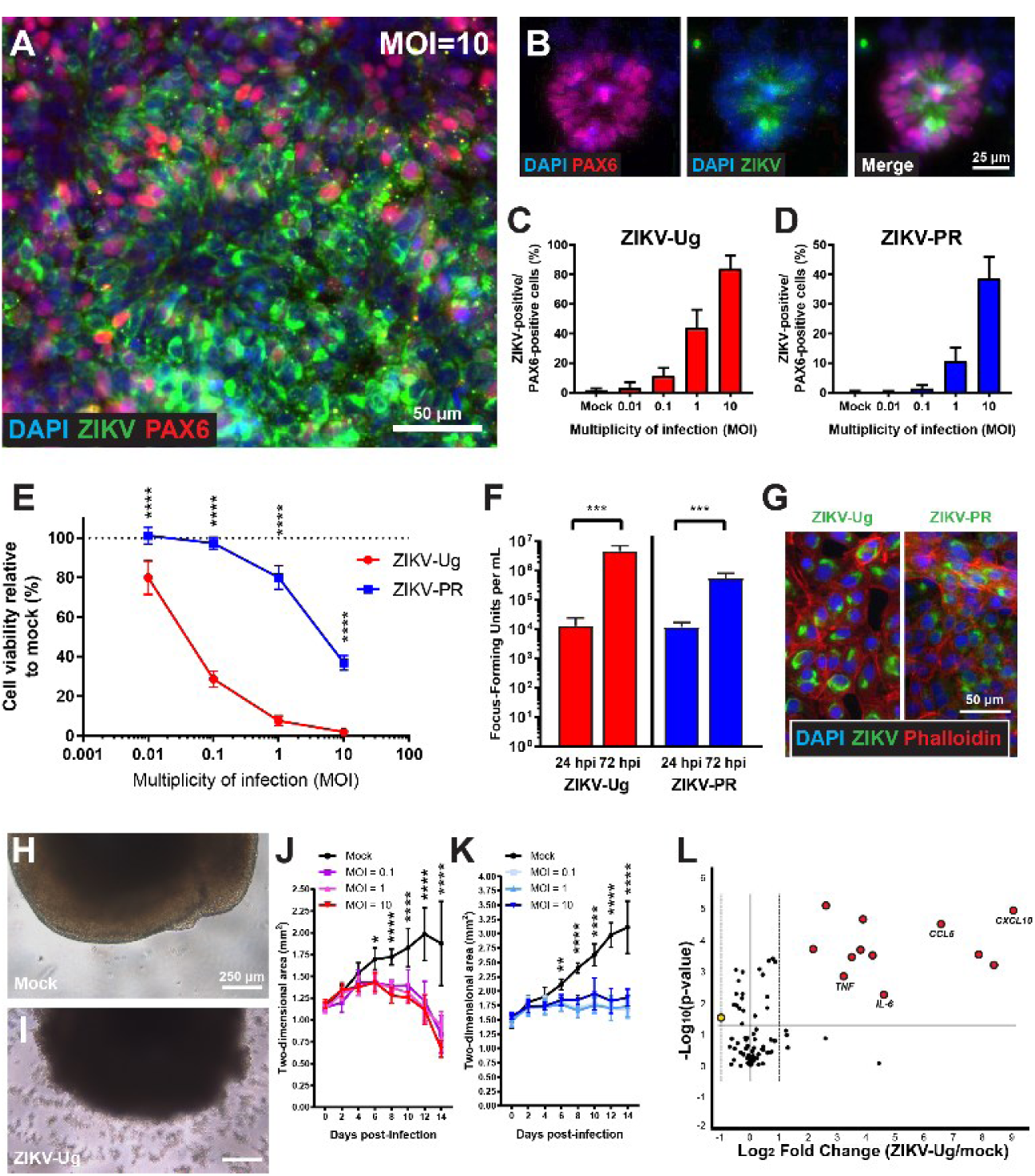
2-D and 3-D SNaPs are susceptible to ZIKV infection. **A**, Immunostaining of ZIKV 4G2 envelope protein at 54 hours post-infection (hpi) shows that ZIKV-Ug (MOI = 10) can infect a high percentage of PAX6-positive SNaPs. Scale bar = 50 μm. **B**, ZIKV-Ug (MOI = 1) infection of PAX6-positive neural rosette. Scale bar = 25 μm. **C-D**, Quantification of SNaP infections with (C) ZIKV-Ug and (D) ZIKV-PR at 54 hpi (n = 8 wells). **E**, Cell viability of SNaPs infected with ZIKV-Ug (red line) or ZIKV-PR (blue line) at 96 hpi. Viability is normalized to mock-infected controls (n = 8 wells). **F**, Quantification of ZIKV-Ug and ZIKV-PR viral RNA from conditioned media of infected (MOI = 10) SNaPs at 24 hpi and 72 hpi (n = 6 wells). **G**, ZIKV 4G2 envelope protein immunostaining of Vero cells at 24 hpi show that SNaP-infected conditioned media contain infectious ZIKV particles. F-actin Phalloidin dye used to outline cells. Scale bar = 50 μm. **H-I**, Bright field images of (H) mock or (I) ZIKV-Ug (MOI = 1) infected SNaPspheres at 8 days post-infection (dpi). Scale bar = 250 μm. **J-K**, Quantification of SNaPsphere two-dimensional change in area over time in response to (J) ZIKV-Ug or (K) ZIKV-PR infection compared to mock controls (n = 6 spheres). **L**, qPCR analysis of human antiviral response genes shows upregulation of multiple cytokines and chemokines (red dots) in SNaPs at 60 hpi of MOI = 10 ZIKV-Ug infection (n = 4). Two-way ANOVA with Sidak’s tests for multiple comparisons (E), Student’s t-test (F), and repeated measures Two-way ANOVA with Dunnett’s tests for multiple comparisons (J-K) were used for statistical analysis. Data are represented as mean ± S.D. **p<0.01, ***p<0.001, ****p<0.0001.

Given that glia cells are also targets of ZIKV neuropathogenesis, we tested the vulnerability of SNaP-derived glial cells to ZIKV infection. SNaPs were differentiated in astrocyte media for 2-3 weeks and then exposed to virus. Viral administration led to infectivity levels of 57.83% and 59.47% at 54 hpi in response to MOI = 10 of ZIKV-Ug and ZIKV-PR, respectively, in 4G2^+^/CD44^+^ putative astrocytes (Figure S5a-c). We observed decreases in astrocyte viability relative to mock controls at 96 hpi (Figure S5d). Modest, though statistically-significant differences in viral-mediated cell death were observed between the two ZIKV strains, but only at low MOIs (p = 0.024 for MOI = 1; p < 0.0001 for MOI = 0.1). Furthermore, we found that SNaP-derived glia could support ZIKV replication as shown by qPCR analysis of supernatant ZIKV RNA and further infection of Vero cells (Figure S5e-f). Together, these findings demonstrate the usefulness of two-dimensional SNaP cultures in investigations of ZIKV pathogenesis in cell types present early in the development of the human brain.

Several recent studies have used three-dimensional culture systems to study ZIKV and the brain, as these approaches allow for experimentation in a mixed population of neural cell types, including NPCs (Gabriel et al., 2017; Qian et al., 2016). To determine if SNaP-derived neurospheres were susceptible to ZIKV infection, we exposed 7-12 day old SNaPspheres to ZIKV-Ug and ZIKV-PR at a range of MOIs and measured their change in size relative to mock infection over the course of 2 weeks (Figure 5h-i). We found that SNaPspheres decreased from 1.19 cm^2^ to 0.67 cm^2^ (−43.44%) in response to high concentrations (MOI = 10) of ZIKV-Ug, while mock infected SNaPspheres increased from 1.15 cm^2^ to 1.88 cm^2^ (+63.08%) over this time (Mock vs MOI = 10: p < 0.0001 at Days 10-14; Figure 5j). ZIKV-PR infection also significantly attenuated SNaPsphere growth (Mock vs MOI = 10: p < 0.0001 at Days 8-14; Figure 5k).

Recent studies have demonstrated that, in addition to leading to the destruction of established neurospheres and cerebral organoids, ZIKV can inhibit the formation of these three-dimensional structures in culture (Garcez et al., 2016). We infected two-dimensional SNaP cultures with ZIKV-Ug (MOI = 1) before dissociating these cells and plating them under the low-attachment conditions necessary for neurosphere formation. Consistent with previous reports, we found that pre-infection of SNaPs with ZIKV-Ug interfered with the formation of SNaPspheres, as measured by the number of aggregates greater than 0.1 mm^2^ present after 6 days in culture (p = 0.0436; Figure S5g-h). Infection of SNaPspheres with ZIKV-Ug (MOI = 1) also led to significant deficits in a neurite outgrowth assay (p < 0.0001; Figure S5i-l).

Several previous studies have examined the impact of ZIKV on the transcriptional profile of human cells, including fibroblasts and neural stem cells, and observed prototypical responses to infection, namely activation of cytokines and chemokine expression (Hamel et al., 2015; Simonin et al., 2016). To determine the antiviral response of SNaPs to ZIKV, we harvested mock and ZIKV-infected SNaPs at 8 hpi and 60 hpi and then measured the expression levels of 84 immunity-related genes. While few changes were observed at 8 hpi, we found a significant increase in the expression of several genes, including *CXCL10*, *CCL5*, and *OAS2*, at 60 hpi in infected SNaPs when compared to time-matched mock controls (Figure 5l, Figure S5m). Many of the same induced genes (p<0.05 and fold change >2.0) were found in the ZIKV-Ug (MOI = 10) and ZIKV-PR (MOI = 20) datasets, indicating that common innate immunity pathways were activated in response to both strains (Table S2). Importantly, our findings mirror a recent report in which *CXCL10*, *CXCL11*, *FOS*, *IFNB1*, and *CCL5* were also found to be highly upregulated in hPSC-derived NPCs in response to infection with an African strain of ZIKV (Mesci et al., 2018).

### Genome-wide CRISPR-Cas9 ZIKV survival screen in SNaPs

To date, no FDA-approved treatments or vaccines have been identified that can limit ZIKV infection. In addition, the mechanisms by which ZIKV leads to microcephaly and NPC death are not well-understood. To identify ZIKV host factors in NPCs, and therefore potential future targets for therapeutic intervention, we conducted independent whole-genome CRISPR-Cas9 positive selection survival screens in SNaPs using both ZIKV-Ug and ZIKV-PR. To do so, we produced SNaPs from SW.1 hiPS cells and expanded them before transducing 100 million SNaP cells per replicate (3 replicates per ZIKV strain) with the Brunello human CRISPR knockout pooled library, which consists of 76,441 unique single-guide RNAs (sgRNAs) targeting 19,114 genes (approximately 4 guides per gene; Data File 1) as well as 1,000 non-targeting (NT) control sgRNAs, packaged in the all-in-one (Cas9 + sgRNA) LentiCRISPRv2.0 vector (Figure 6a). LentiCRISPR-expressing SNaPs underwent 8 days of puromycin selection and expansion before being passaged. Two days post-passage, SNaPs were either harvested (Day 0), infected with ZIKV-Ug (MOI = 1) or ZIKV-PR (MOI = 5), or treated with mock-infection media. Ten days later, we observed high levels of cell death in the ZIKV-infected SNaPs relative to mock controls (ZIKV-Ug: 98% cell death, ZIKV-PR: 76% cell death; data not shown). All samples were harvested and prepared for DNA extraction and sequencing before the number of sgRNA reads in each sample was counted and used for subsequent analysis. Our results showed high sgRNA coverage (<0.15% missing sgRNAs across replicates), a high correlation among replicates (Pearson r values >0.92), and the expected limited effects of NT sgRNA controls (Figure S6a-c).

**Figure 6:**
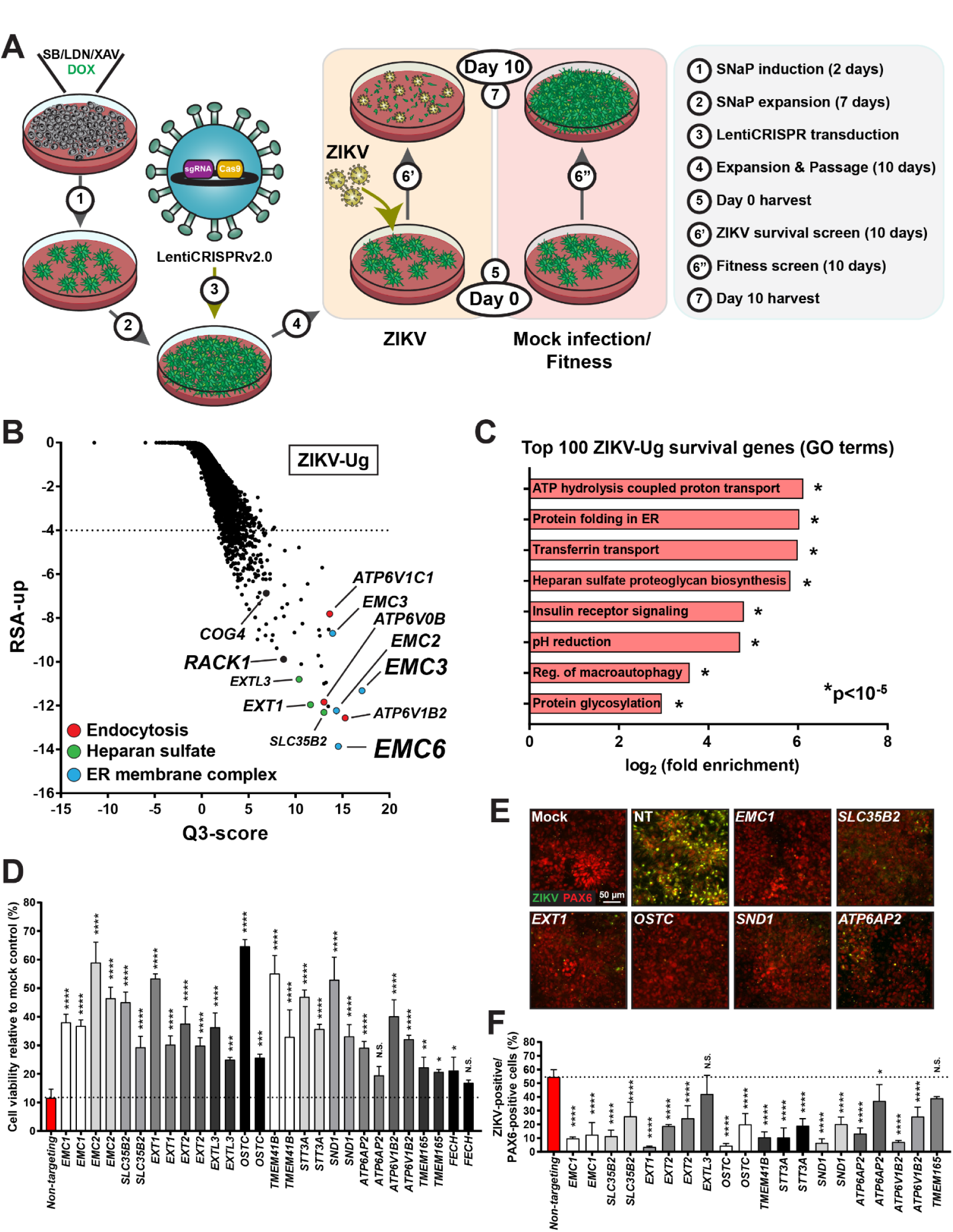
ZIKV host factors in SNaPs found through genome-wide CRISPR-Cas9 screens. **A**, Design of whole-genome CRISPR-Cas9 screens. SW.1 hiPSCs were induced to SNaPs before transduction of the Brunello sgRNA library via the LentiCRISPRv2.0 system. SNaPs used for the ZIKV survival screen were infected with ZIKV-Ug (MOI = 1), ZIKV-PR (MOI = 5), or mock infection media, and harvested 10 days later for DNA sequencing. ZIKV-infected samples were compared to Day 10 mock controls to identify genes that enhance ZIKV survival. Day 10 samples were also compared to Day 0 SNaPs to assay fitness genes. **B**, Scatter plot depicting gene level results of the ZIKV-Ug survival screen. Disruption of genes involved in the ER membrane complex (blue dots), endocytosis (red dots), and heparan sulfate biosynthesis (green dots) led to increased SNaP survival after 10 days of infection at MOI = 1. Dashed line at RSA_up_ = -4.0 denotes hit cutoff for genes enriched in Day 10 ZIKV samples relative to mock controls. **C**, GO term analysis of the top 100 RSA-scoring genes. **D-F**, Validation of primary genome-wide ZIKV survival screen. Top hits were confirmed with the (D) cell viability assay at 120 hpi and (E-F) infectivity assay at 54 hpi (n = 4 wells). Scale bar = 50 μm. Fisher’s exact test (C) and one-way ANOVA with Dunnett’s test for multiple comparisons (D, F) were used for statistical analysis. Data are represented as mean ± S.D. N.S. = not significant, *p<0.05, **p<0.01, *** p<0.001, ****p<0.0001.

The enrichment of sequencing reads for a sgRNA targeting a particular gene in the Day 10 ZIKV-infected samples relative to mock controls would suggest that disruption of said gene confers protection from ZIKV-mediated cell death. To identify such genes in SNaPs, we analyzed our sequencing results using a statistical method originally developed for siRNA screening, redundant siRNA activity (RSA) scores, which is a probability-based measure for the performance of RNAs targeting a given gene within a library to determine the gene’s impact on a screening assay (König et al., 2007) (Figure 6b). We detected 283 significantly-enriched candidate host factor genes in the ZIKV-Ug samples relative to mock controls (RSA_up_ < -4.0, adjusted p-value < 0.05; Data File 2). When we subjected the top 100 ranked genes from this screen to gene ontology (GO) analysis, we discovered enrichment of biological terms related to ER protein folding, heparan sulfate proteoglycan biosynthesis, and protein glycosylation (Figure 6c, Data File 2). A subset of these genes are involved in mechanisms with previously established connections to ZIKV infection and replication, including the oligosaccharyltransferase (*OSTC* and *STT3A*) and endoplasmic reticulum membrane complexes (*EMC1*-*7*), heparin sulfate biosynthesis (*EXT1*, *SLC35B2*, *B4GALT7*), and endocytosis (*ATP6AP2*, *ATPV1B2*, *ATP6V0B*), thus externally validating our screen (Savidis et al., 2016a).

At the same time, our analysis returned hits not previously associated with ZIKV pathogenesis. When we compared our results to a recently published genome-wide ZIKV-Ug survival screen conducted in HeLa cells (Savidis et al., 2016a), we found that of the top 100 scoring genes in these screens, only 15 were found in both datasets (Figure S6d). EMC and OSTC genes were identified as ZIKV host factors in both screens, indicating that these biological processes are universally involved in ZIKV-mediated cell death regardless of cell type. However, of the 85 non-overlapping genes found in the SNaP dataset, 10 are vacuolar ATPase (vATPase) subunits and many are involved in heparan sulfation (*SLC35B2*, *B4GALT7*, *B3GALT6*, *EXT2*). In addition, genes that comprise the oligomeric Golgi complex (*COG3*, *COG4*, *COG8*, *COG1*), which is known to play a role in the replication cycle of other brain-infecting viruses (Brass et al., 2008), and *RACK1*, which regulates translation at viral internal ribosome entry sites (Majzoub et al., 2014), were significantly enriched in our screen but went undetected in the HeLa cell screen. Overall, our screen detected known ZIKV host factors, suspected viral host factors not previously associated with flaviviruses, and several candidate host factors with no elucidated role in viral mechanisms, including *TIMM29*, integrin beta subunit *ITGB5*, and kinase-like protein *C3ORF58* (also known as *DIA1* or deleted in autism-1).

The loss of certain genes could also render cells more susceptible to ZIKV-mediated cell death. To identify such genes, we used RSA analysis to determine which were most depleted in the ZIKV-Ug screen. There were 96 genes that reached genome-wide significance, including many critical to the interferon-mediated antiviral response such as *IFNAR1-2*, *STAT2*, and *IFITM3* (RSA_down_ < -4.0, adjusted p-value < 0.05; Figure S6e, Data File 2). These results are in agreement with recent investigations on the protective measures taken by cells to fight ZIKV infection (Savidis et al., 2016b).

We also performed a ZIKV-PR survival screen in parallel to identify potential explanations for the strain differences in ZIKV-mediated SNaP death. We identified 69 significantly-enriched genes in the ZIKV-PR samples, which was noticeably less than the ZIKV-Ug screen and perhaps due to the higher noise associated with the reduced levels of cell death in this second screen (Figure S6f, Data File 2). Of these genes, 70% were also genome-wide hits in the ZIKV-Ug screen (Data File 2), suggesting the mechanisms that mediate ZIKV neuropathogenesis are grossly similar between the two strains. It is possible that the differences in virulence that we observed are the result of the hundreds of passages undergone by ZIKV-Ug over the past 70 years that made it more adapted to culturing conditions compared to recently isolated strains, such as ZIKV-PR (Rossi et al., 2018). To facilitate rapid confirmation of the results of the primary survival screen for ZIKV host factors, we generated a H1 hESC line that constitutively expresses Cas9 (clone H1-36-23) and validated its genome-editing properties (Figure S6g-j, Data File 1). H1-Cas9 hESCs were then induced into SNaPs and transduced with lenti-sgRNAs in an arrayed format for cell viability assays at 120 hpi and infectivity assays at 54 hpi using ZIKV-Ug (MOI = 1). We targeted a subset of genes that were significantly enriched in both SNaP survival screens and represented the spectrum of ZIKV-relevant biological processes identified by our primary screens. As predicted by our primary screen, administering guides to SNaPs that target the EMC genes, *SLC35B2*, OST complex components, and vATPase subunits resulted in significantly improved cell viability compared to NT sgRNA controls in response to ZIKV-Ug infection, with several reaching >50% survival (Figure 6d). These results were corroborated by infectivity assays that showed dramatic reductions in 4G2^+^/PAX6^+^ SNaPs 54 hours after exposure to ZIKV-Ug (Figure 6e-f). In total, this work provides a list of hundreds of statistically-enriched known and candidate ZIKV host factors, several of which were validated in a secondary screen, in an *in vitro* model of human NPCs.

The transmembrane protein AXL, which is a member of the TYRO3-AXL-MERTK (TAM) receptor tyrosine kinase subfamily, was found to be a top ranking gene in a previously reported HeLa cell screen (Savidis et al., 2016a). This is in line with other reports suggesting a potential role for *AXL* in ZIKV cell attachment and/or entry in various cell types (Nowakowski et al., 2016). We previously tested this hypothesis in human NPCs and found that *AXL* was not essential for ZIKV infection (Wells et al., 2016), a finding corroborated by other groups in several model systems (Hastings et al., 2017; Meertens et al., 2017). To determine the effects of *AXL* and other TAMs in our primary screen, we converted all sgRNA log_2_ fold change values to z-scores. As expected, our most enriched genes showed relatively high positive z-scores in the ZIKV-Ug and ZIKV-PR screens (Figure S6k-l). The disruption of *AXL*, however, had no discernible effect on SNaP survival, as the mean z-score of *AXL* sgRNAs was -0.157 and -0.283 in the ZIKV-Ug and ZIKV-PR screens, respectively. Interestingly, the z-scores of TAM genes *TYRO3* and *MERTK*, as well as other candidate ZIKV receptors *CD209*, *HAVCR1*, and *TIMD4*, were also near zero. Furthermore, the CRISPR-Cas9 disruption of individual TAM family genes did not consistently promote SNaP survival at 120 hpi (Figure S6m). These data support the argument that contrary to other cell types, AXL and other TAM proteins do not serve as attachment factors for ZIKV infection in human NPCs.

### Whole genome CRISPR-Cas9 screen for SNaP fitness and proliferation genes

The regulation of NPC growth dynamics is critical to normal neurodevelopment, and defects in these mechanisms contribute to several types of brain disorders including cancer, ASDs, and intellectual disability (Piven et al., 2017). To identify SNaP fitness genes (defined as genes whose disruption results in a proliferation defect) and proliferation-enhancing genes, we compared the sequencing results of Day 10 mock infection samples to the Day 0 samples to ascertain which genes became enriched or depleted over the course of 10 days under normal growth conditions. We again observed high correlation among the three replicates (r values >0.92), and NT sgRNA controls had minimal effects (Figure S7a-c).

Our screen identified clear genetic contributors to SNaP proliferation, as RSA analysis detected 87 significantly-enriched genes that conferred enhanced cell survival or proliferation after CRISPR-mediated ablation (RSA_up_ < -4.0, adjusted p-value < 0.05; Figure 7a, Data File 3). This gene set consists of several known tumor suppressor genes as defined by the COSMIC cancer gene census (Forbes et al., 2017) including *NF2*, *SUFU*, *PTCH1*, and *TGFBR2*. Analysis of this gene set for GO biological processes yielded a statistical overrepresentation of terms relevant to primary cilia formation and maintenance, neural tube development, and regulation of epithelial cell proliferation (Figure 7b, Data File 3). Kyoto Encyclopedia of Genes and Genomes (KEGG) analysis revealed an enrichment of genes involved in multiple cancer mechanisms, as well as the sonic hedgehog (SHH) and WNT signal transduction pathways, both of which are intricately linked to cilia function (Guemez-Gamboa et al., 2014) (Figure S7d). In fact, of the 87 significantly-enriched gene hits, 42 are affiliated with the “cilium organization” (GO: 0044782) category, including *IFT57*, *IFT80*, *B9D1*, and the kinesin superfamily (KIF) proteins *KIFAP3*, *KIF3A*, and *KIF3B*. While previous *in vitro* and *in vivo* studies have uncovered the involvement of these molecular pathways in NPC proliferation and embryonal tumors of the central nervous system (Liu et al., 2017), several genes with no established role in proliferation were detected in our screen, including *SCRG1*, *KIAA1109*, and the putative voltage-gated calcium channel regulator *CACHD1*. These results, which demonstrate that known genetic drivers of NPC proliferation can be recapitulated in SNaPs, suggest that the novel proliferation genes identified in this primary screen should be considered for similar roles *in vivo*.

**Figure 7:**
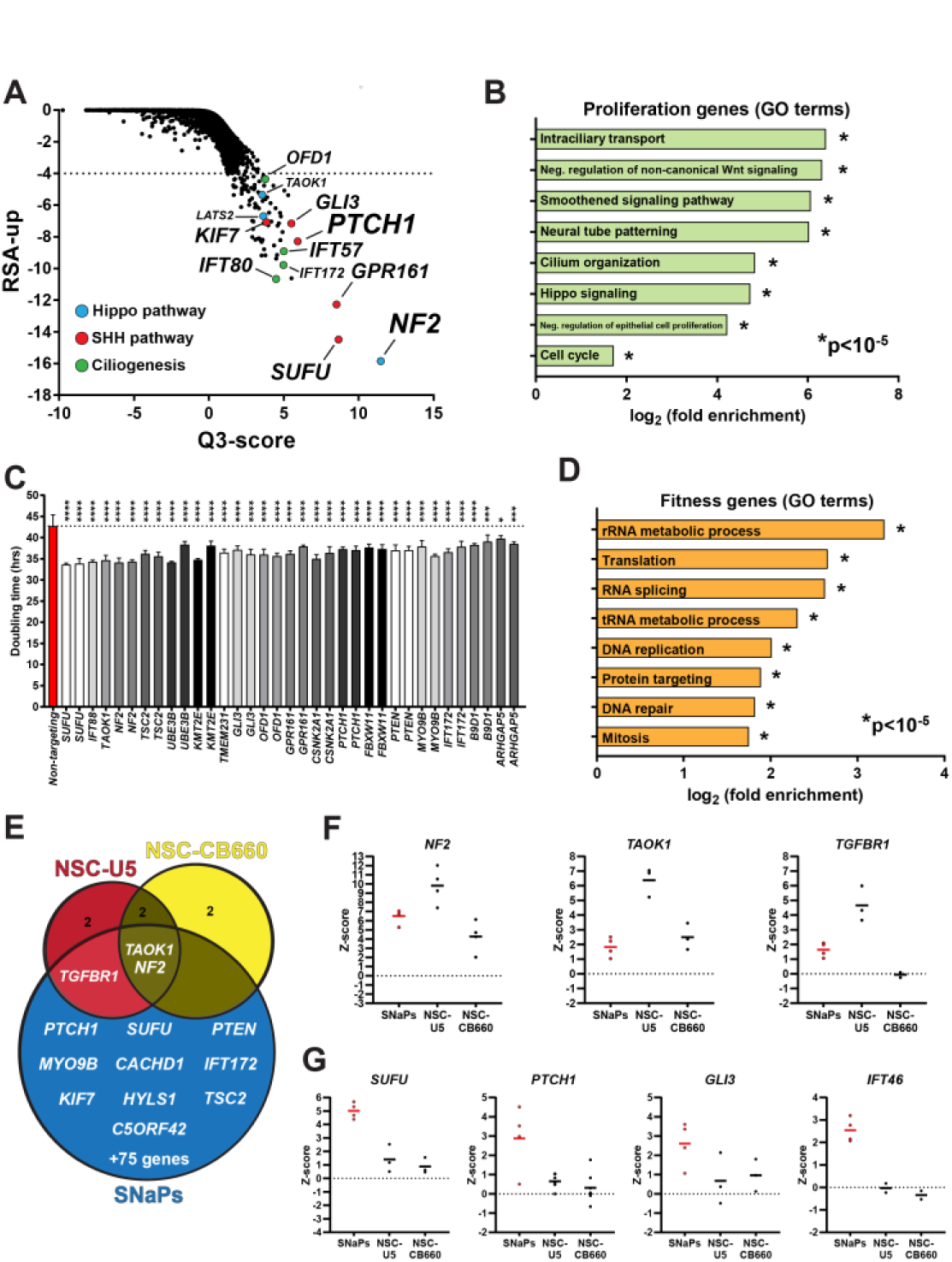
Fitness screen identifies SHH signaling and primary cilia genes as top contributors to enhanced SNaP proliferation. **A**, RSA analysis of genome-wide CRISPR-Cas9 SNaP fitness screen. Dashed line at RSA_up_ = -4.0 denotes hit cutoff for genes enriched at Day 10 relative to Day 0 samples. **B**, GO term analysis of the 87 RSA-identified significantly-enriched genes. **C**, Validation of top proliferation genes from primary fitness screen. Data presented as calculated doubling rates over 9 days of proliferation (n = 4 wells per sgRNA). **D**, GO term analysis of 1,449 fitness genes (Bayes Factor >10). E, Venn diagram showing overlap of MaGeCK-ranked genes in SNaP and primary fetal-derived neural stem cell (NSC-U5 and NSC-CB660) fitness screens. **F-G**, Z-scores of sgRNA log_2_ fold change for (F) overlapping hit genes and (G) top-scoring genes in the SNaP screen. Fisher’s exact test (B, D) and one-way ANOVA with Dunnett’s test for multiple comparisons (C) were used for statistical analysis. Data are represented as mean ± S.D. N.S. = not significant, *p<0.05, **p<0.01, ***p<0.001, ****p<0.0001.

To confirm the results of the primary fitness screen, we selected a subset of the positively-enriched genes and individually assessed the effects of their knockdown on SNaP proliferation. Using an arrayed screening format, we transduced Cas9-expressing SW.1 SNaPs with lentiviral sgRNAs that target our genes of interest. After 7 days of puromycin selection and SNaP expansion, cells were seeded onto 96-well plates at a low density (3,000 cells/cm^2^) and then imaged at Day 1 and Day 10 after plating using the cell-permeable Hoechst-33342 dye. As expected, the SNaPs containing sgRNAs directed against our genes of interest proliferated significantly faster than NT control sgRNAs (Figure 7c).

Previous genome-wide CRISPR screens have identified a set of core fitness genes whose disruption lead to decreases in viability regardless of cell type (Hart et al., 2014, 2015). Using Bayesian Analysis of Gene Essentiality (BAGEL) analysis (Hart and Moffat, 2016), in which more positive Bayes Factor (BF) scores indicate higher confidence that deletion of a specific gene results in decreased cell fitness, we identified 1,449 SNaP fitness genes (BF >10; Supplementary Data File 4). This gene set showed high concordance with the aforementioned set of core fitness genes (Figure S7e). In agreement with previous reports (Hart et al., 2015), our fitness gene set showed enrichment for critical cellular functions such as RNA processing (*DIMT1*, *HEATR1*, *NOL6*), protein translation (*RPS9, IMP3, EIF3M*), and DNA replication (*TIMELESS, PCNA, POLA2*; Figure 7d, Figure S7f, Data File 4). These results are consistent with the hypothesis that most mammalian cells share a common set of fitness genes required for survival that is independent of cell type (Hart et al., 2015).

To better assess the performance of our fitness screen relative to similar screens conducted in human NPCs, we compared our results to genome-wide CRISPR-Cas9 screens performed on two human fetal-derived neural stem/progenitor cell lines (NSC CB-660 and U5) (Toledo et al., 2015). We used Model-based Analysis of Genome-wide CRISPR/Cas9 Knockout (MAGeCK) analysis (Li et al., 2014) due to the ease at which multiple publicly-available read count datasets could be processed through the CRISPRAnalyzer pipeline (Winter et al., 2017). Similar to the RSA analysis, MaGeCK identified 88 significantly-enriched genes in the Day 10 SNaP samples, which included two genes, *TSC2* and *MYO9B*, that did not meet our initial RSA inclusion criteria but did reach RSA genome-wide significance (*TSC2*: RSA_up_ = -3.923, adjusted p-value = 0.0242; *MYO9B*: RSA_up_ = -3.580, adjusted p-value = 0.0401). On the contrary, only six and seven genes reached MaGeCK-defined significance in the CB660 and U5 primary NSC screens, respectively (Figure 7e, Data File 5). Within this limited set of genes, three were also found to significantly impact SNaP proliferation, including *TAOK1* (RSA_up_ = -5.370, adjusted p-value = 1.54e-3), *TGFBR1* (RSA_up_ = -4.727, adjusted p-value = 0.00491), and our top overall hit *NF2* (RSA_up_ = -15.851, adjusted p-value = 2.73e-12) (Figure 7f). The overlap of significantly-enriched genes in these datasets support the NPC classification of SNaPs by suggesting that they share mechanisms of proliferation, which have been shown to be cell-type specific (Sack et al., 2018), with NPCs derived from human fetal tissue.

Despite these similarities, our screen detected several biological pathways known to regulate NPC proliferation that were not identified by the fetal NSC screens, the most notable being SHH and ciliary genes. When we normalized the log_2_ fold change of each sgRNA within a screen using z-scores, many of our top SHH (*PTCH1*, *GLI3*, *GPR161*) and ciliary (*IFT46*, *IFT88*, *CEP120*) genes showed minimal enrichment, if not depletion, in the fetal NSC screens (Figure 7g, Figure S7g). The robustness of the list of genes that reached genome-wide significance in our screen, in conjunction with the agreement of our results with previous studies and clinical reports on NPC proliferation and cancer development, exemplifies the unique applicability of SNaP technology to large-scale genetic screens.

Finally, underscoring the importance of NPC proliferation in neurodevelopment, our list of genome-wide significant proliferation genes includes ASD candidate genes *PTEN* and *KMT2E*, as well as intellectual disability candidate genes *SOX11* and *CSNK2A1*. In addition, several genes identified by this screen as essential to SNaP viability have been linked to neurodevelopmental disorders, including *DDX3X* and *USP7*. These data suggest that SNaPs could be used to understand how mutations in critical neurodevelopment genes regulate NPC growth dynamics and ultimately lead to the associated brain disorders.

## DISCUSSION

While many seminal studies on the development of the mammalian brain have depended on *in vitro* and *in vivo* rodent models, the translation of these findings to human biology has been hampered by inherent differences in the genomic and architectural structure of the brains of these species. Here, we validated and demonstrated the effectiveness of a 48-hour induction protocol for stem cell-derived human NPCs that eliminates the need for labor-intensive steps and multi-week differentiation treatments. This method allowed us to proceed from cultures of hPSCs to completion of several whole-genome CRISPR-Cas9 screens in NPCs within 30 days.

Our SNaP protocol is reliant on the activities of NGN2, which is a powerful regulator of neuralization (Fode et al., 1998). Early work found that this transcription factor is highly expressed in the ventricular zone and cortical neuroepithelium of the developing mouse brain starting as early as embryonic day 8.5 when NPCs are abundant, and shows a noticeable downregulation as neuronal differentiation progresses (Sommer et al., 1996). NGN2 participates in cell specification and fate by promoting molecular pathways that produce dorsal glutamatergic neurons while inhibiting mechanisms that favor glial identities (Fode et al., 2000; Sun et al., 2001). Due to its established role in neuralization, exogenous expression of NGN2 has been used to generate human neurons from both somatic cells and hPSCs (Vierbuchen et al., 2010; Zhang et al., 2013). In this report, we present evidence to suggest that stem cell-derived neurons produced in this manner, like their *in vivo* counterparts, pass through a progenitor stage. This finding is supported by previously published qPCR data that hinted at this possibility (Zhang et al., 2013), as well as a more extensive RNA expression profiling carried out by our laboratory on patterned neurons (Nehme et al., 2018). Notably, this is in stark contrast to somatic cell-to-neuron transdifferentiation approaches that seem to bypass the proliferative progenitor cell stage of development (Son et al., 2011; Treutlein et al., 2016; Vierbuchen et al., 2010).

### SNaPs are an *in vitro* model of human NPCs

We provide several lines of evidence that the SNaP method produces cells with a human NPC identity. Transcript and protein expression analysis of SNaPs found molecular markers commonly expressed in *in vitro* and *in vivo* early neuroectoderm, such as *PAX6*, *NESTIN*, and *SOX2*. As revealed by immunocytochemistry and single-cell RNA sequencing experiments, SNaPs can give rise to neurons and glia, which is a basic function of NPCs. Importantly, SNaP-derived neurons display electrophysiological and immunocytochemical properties that are similar to NGN2/SMAD inhibition-induced neurons (Nehme et al., 2018), including network silencing in response to AMPA and NMDA receptor antagonism and the expression of glutamatergic cortical neuron protein markers. Clonal analyses verified the self-renewal properties of SNaPs, while three-dimensional SNaPsphere cultures were shown to exhibit behaviors previously observed in neurospheres derived from primary mouse and human tissue (Flax et al., 1998).

Two of the greatest strengths of the SNaP method are its speed and its reproducibility. Using the expression of three NPC protein markers and the absence of a pluripotent stem cell marker as our metric, we concluded that SNaP induction was successful in over 95% of the hPSC lines that we tested. More conventional neural differentiation schemes employing inhibition of SMAD signaling require specific cell densities to ensure successful forebrain NPC induction (Chambers et al., 2009), and this requirement could contribute to line-to-line and run-to-run induction efficiency variation (Engel et al., 2016; Muratore et al., 2014). SNaP induction was successful across a wide range of starting cell numbers (Figure S1e). While it is outside the scope of this investigation, this feature of the SNaP system could be beneficial to future comparative studies that rely on the efficient neuralization of large numbers of patient and control hiPS cell lines to identify disease-relevant molecular and cellular perturbations.

### SNaPs provide an *in vitro* model of ZIKV neuropathogenesis

The SNaP induction protocol allows for the rapid production of ZIKV-susceptible human NPCs that display characteristic antiviral and viability responses to infection, and as such could be used to study the mechanisms by which ZIKV infections lead to cell death in a microcephaly-relevant cell type. Since the recent and highly-publicized emergence of this virus as a public health risk, considerable efforts have been taken to better understand the cellular mechanisms of neuropathogenesis. Recent work has shown that heparan sulfate, which is expressed on the surface of most animal cells, acts as an attachment factor for ZIKV and other pathogens, including Chikungunya and the herpes simplex virus (Kim et al., 2017; Tanaka et al., 2017). In addition, it has been shown that ZIKV enters cells through clathrin-mediated endocytosis (CME) in a manner that involves EMC proteins (Savidis et al., 2016a). Furthermore, researchers have demonstrated that once in the endosome, ZIKV is able to continue the infection process with the help of vATPases, which are ATP-dependent proton pumps that help maintain endosomal pH and allow for viral release into lysosomes (Kozik et al., 2012). Our genome-wide CRISPR-Cas9 screens identified and validated these cellular processes as important players in the ZIKV-induced death of human NPCs, while also illuminating other genes and pathways not previously associated with flaviviruses.

*RACK1* and many O-linked glycosylation oligomeric Golgi complex (COG) genes went undetected in other flavivirus screens while reaching genome-wide significance in our SNaP screen (Krishnan et al., 2008; Ma et al., 2015; Qian et al., 2016; Savidis et al., 2016a; Zhang et al., 2016). The inhibition of *RACK1* was recently found to block internal ribosome entry site (IRES)-mediated viral translation without affecting cell viability (Majzoub et al., 2014). This same study showed that *RACK1* had no effect on cap-dependent translation, which is particularly interesting given that ZIKV is believed to utilize cap-dependent mechanisms (Göertz et al., 2018). There are two possible explanations for this phenomenon that could be addressed in future investigations: the kinase receptor *RACK1* could have an as-of-yet identified role in viral entry or replication; or *RACK1* does in fact play a role in cap-dependent translation. Regardless, our results indicate that the potentially broad antiviral effects of *RACK1* inhibition could extend to ZIKV in NPCs.

Previous studies have implicated COG mechanisms in the brain-infecting Rift Valley Fever and HIV-1 viruses (Brass et al., 2008; Riblett et al., 2016). Interestingly, our screen identified several other genes involved in the HIV-1 replication cycle, including *SLC35B2*, *MATR3*, *SND1*, *TNPO*, and *PAPSS1* (Park et al., 2017), as well as *ATP6V0C*, which is critical for replication of the NPC-infecting human cytomegalovirus (Pavelin et al., 2013). This suggests an interesting and potentially therapeutically-relevant scenario in which ZIKV infects neural cells and causes damage to the central nervous system through host mechanisms shared with viruses that show brain tropism but differ dramatically in structure, genome composition, and phylogenetic classification.

In addition to strengthening the argument that TAM receptors, such as *AXL* and *TYRO3*, play little to no role in ZIKV pathogenesis of human NPCs, our screen identified potential mechanisms of ZIKV attachment and entry into this cell type (e.g. heparan sulfate, vATPases). These processes, however, are not unique to ZIKV nor NPCs, and therefore do not fully explain the preferential targeting of NPCs over hPSCs and differentiated neurons. While it should be noted that it is entirely possible that there is no unique attachment or entry factor on the surface of human NPCs that could explain why this cell type is particularly vulnerable to ZIKV infection and death, the use of SNaPs could accelerate the search for these mechanisms by offering a quick and straightforward alternative to traditional NPC induction protocols.

### SHH and primary cilia govern SNaP proliferation

Regulation of NPC growth and viability has wide-ranging effects on neurodevelopment and cancers. Our genome-wide fitness screen produced a ranked list of genetic contributors to SNaP proliferation, including known tumor suppressor genes *TSC2*, which is a causative factor in tuberous sclerosis (The European Chromosome 16 Tuberous Sclerosis Consortium, 1993), and *PTEN*, which is a known inhibitor of the PI3K/AKT pathway and a commonly-mutated gene in brain tumors (Northcott et al., 2017). Pathway analysis of the top scoring genes pointed to two biological processes as strong governors of SNaP proliferation—SHH signal transduction and ciliogenesis.

SHH pathway activation is initiated by the SHH ligand binding to PTCH1 at cilia, which leads to glioma-associated oncogene (GLI) translocation to the nucleus, induction of target genes, and ultimately cell proliferation. Dysregulation of the SHH pathway is a leading contributor to the formation of the most common type of malignant brain tumor— medulloblastoma (MB)—and loss-of-function mutations in the SHH pathway repressors *PTCH1* and *SUFU* account for nearly 30% percent of all MB cases (Parsons et al., 2011). Therapeutics that inhibit SHH signaling, primarily in the form of Smoothened (SMO) antagonists, have been shown to be effective in both animal models and a subset of patients with MBs, though long-term efficacy has been hindered by the onset of drug resistance (Bautista et al., 2017; Rudin et al., 2009). Focused investigations of SMO inhibitor-resistant MBs found that it is the loss of primary cilia, driven by mutations in ciliary genes *IFT80*, *KIF3A*, and *OFD1,* that leads to persistent, low-level activation of the SHH pathway and abnormal cell proliferation (Goetz et al., 2009; Zhao et al., 2017). The preponderance of top scoring SHH and ciliary genes in our SNaP screen suggests that this model system could be used for future *in vitro* investigations of the cilia-driven mechanisms that lead to the onset of drug resistance, while also nominating a longer list of genes that could be involved in this process including *B9D2*, *IFT20*, and *TMEM17*. Ciliogenesis and SHH signaling were not the only mechanisms associated with cell proliferation enriched in our screen. Ablation of our top scoring gene neurofibromin-2 (*NF2*) underlies type II neurofibromatosis, which is characterized by the development of benign brain tumors that cause hearing loss and vestibular problems (Trofatter et al., 1993). *NF2* regulates cell proliferation through its activation of the Hippo pathway. This pathway mediates cell-cell contact inhibition and is activated by the phosphorylation of the large tumor suppressor kinase *LATS2*. *NF2* as well as *AMOT* and *TAOK1* regulate *LATS2* activities and in doing so inhibit the pro-proliferation transcription factors YAP and TAZ (Ahmed et al., 2017). The absence of these LATS regulators results in the inhibition of the Hippo pathway, which allows for the nuclear translocation of YAP and TAZ and unmitigated transcription of the target genes that lead to excessive proliferation.

To our knowledge, this is the first whole-genome CRISPR-Cas9 fitness screen conducted in hPSC-derived NPCs. As we hoped, many tumor suppressor genes associated with brain cancer reached genome-wide significance. Similar to our findings, a screen conducted in two fetal-derived neural stem cell lines that used the human GeCKO sgRNA library (64,751 unique sgRNAs targeting 18,080 genes) found that the *NF2* gene was the most enriched (Toledo et al., 2015). However, overall overlap with the SNaP screen was minimal, with *PTCH1*, *TSC2*, *GPR161*, and most of the IFT genes being noticeably absent from their list of top 500 sgRNAs. It is likely that the lack of congruence among these screens is the direct result of differences in experimental design. Though the sgRNA libraries used in these screens cover relatively similar genomic ground (18,000+ genes, 3.5+ sgRNAs per gene), the Brunello library has improved sgRNA designs that maximize on-target activity predictions and minimize off-target sites, and has shown a clear improvement over existing libraries including GeCKO (Doench et al., 2016). Given that previous reports have suggested that prolonged *in vitro* expansions of fetal NSCs can alter certain cellular phenotypes, such as proliferation rates, *in vivo* graft viability, and permissiveness to viral infection (Pan et al., 2013; Sun et al., 2011; Zietlow et al., 2012), it is also possible that cell culture history significantly influenced the fetal NSC screen results, as these cells were serially passaged many times after derivation prior to sgRNA delivery compared to the single passage for SNaPs. Overall, we believe that the 12-fold increase in significantly-enriched genes in our fitness screen compared to the NSC screens, as well as the strong parallels between our gene set and well-known drivers of NPC proliferation, illustrate the advantages of the Brunello-SNaP approach relative to previous screens in this cell type. More broadly, we conclude that the SNaP induction system detailed in this report is superior to accepted practices for producing NPCs in terms of efficiency, reproducibility, and applicability towards large-scale screens.

## Supporting information

## ACKNOWLEDGEMENTS

This work was supported by the Stanley Center for Psychiatric Research at the Broad Institute and the Harvard University Faculty of Arts and Sciences Dean’s Competitive Fund for Promising Scholarship, as well as NIH/NIMH grants U01MH105669 and U01MH115727. M.F.W. is supported by the Burroughs Wellcome Fund Postdoctoral Enrichment Program. We thank Janell Smith, Kavya Raghunathan (Harvard University), and Dr. Martin Berryer (Broad Institute) for their assistance in maintaining cell cultures. We acknowledge Olivia Bare, Adam Brown, and David Root (Broad Institute) for their contributions to the design and implementation of the CRISPR-Cas9 screens.

## AUTHOR CONTRIBUTIONS

M.F.W., E.J.H., and K.E. conceived the project. M.F.W., M.R.S., F.P., A.K., and K.E. designed and supervised experiments and interpreted results. M.F.W. and E.J.H. designed and optimized the SNaP induction protocol. M.F.W. performed all SNaP validation experiments. M.R.S. and J.J.R. performed and analyzed scRNAseq experiments. M.F.W. and M.T.S. conducted SNaPsphere assays. M.F.W. and F.P. designed and analyzed the CRISPR-Cas9 screens. D.H. and B.K.P. performed the RSA analysis. S.K., K.C., J. J.R., and K.A.W. generated the H1-Cas9 cell line used for validation of the ZIKV survival screen. R.N. and O.P. generated the time course RNAseq data. M.F.W. and K.E. wrote the paper, and all authors provided feedback.

## DECLARATION OF INTERESTS

The authors declare no competing interests.

## REFERENCES

Ahmed, A.A., Mohamed, A.D., Gener, M., Li, W., and Taboada, E. (2017). YAP and the Hippo pathway in pediatric cancer. Mol. Cell. Oncol. 4, e1295127.

Alvarez-Buylla, A., Theelen, M., and Nottebohm, F. (1990). Proliferation “hot spots” in adult avian ventricular zone reveal radial cell division. Neuron 5, 101–109.

Bautista, F., Fioravantti, V., de Rojas, T., Carceller, F., Madero, L., Lassaletta, A., and Moreno, L. (2017). Medulloblastoma in children and adolescents: a systematic review of contemporary phase I and II clinical trials and biology update. Cancer Med. 6, 2606–2624.

Brasil, P., Pereira, J.P., Moreira, M.E., Ribeiro Nogueira, R.M., Damasceno, L., Wakimoto, M., Rabello, R.S., Valderramos, S.G., Halai, U.-A., Salles, T.S., et al. (2016). Zika Virus Infection in Pregnant Women in Rio de Janeiro. N. Engl. J. Med. 375, 2321–2334.

Brass, A.L., Dykxhoorn, D.M., Benita, Y., Yan, N., Engelman, A., Xavier, R.J., Lieberman, J., and Elledge, S.J. (2008). Identification of Host Proteins Required for HIV Infection Through a Functional Genomic Screen. Science (80-.). 319, 921 LP-926.

Brennand, K., Savas, J.N., Kim, Y., Tran, N., Simone, A., Hashimoto-Torii, K., Beaumont, K.G., Kim, H.J., Topol, A., Ladran, I., et al. (2015). Phenotypic differences in hiPSC NPCs derived from patients with schizophrenia. Mol. Psychiatry 20, 361–368.

Chambers, S.M., Fasano, C.A., Papapetrou, E.P., Tomishima, M., Sadelain, M., and Studer, L. (2009). Highly efficient neural conversion of human ES and iPS cells by dual inhibition of SMAD signaling. Nat. Biotechnol. 27, 275–280.

Doench, J.G., Fusi, N., Sullender, M., Hegde, M., Vaimberg, E.W., Donovan, K.F., Smith, I., Tothova, Z., Wilen, C., Orchard, R., et al. (2016). Optimized sgRNA design to maximize activity and minimize off-target effects of CRISPR-Cas9. Nat. Biotechnol. 34, 184–191.

Elkabetz, Y., Panagiotakos, G., Al Shamy, G., Socci, N.D., Tabar, V., and Studer, L. (2008). Human ES cell-derived neural rosettes reveal a functionally distinct early neural stem cell stage. Genes Dev. 22, 152–165.

Engel, M., Do-Ha, D., Muñoz, S.S., and Ooi, L. (2016). Common pitfalls of stem cell differentiation: a guide to improving protocols for neurodegenerative disease models and research. Cell. Mol. Life Sci. 73, 3693–3709.

Flax, J.D., Aurora, S., Yang, C., Simonin, C., Wills, A.M., Billinghurst, L.L., Jendoubi, M., Sidman, R.L., Wolfe, J.H., Kim, S.U., et al. (1998). Engraftable human neural stem cells respond to development cues, replace neurons, and express foreign genes. Nat. Biotechnol. 16, 1033.

Fode, C., Gradwohl, G., Morin, X., Dierich, A., LeMeur, M., Goridis, C., and Guillemot, F. (1998). The bHLH protein NEUROGENIN 2 is a determination factor for epibranchial placode-derived sensory neurons. Neuron 20, 483–494.

Fode, C., Ma, Q., Casarosa, S., Ang, S.L., Anderson, D.J., and Guillemot, F. (2000). A role for neural determination genes in specifying the dorsoventral identity of telencephalic neurons. Genes Dev. 14, 67–80.

Forbes, S.A., Beare, D., Boutselakis, H., Bamford, S., Bindal, N., Tate, J., Cole, C.G., Ward, S., Dawson, E., Ponting, L., et al. (2017). COSMIC: Somatic cancer genetics at high-resolution. Nucleic Acids Res. 45, D777–D783.

Gabriel, E., Ramani, A., Karow, U., Gottardo, M., Natarajan, K., Gooi, L.M., Goranci-Buzhala, G., Krut, O., Peters, F., Nikolic, M., et al. (2017). Recent Zika Virus Isolates Induce Premature Differentiation of Neural Progenitors in Human Brain Organoids. Cell Stem Cell 20, 397–406.e5.

Garcez, P.P., Loiola, E.C., Da Costa, R.M., Higa, L.M., Trindade, P., Delvecchio, R., Nascimento, J.M., Brindeiro, R., Tanuri, A., and Rehen, S.K. (2016). Zika virus: Zika virus impairs growth in human neurospheres and brain organoids. Science (80-.). 352, 816–818.

Göertz, G.P., Abbo, S.R., Fros, J.J., and Pijlman, G.P. (2018). Functional RNA during Zika virus infection. Virus Res. 254, 41–53.

Goetz, S.C., Ocbina, P.J.R., and Anderson, K. V. (2009). The primary cilium as a Hedgehog signal transduction machine. (Elsevier).

Guemez-Gamboa, A., Coufal, N.G., and Gleeson, J.G. (2014). Primary Cilia in the Developing and Mature Brain. Neuron 82, 511–521.

Hamel, R., Dejarnac, O., Wichit, S., Ekchariyawat, P., Neyret, A., Luplertlop, N., Perera-Lecoin, M., Surasombatpattana, P., Talignani, L., Thomas, F., et al. (2015). Biology of Zika Virus Infection in Human Skin Cells. J. Virol. 89, 8880–8896.

Hart, T., and Moffat, J. (2016). BAGEL: A computational framework for identifying essential genes from pooled library screens. BMC Bioinformatics 17, 1–7.

Hart, T., Brown, K.R., Sircoulomb, F., Rottapel, R., and Moffat, J. (2014). Measuring error rates in genomic perturbation screens: gold standards for human functional genomics. Mol. Syst. Biol. 10, 733.

Hart, T., Chandrashekhar, M., Aregger, M., Steinhart, Z., Brown, K.R., MacLeod, G., Mis, M., Zimmermann, M., Fradet-Turcotte, A., Sun, S., et al. (2015). High-Resolution CRISPR Screens Reveal Fitness Genes and Genotype-Specific Cancer Liabilities. Cell 163, 1515–1526.

Hastings, A.K., Yockey, L.J., Jagger, B.W., Hwang, J., Uraki, R., Gaitsch, H.F., Parnell, L.A., Cao, B., Mysorekar, I.U., Rothlin, C. V., et al. (2017). TAM Receptors Are Not Required for Zika Virus Infection in Mice. Cell Rep. 19, 558–568.

Hemmati, H.D., Nakano, I., Lazareff, J.A., Masterman-Smith, M., Geschwind, D.H., Bronner-Fraser, M., and Kornblum, H.I. (2003). Cancerous stem cells can arise from pediatric brain tumors. Proc. Natl. Acad. Sci. 100, 15178–15183.

Hemmati-Brivanlou, A., and Meltont, D. (1997). Vertebrate embryonic cells will become nerve cells unless told otherwise. Cell 88, 13–17.

Homem, C.C.F., Repic, M., and Knoblich, J.A. (2015). Proliferation control in neural stem and progenitor cells. Nat. Rev. Neurosci. 16, 647–659.

Kelava, I., and Lancaster, M.A. (2016). Stem Cell Models of Human Brain Development. Cell Stem Cell 18, 736–748.

Kim, S.Y., Zhao, J., Liu, X., Fraser, K., Lin, L., Zhang, X., Zhang, F., Dordick, J.S., and Linhardt, R.J. (2017). Interaction of Zika Virus Envelope Protein with Glycosaminoglycans. Biochemistry 56, 1151–1162.

Koch, P., Opitz, T., Steinbeck, J.A., Ladewig, J., and Brustle, O. (2009). A rosette-type, self-renewing human ES cell-derived neural stem cell with potential for in vitro instruction and synaptic integration. Proc. Natl. Acad. Sci. 106, 3225–3230.

König, R., Chiang, C.Y., Tu, B.P., Yan, S.F., DeJesus, P.D., Romero, A., Bergauer, T., Orth, A., Krueger, U., Zhou, Y., et al. (2007). A probability-based approach for the analysis of large-scale RNAi screens. Nat. Methods 4, 847–849.

Kozik, P., Hodson, N.A., Sahlender, D.A., Simecek, N., Soromani, C., Wu, J., Collinson, L.M., and Robinson, M.S. (2012). A human genome-wide screen for regulators of clathrin-coated vesicle formation reveals an unexpected role for the V-ATPase. Nat. Cell Biol. 15, 50.

Kriegstein, A., and Alvarez-Buylla, A. (2009). The Glial Nature of Embryonic and Adult Neural Stem Cells. Annu. Rev. Neurosci. 32, 149–184.

Krishnan, M.N., Ng, A., Sukumaran, B., Gilfoy, F.D., Uchil, P.D., Sultana, H., Brass, A.L., Adametz, R., Tsui, M., Qian, F., et al. (2008). RNA interference screen for human genes associated with West Nile virus infection. Nature 455, 242.

Li, W., Xu, H., Xiao, T., Cong, L., Love, M.I., Zhang, F., Irizarry, R.A., Liu, J.S., Brown, M., and Liu, X.S. (2014). MAGeCK enables robust identification of essential genes from genome-scale CRISPR/Cas9 knockout screens. Genome Biol. 15, 554.

Liu, K.-W., Pajtler, K.W., Worst, B.C., Pfister, S.M., and Wechsler-Reya, R.J. (2017). Molecular mechanisms and therapeutic targets in pediatric brain tumors. Sci. Signal. 10, eaaf7593.

Lui, J.H., Hansen, D. V., and Kriegstein, A.R. (2011). Development and evolution of the human neocortex. Cell 146, 18–36.

Ma, H., Dang, Y., Wu, Y., Jia, G., Anaya, E., Zhang, J., Abraham, S., Choi, J.G., Shi, G., Qi, L., et al. (2015). A CRISPR-based screen identifies genes essential for west-nile-virus-induced cell death. Cell Rep. 12, 673–683.

Majzoub, K., Hafirassou, M.L., Meignin, C., Goto, A., Marzi, S., Fedorova, A., Verdier, Y., Vinh, J., Hoffmann, J.A., Martin, F., et al. (2014). RACK1 controls IRES-mediated translation of viruses. Cell 159, 1086–1095.

Marchetto, M.C., Belinson, H., Tian, Y., Freitas, B.C., Fu, C., Vadodaria, K.C., Beltrao-Braga, P.C., Trujillo, C.A., Mendes, A.P.D., Padmanabhan, K., et al. (2017). Altered proliferation and networks in neural cells derived from idiopathic autistic individuals. Mol. Psychiatry 22, 820–835.

Meertens, L., Labeau, A., Dejarnac, O., Cipriani, S., Sinigaglia, L., Bonnet-Madin, L., Le Charpentier, T., Hafirassou, M.L., Zamborlini, A., Cao-Lormeau, V.M., et al. (2017). Axl Mediates ZIKA Virus Entry in Human Glial Cells and Modulates Innate Immune Responses. Cell Rep. 18, 324–333.

Merkle, F.T., and Eggan, K. (2013). Modeling human disease with pluripotent stem cells: From genome association to function. Cell Stem Cell 12, 656–668.

Mesci, P., Macia, A., Moore, S.M., Shiryaev, S.A., Pinto, A., Huang, C.-T., Tejwani, L., Fernandes, I.R., Suarez, N.A., Kolar, M.J., et al. (2018). Blocking Zika virus vertical transmission. Sci. Rep. 8, 1218.

Mlakar, J., Korva, M., Tul, N., Popović, M., Poljšak-Prijatelj, M., Mraz, J., Kolenc, M., Resman Rus, K., Vesnaver Vipotnik, T., Fabjan Vodušek, V., et al. (2016). Zika Virus Associated with Microcephaly. N. Engl. J. Med. 374, 951–958.

Muratore, C.R., Srikanth, P., Callahan, D.G., and Young-Pearse, T.L. (2014). Comparison and optimization of hiPSC forebrain cortical differentiation protocols. PLoS One 9.

Nehme, R., Zuccaro, E., Ghosh, S.D., Li, C., Sherwood, J., Pietilainen, O., Barrett, L.E., Limone, F., Worringer, K.A., Kommineni, S., et al. (2018). Combining NGN2 programming with developmental patterning generates human excitatory neurons with NMDAR-mediated synaptic transmission. Cell Rep. 23, 2509–2523.

Noctor, S.C., Martinez-Cerdeño, V., Ivic, L., and Kriegstein, A.R. (2004). Cortical neurons arise in symmetric and asymmetric division zones and migrate through specific phases. Nat. Neurosci. 7, 136–144.

Northcott, P.A., Buchhalter, I., Morrissy, A.S., Hovestadt, V., Weischenfeldt, J., Ehrenberger, T., Gröbner, S., Segura-Wang, M., Zichner, T., Rudneva, V.A., et al. (2017). The whole-genome landscape of medulloblastoma subtypes. Nature 547, 311–317.

Nowakowski, T.J., Pollen, A.A., Di Lullo, E., Sandoval-Espinosa, C., Bershteyn, M., and Kriegstein, A.R. (2016). Expression analysis highlights AXL as a candidate zika virus entry receptor in neural stem cells. Cell Stem Cell 18, 591–596.

Pan, X., Li, X.-J., Liu, X.-J., Yuan, H., Li, J.-F., Duan, Y.-L., Ye, H.-Q., Fu, Y.-R., Qiao, G.-H., Wu, C.-C., et al. (2013). Later Passages of Neural Progenitor Cells from Neonatal Brain Are More Permissive for Human Cytomegalovirus Infection. J. Virol. 87, 10968–10979.

Park, R.J., Wang, T., Koundakjian, D., Hultquist, J.F., Lamothe-Molina, P., Monel, B., Schumann, K., Yu, H., Krupzcak, K.M., Garcia-Beltran, W., et al. (2017). A genome-wide CRISPR screen identifies a restricted set of HIV host dependency factors. Nat. Genet. 49, 193–203.

Parsons, D.W., Li, M., Zhang, X., Jones, S., Leary, R.J., Lin, J.C.-H., Boca, S.M., Carter, H., Samayoa, J., Bettegowda, C., et al. (2011). The genetic landscape of the childhood cancer medulloblastoma. Science (80-.). 331, 435–439.

Pavelin, J., Reynolds, N., Chiweshe, S., Wu, G., Tiribassi, R., and Grey, F. (2013). Systematic MicroRNA Analysis Identifies ATP6V0C as an Essential Host Factor for Human Cytomegalovirus Replication. PLoS Pathog. 9, 1–13.

Piven, J., Elison, J.T., and Zylka, M.J. (2017). Toward a conceptual framework for early brain and behavior development in autism. Mol. Psychiatry 22, 1385–1394.

Qian, X., Nguyen, H.N., Song, M.M., Hadiono, C., Ogden, S.C., Hammack, C., Yao, B., Hamersky, G.R., Jacob, F., Zhong, C., et al. (2016). Brain-Region-Specific Organoids Using Mini-bioreactors for Modeling ZIKV Exposure. Cell 165, 1238–1254.

Rakic, P. (2009). Evolution of the neocortex: A perspective from developmental biology. Nat. Rev. Neurosci. 10, 724–735.

Reynolds, B.A., and Weiss, S. (1992). Generation of neurons and astrocytes from isolated cells of the adult mammalian central nervous system. Science (80-.). 255, 1707 LP-1710.

Riblett, A.M., Blomen, V.A., Jae, L.T., Altamura, L.A., Doms, R.W., Brummelkamp, T.R., and Wojcechowskyj, J.A. (2016). A Haploid Genetic Screen Identifies Heparan Sulfate Proteoglycans Supporting Rift Valley Fever Virus Infection. J. Virol. 90, 1414–1423.

Rice, M.E., Galand, R.R., Roth, N.M., Ellington, S.R., Moore, C.A., Prado, M.V., Ellis, E.M., Tufa, A.J., Taulung, L.A., Alfred, J.M., et al. (2018). Vital Signs: Zika-Associated Birth Defects and Neurodevelopmental Abnormalities Possibly Associated with Congenital Zika Virus Infection — U.S. Territories and Freely Associated States, 2018. Morb. Mortal. Wkly. Rep. 67, 858–867.

Rossi, S.L., Ebel, G.D., Shan, C., Shi, P.Y., and Vasilakis, N. (2018). Did Zika Virus Mutate to Cause Severe Outbreaks? Trends Microbiol. 26, 877–885.

Rudin, C.M., Hann, C.L., Laterra, J., Yauch, R.L., Callahan, C.A., Fu, L., Holcomb, T., Stinson, J., Gould, S.E., Coleman, B., et al. (2009). Treatment of Medulloblastoma with Hedgehog Pathway Inhibitor GDC-0449. N. Engl. J. Med. 361, 1173–1178.

Sack, L.M., Davoli, T., Li, M.Z., Li, Y., Xu, Q., Naxerova, K., Wooten, E.C., Bernardi, R.J., Martin, T.D., Chen, T., et al. (2018). Profound Tissue Specificity in Proliferation Control Underlies Cancer Drivers and Aneuploidy Patterns. Cell 173, 499–514.e23.

Salick, M.R., Wells, M.F., Eggan, K., and Kaykas, A. (2017). Modelling Zika Virus Infection of the Developing Human Brain In Vitro Using Stem Cell Derived Cerebral Organoids. J. Vis. Exp. 1–10.

Sandoe, J., and Eggan, K. (2013). Opportunities and challenges of pluripotent stem cell neurodegenerative disease models. Nat. Neurosci. 16, 780–789.

Savidis, G., McDougall, W.M., Meraner, P., Perreira, J.M., Portmann, J.M., Trincucci, G., John, S.P., Aker, A.M., Renzette, N., Robbins, D.R., et al. (2016a). Identification of Zika Virus and Dengue Virus Dependency Factors using Functional Genomics. Cell Rep. 16, 232–246.

Savidis, G., Perreira, J.M., Portmann, J.M., Meraner, P., Guo, Z., Green, S., and Brass, A.L. (2016b). The IFITMs Inhibit Zika Virus Replication. Cell Rep. 15, 2323–2330.

Saxena, S.K., Kumar, S., Sharma, R., Maurya, V.K., Dandu, H.R., and Bhatt, M.L. (2018). Zika virus disease in India - Update October 2018. Travel Med. Infect. Dis.

Simonin, Y., Loustalot, F., Desmetz, C., Foulongne, V., Constant, O., Fournier-Wirth, C., Leon, F., Molès, J.P., Goubaud, A., Lemaitre, J.M., et al. (2016). Zika Virus Strains Potentially Display Different Infectious Profiles in Human Neural Cells. EBioMedicine 12, 161–169.

Simonin, Y., van Riel, D., Van de Perre, P., Rockx, B., and Salinas, S. (2017). Differential virulence between Asian and African lineages of Zika virus. PLoS Negl. Trop. Dis. 11, e0005821.

Smith, J.R., Vallier, L., Lupo, G., Alexander, M., Harris, W.A., and Pedersen, R.A. (2008). Inhibition of Activin/Nodal signaling promotes specification of human embryonic stem cells into neuroectoderm. Dev. Biol. 313, 107–117.

Sommer, L., Ma, Q., and Anderson, D.J. (1996). Neurogenins, a Novel Family of atonal-Related bHLH Transcription Factors, are Putative Mammalian Neuronal Determination Genes That Reveal Progenitor Cell Heterogeneity in the Developing CNS and PNS. 241, 221–241.

Son, E.Y., Ichida, J.K., Wainger, B.J., Toma, J.S., Rafuse, V.F., Woolf, C.J., and Eggan, K. (2011). Conversion of mouse and human fibroblasts into functional spinal motor neurons. Cell Stem Cell 9, 205–218.

Sun, T., Wang, X.-J., Xie, S.-S., Zhang, D.-L., Wang, X.-P., Li, B.-Q., Ma, W., and Xin, H. (2011). A comparison of proliferative capacity and passaging potential between neural stem and progenitor cells in adherent and neurosphere cultures. Int. J. Dev. Neurosci. 29, 723–731.

Sun, Y., Nadal-Vicens, M., Misono, S., Lin, M.Z., Zubiaga, A., Hua, X., Fan, G., and Greenberg, M.E. (2001). Neurogenin promotes neurogenesis and inhibits glial differentiation by independent mechanisms. Cell 104, 365–376.

Takahashi, K., Tanabe, K., Ohnuki, M., Narita, M., Ichisaka, T., Tomoda, K., and Yamanaka, S. (2007). Induction of Pluripotent Stem Cells from Adult Human Fibroblasts by Defined Factors. Cell 131, 861–872.

Tanaka, A., Tumkosit, U., Nakamura, S., Motooka, D., Kishishita, N., Priengprom, T., Sa-ngasang, A., Kinoshita, T., Takeda, N., and Maeda, Y. (2017). Genome-Wide Screening Uncovers the Significance of N-Sulfation of Heparan Sulfate as a Host Cell Factor for Chikungunya Virus Infection. J. Virol. 91, e00432-17.

Tang, H., Hammack, C., Ogden, S.C., and Jin, P. (2016). Zika Virus Infects Human Cortical Neural Progenitors and Attenuates Their Growth. Stem Cell 1–4.

TCW, J., Wang, M., Pimenova, A.A., Bowles, K.R., Hartley, B.J., Lacin, E., Machlovi, S.I., Abdelaal, R., Karch, C.M., Phatnani, H., et al. (2017). An Efficient Platform for Astrocyte Differentiation from Human Induced Pluripotent Stem Cells. Stem Cell Reports 9, 600–614.

The European Chromosome 16 Tuberous Sclerosis Consortium (1993). Identification and characterization of the tuberous sclerosis gene on chromosome 16. Cell 75, 1305–1315.

Thomson, J.A. (1998). Embryonic Stem Cell Lines Derived from Human Blastocysts. Science (80-.). 282, 1145–1147.

Toledo, C.M., Ding, Y., Hoellerbauer, P., Davis, R.J., Basom, R., Girard, E.J., Lee, E., Corrin, P., Hart, T., Bolouri, H., et al. (2015). Genome-wide CRISPR-Cas9 Screens Reveal Loss of Redundancy between PKMYT1 and WEE1 in Glioblastoma Stem-like Cells. Cell Rep. 13, 2425–2439.

Treutlein, B., Lee, Q.Y., Camp, J.G., Mall, M., Koh, W., Shariati, S.A.M., Sim, S., Neff, N.F., Skotheim, J.M., Wernig, M., et al. (2016). Dissecting direct reprogramming from fibroblast to neuron using single-cell RNA-seq. Nature 534, 391–395.

Trofatter, J.A., MacCollin, M.M., Rutter, J.L., Murrell, J.R., Duyao, M.P., Parry, D.M., Eldridge, R., Kley, N., Menon, A.G., Pulaski, K., et al. (1993). A novel moesin-, ezrin-, radixin-like gene is a candidate for the neurofibromatosis 2 tumor suppressor. Cell 72, 791–800.

Vierbuchen, T., Ostermeier, A., Pang, Z.P., Kokubu, Y., Südhof, T.C., and Wernig, M. (2010). Direct conversion of fibroblasts to functional neurons by defined factors. Nature 463, 1035.

Wells, M.F., Salick, M.R., Wiskow, O., Ho, D.J., Worringer, K.A., Ihry, R.J., Kommineni, S., Bilican, B., Klim, J.R., Hill, E.J., et al. (2016). Genetic Ablation of AXL Does Not Protect Human Neural Progenitor Cells and Cerebral Organoids from Zika Virus Infection. Cell Stem Cell 19, 703–708.

Winter, J., Schwering, M., Pelz, O., Rauscher, B., Zhan, T., Heigwer, F., and Boutros, M. (2017). CRISPRAnalyzeR: Interactive analysis, annotation and documentation of pooled CRISPR screens. bioRxiv.

Yi, F., Danko, T., Botelho, S.C., Patzke, C., Pak, C., Wernig, M., and Südhof, T.C. (2016). Autism-associated SHANK3 haploinsufficiency causes Ih channelopathy in human neurons. Science (80-.). 352.

Zhang, R., Miner, J.J., Gorman, M.J., Rausch, K., Ramage, H., White, J.P., Zuiani, A., Zhang, P., Fernandez, E., Zhang, Q., et al. (2016). A CRISPR screen defines a signal peptide processing pathway required by flaviviruses. Nature 535, 164–168.

Zhang, S.-C., Wernig, M., Duncan, I.D., Brüstle, O., and Thomson, J.A. (2001). In vitro differentiation of transplantable neural precursors from human embryonic stem cells. Nat. Biotechnol. 19, 1129–1133.

Zhang, Y., Pak, C.H., Han, Y., Ahlenius, H., Zhang, Z., Chanda, S., Marro, S., Patzke, C., Acuna, C., Covy, J., et al. (2013). Rapid single-step induction of functional neurons from human pluripotent stem cells. Neuron 78, 785–798.

Zhao, X., Pak, E., Ornell, K.J., Murphy, M.F.P., Mackenzie, E.L., Chadwick, E.J., Ponomaryov, T., Kelleher, J.F., and Segal, R.A. (2017). A transposon screen identifies loss of primary cilia as a mechanism of resistance to SMO inhibitors. Cancer Discov. 7, 1436–1439.

Zietlow, R., Precious, S. V, Kelly, C.M., Dunnett, S.B., and Rosser, A.E. (2012). Long-term expansion of human foetal neural progenitors leads to reduced graft viability in the neonatal rat brain. Exp. Neurol. 235, 563–573.

