## Supplementary material for "Genome-wide screens in accelerated human stem cell-derived neural progenitor cells identify Zika virus host factors and drivers of proliferation"

### Supplemental Figure 1

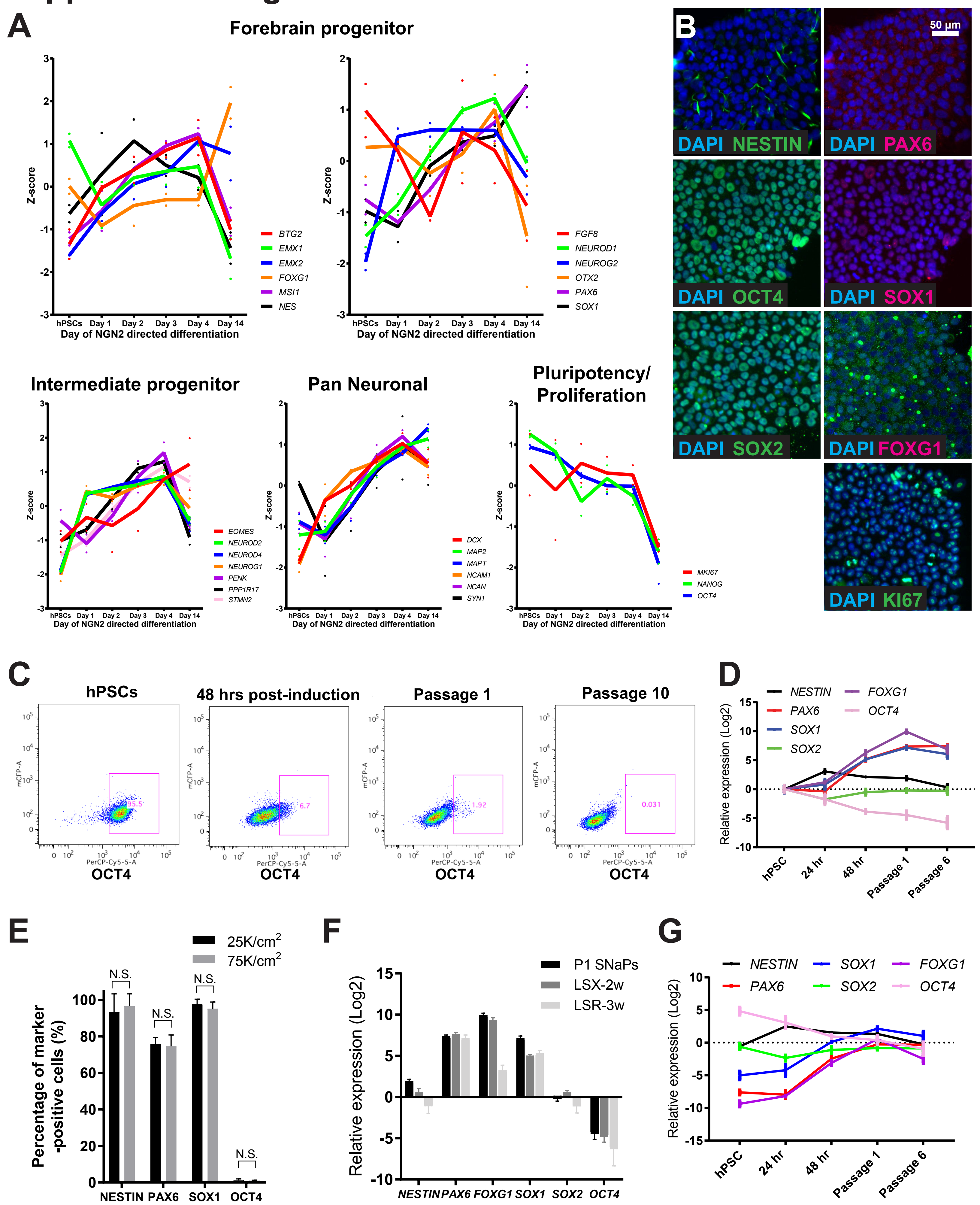

**Supplemental Figure 1 (Related to Figure 1): Developmental expression patterns of human pluripotent stem cell-derived neurons and SNaPs.** **A**, Normalized expression of forebrain progenitor, intermediate progenitor, neuronal, and pluripotency transcript markers over the course of NGN2-directed differentiation of neurons from SW.1 and SW.11 hiPSC starting materials (from *Nehme et al., 2018*). **B**, SW.1 hiPS cells show high levels of pluripotent stem cell markers OCT4 and SOX2, as well as proliferation marker KI67. NPC markers NESTIN, PAX6, SOX1, and FOXG1 are weakly expressed. **C**, FACS plots at different stages of the SNaP induction protocol. The high percentage of OCT4-positive cells (x-axis; PerCP-Cy5.5) decreases dramatically over the course of the SNaP induction protocol. **D**, qPCR analysis of SNaP cultures harvested at different stages of the induction protocol reveals upregulation of multiple NPC markers and downregulation of the OCT4 pluripotent stem cell marker compared to hPSC starting materials (n = 4). **E**, Comparison of SNaP protocol efficiency at 48 hours post-induction between 25k/cm$^2$ and 75K/cm$^2$ hPSC starting densities (n = 4 wells per condition). **F**, Comparison of P1 SNaP transcript levels to standard 2-week NPC induction protocol (LDN, SB431542, and XAV939; LSX-2w) and a modified 3-week method (LDN, SB431542, and Retinoic acid; LSR-3w). **G**, Same dataset in Panel D normalized to LSX-2w samples shows convergence of expression profiles. Data are represented as mean ± S.D. N.S. = not significant.

### Supplemental Figure 2

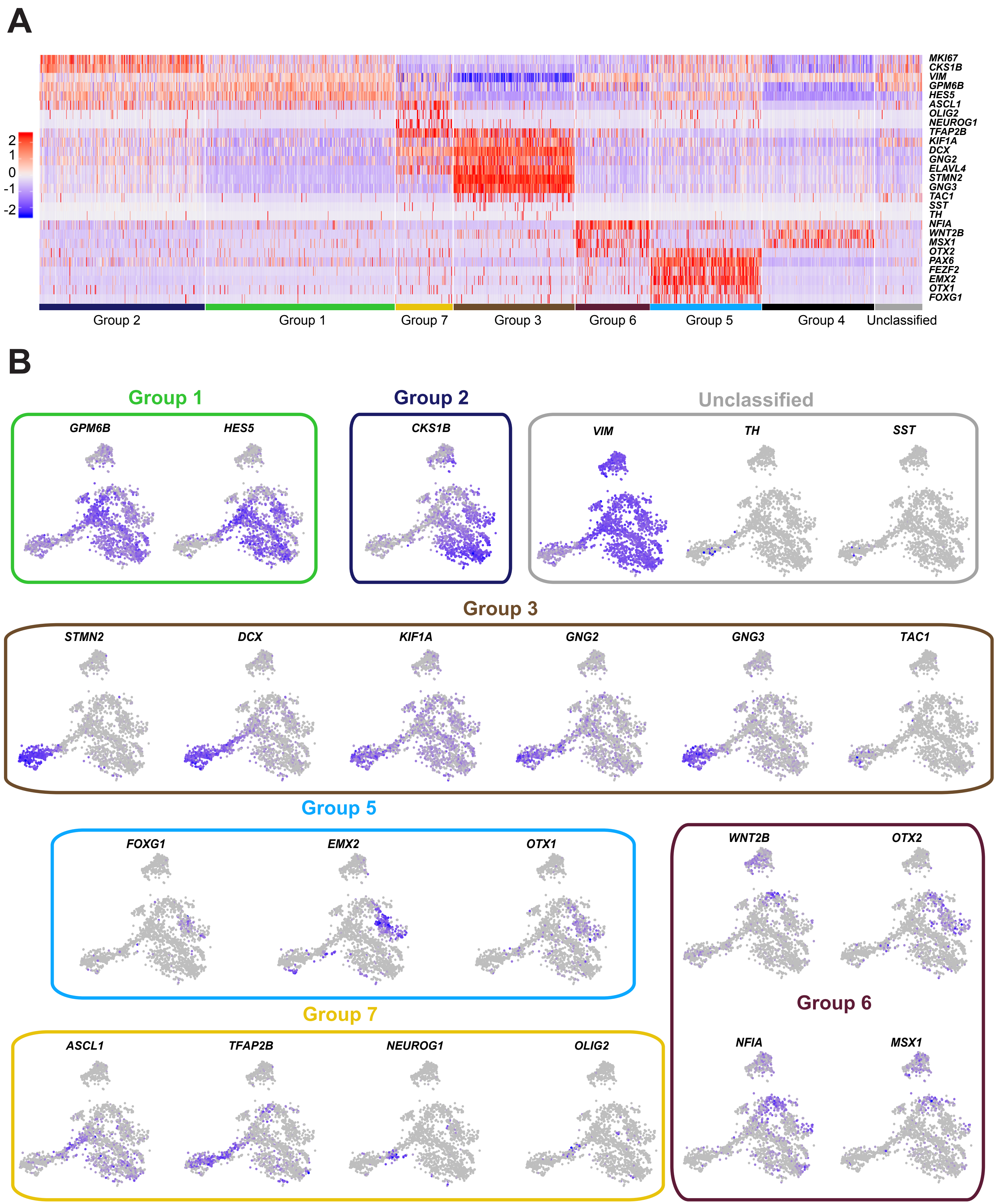

**Supplemental Figure 2 (Related to Figure 2): Single-cell characterization of SNaP spontaneous differentiation.** **A**, Heat map showing single-cell expression of cell type-marker transcripts after two weeks of spontaneous differentiation of SNaP cells in base media. **B**, TSNE plots of representative genes for each group.

### Supplemental Figure 3

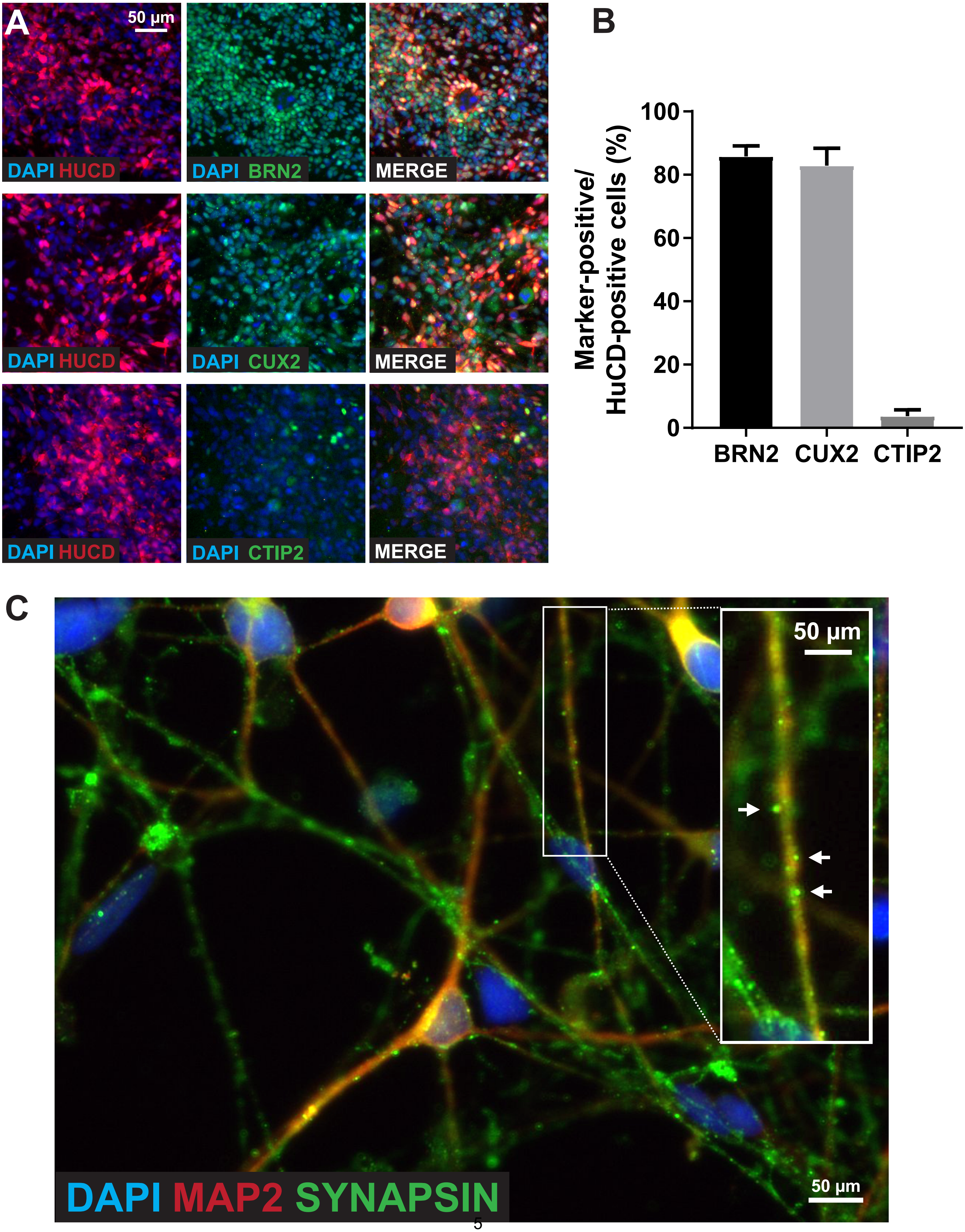

**Supplemental Figure 3 (Related to Figure 3): Immunocytochemical characterization of SNaP-derived neurons.** **A**, Co-localization of pan-neuronal marker HuC/D (red) and superficial cortical layer markers BRN2 and CUX2, as well as deep layer marker CTIP2 (green) in 30-day-old SNaP-derived neurons. Scale bar = 50 µm. **B**, Quantification of co-localization in Panel A (n = 3 wells). **C**, Punctate expression of Synapsin I (green) synaptic marker in 50-day-old SNaP-derived neurons co-cultured with primary mouse glia. MAP2 neurite marker in red. Scale bars = 50 µm. White arrow heads denote putative synapses. Data are represented as mean ± S.D.

### Supplemental Figure 4

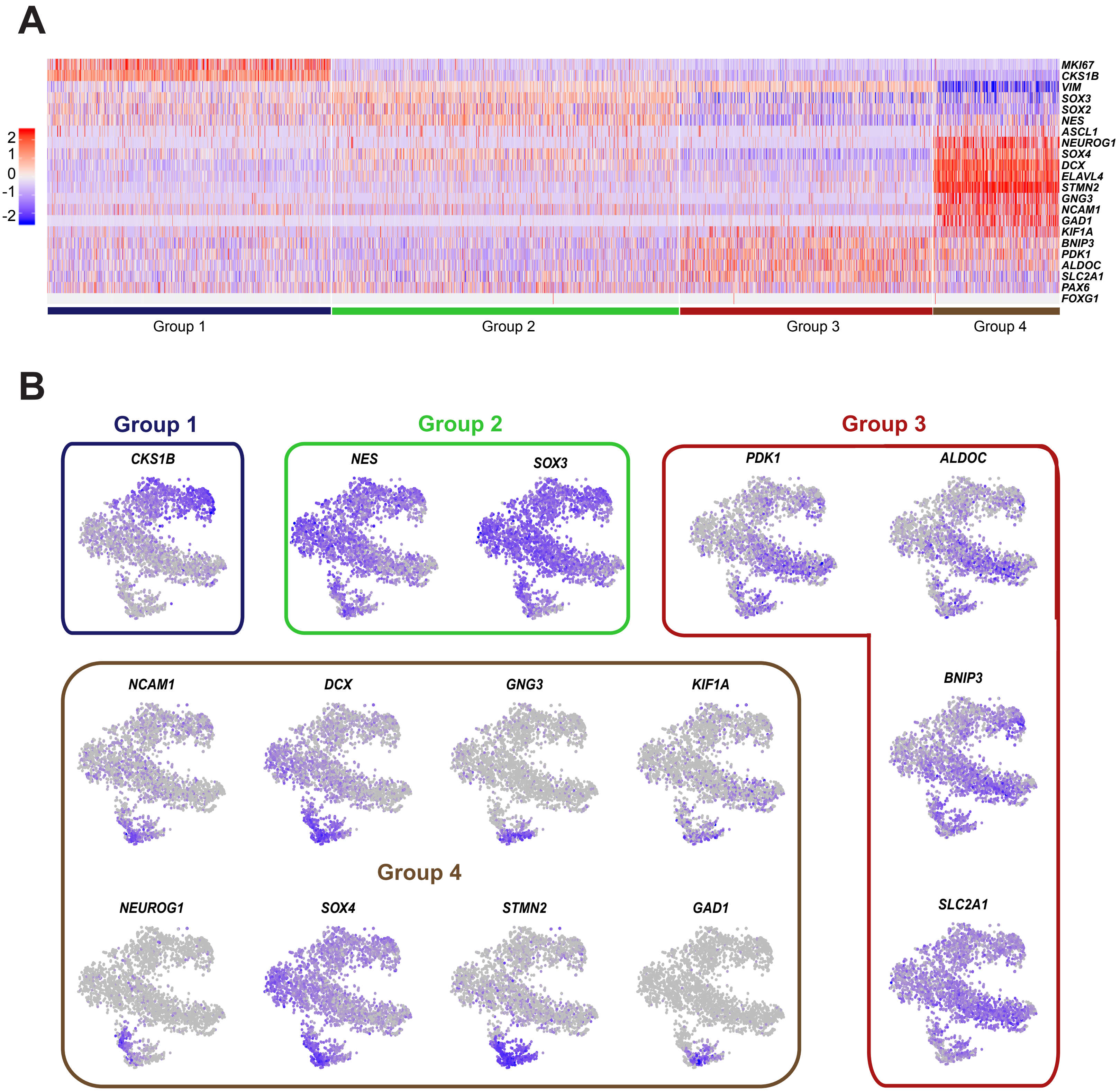

**Supplemental Figure 4 (Related to Figure 4): Single-cell characterization of SNaPspheres. A**, Heat map showing single-cell expression of cell type-marker transcripts of 7-day-old SW.1 SNaPspheres. **B**, TSNE plots of representative genes for each group.

### Supplemental Figure 5

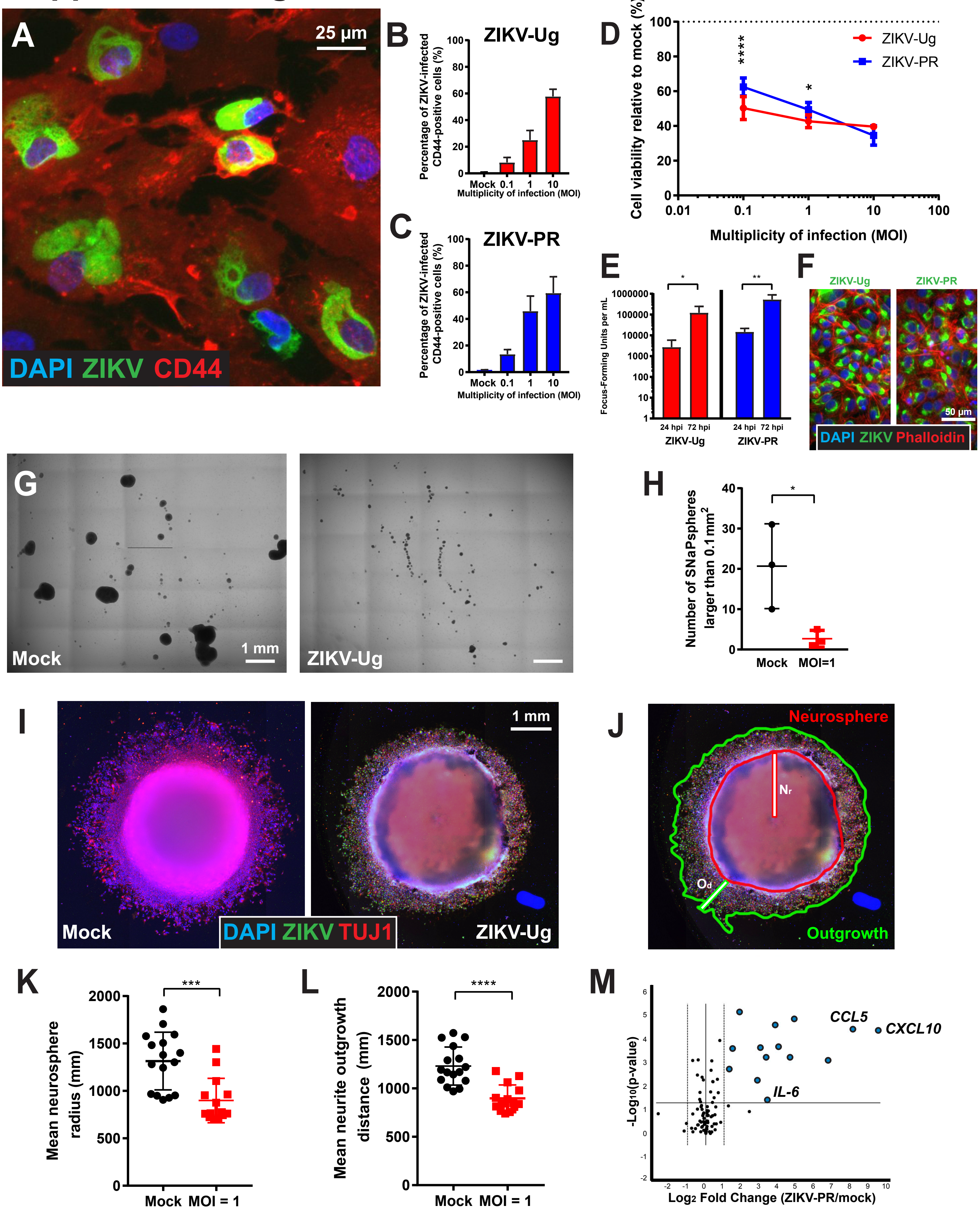

**Supplemental Figure 5 (Related to Figure 5): SNaP-derived glial cells are susceptible to ZIKV infection and viral-mediated cell death. A**, Representative image of SNaP-derived CD44-positive glia cells infected with ZIKV-Ug (MOI = 10) at 54 hpi. **B**-**C**, Quantification of SNaP-derived glia infections with (B) ZIKV-Ug and (C) ZIKV-PR at 54 hpi (n = 8 wells). **D**, Cell viability of SNaP-derived glia infected with ZIKV-Ug (red line) or ZIKV-PR (blue line) at 96 hpi. Viability is normalized to mock-infected controls (n = 8 wells). **E**, Quantification of ZIKV-Ug and ZIKV-PR viral RNA from conditioned media of infected (MOI = 10) SNaP-derived glia at 24 hpi and 72 hpi (n = 6 wells). **F**, ZIKV 4G2 envelope protein immunostaining of Vero cells at 24 hpi show that infected glia conditioned media contains infectious ZIKV particles. **G**-**H**, Infected SNaPs do not form neurospheres as shown in the (G) representative bright field images of mock or ZIKV-Ug infected (MOI = 1) SNaPspheres at 6 days post-plating. (H) Quantification of Panel G (n = 3 wells). **I**-**L**, ZIKV inhibits neurite outgrowth of SNaPspheres. (I) Representative images of mock and ZIKV-Ug infected (MOI = 1) SNaPspheres 48 hours post-plating. Sample neurosphere radius ($N_r$) and outgrowth distance ($O_d$) measurements shown in Panel J. ZIKV infection resulted in reduced (K) neurosphere radius and (L) neurite outgrowth of SNaPspheres (n = 14-16 spheres per condition). **M**, qPCR analysis of human antiviral response genes shows upregulation of multiple cytokines and chemokines (blue dots) in SNaPs at 60 hpi of MOI = 20 ZIKV-PR infection (n = 4). Two-way ANOVA with Sidak's tests for multiple comparisons (D) and the Student's t-test (E, H, K-L) were used for statistical analysis. Data are represented as mean ± S.D. *p<0.05, **p<0.01, ***p<0.001, ****p<0.0001.

### Supplemental Figure 6

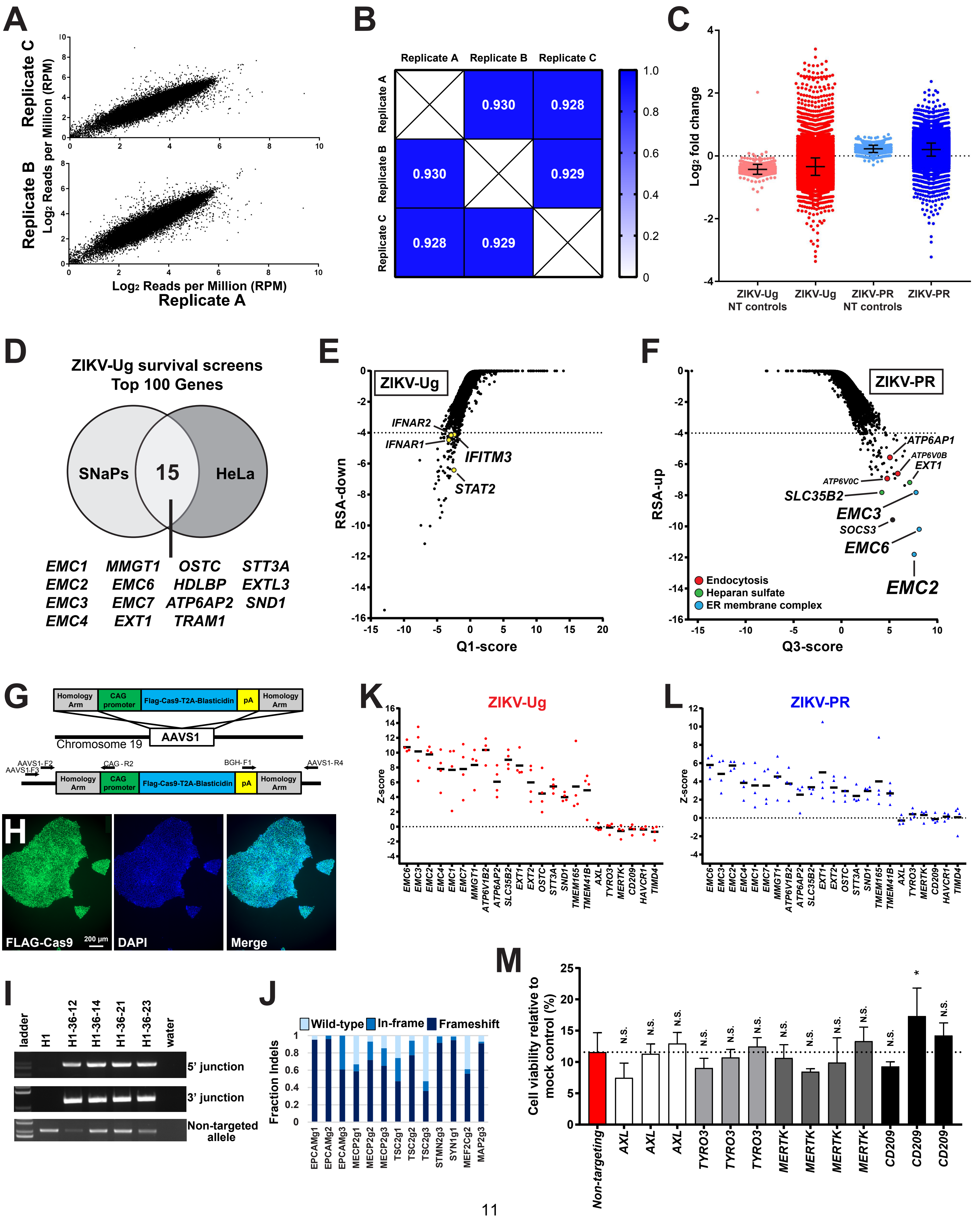

**Supplemental Figure 6 (Related to Figure 6): Quality controls and comparative results of ZIKV survival screens in SNaPs.** **A**, Plots comparing reads per million (RPM) of individual sgRNAs in Replicate A to Replicates B and C show linear relationship at Day 10 of ZIKV-Ug infection. **B**, Correlation matrix of Pearson r values across all replicates (range 0 to 1). **C**, $Log_2$ fold change of individual guides in the ZIKV-Ug (red dots) and ZIKV-PR (blue dots) survival screens. Non-targeting (NT) controls (pink, light blue dots) are also shown. **D**, Venn diagram showing the overlap of the top 100 genes from the ZIKV-Ug survival screen in SNaPs and HeLa cells (from *Savidis et al., 2016*). Co-hits listed below the diagram. **E**, Depleted genes from the ZIKV-Ug screen. Dashed line at $RSA_{up}$ = -4.0 denotes hit cutoff. **F**, Gene level results of the ZIKV-PR survival screen. Disruption of genes involved in the ER membrane complex (blue dots), endocytosis (red dots), and heparan sulfate biosynthesis (green dots) led to increased SNaP survival after 10 days of infection at MOI = 5. Dashed line at $RSA_{up}$ = -4.0 denotes hit cutoff. **G**, Diagram of AAVS1 targeting vector and site of genomic integration. **H**, Immunostaining using an anti-FLAG antibody for clone H1-36-23. Scale bar = 200 μM. **I**, PCR across the junctions for 4 clones, indicating proper targeting into one allele. **J**, Fraction indels in neurons differentiated from H1-36-23 clone and infected with lenti-gRNAs, measured by next-generation sequencing. **K**-**L**, Z-scores of $log_2$ fold change for top hits and TAM receptors (*AXL*, *TYRO3*, *MERTK*, *CD209*) from (K) ZIKV-Ug (red dots) and (L) ZIKV-PR (blue dots) SNaP survival screens. **M**, Validation experiments confirmed that ablation of TAM receptor proteins does not consistently protect from ZIKV-Ug mediated cell death at 120 hpi. Cell viability normalized to mock control values (n = 4 wells per sgRNA). One-way ANOVA with Dunnett's test for multiple comparisons was used for statistical analysis. Data are represented as mean ± S.D. N.S. = not significant, $*p<0.05$.

### Supplemental Figure 7

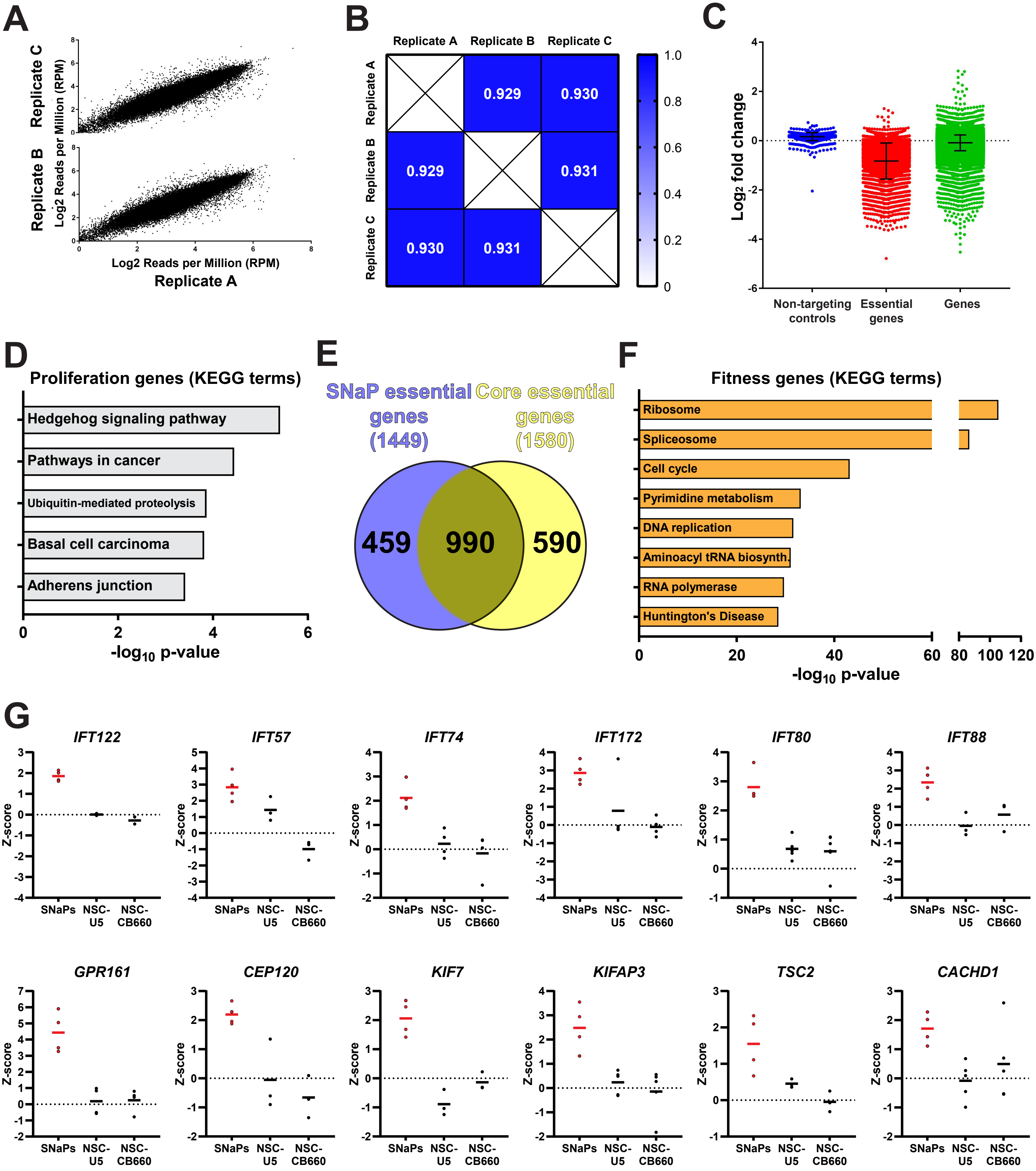

**Supplemental Figure 7 (Related to Figure 7): Quality controls and comparative results of SNaP fitness screen.** **A**, Plots comparing reads per million (RPM) of individual sgRNAs in Replicate A to Replicates B and C show linear relationship at Day 10 of SNaP fitness screen. **B**, Correlation matrix of Pearson r values across all replicates (range 0 to 1). **C**, $Log_2$ fold change of individual guides at Day 10 of the fitness screen. Non-targeting (NT) controls shown in blue, core essential genes (from Hart et al, 2015) shown in red, and remaining non-essential genes shown in green. **D**, KEGG pathway analysis results on 87 significantly-enriched genes. **E**, Venn diagram depicting overlap of 1,449 SNaP essential genes and 1,580 core essential genes (from Hart et al, 2015). **F**, KEGG pathway analysis results on 1,449 essential genes. **G**, Z-scores of individual sgRNAs from genome-wide CRISPR-Cas9 fitness screens in SNaPs and fetal-derived NSCs (from Toledo et al., 2015). Data are represented as mean ± S.D.

### Supplemental Table 1 (Related to Figure 1)

**Passage 1-2 SNaP immunostaining results**

| Line | %Nestin | %PAX6 | %SOX1 | %OCT4 |
|---|---|---|---|---|
| HS420 | 99.6 | 88.6 | 96.6 | 0.0 |
| GENEA016 | 99.8 | 94.0 | 95.7 | 0.0 |
| UM33-4 | 99.7 | 95.6 | 98.3 | 0.0 |
| UCLA_16 | 100.0 | 97.8 | 96.1 | 0.0 |
| WA14 | 100.0 | 95.9 | 95.9 | 0.0 |
| UM4-6 | 99.9 | 97.6 | 97.3 | 0.0 |
| HUES 63 HC | 99.9 | 95.1 | 98.1 | 0.0 |
| HUES 68 | 99.8 | 79.5 | 88.4 | 0.0 |
| WA01 | 100.0 | 88.6 | 87.1 | 0.0 |
| UCLA_10 | 100.0 | 91.2 | 98.7 | 0.0 |
| Genea43 | 99.8 | 87.2 | 96.4 | 0.0 |
| HUES 45 | 99.9 | 89.4 | 93.7 | 0.0 |
| UCLA_13 | 94.7 | 93.5 | 95.6 | 0.0 |
| UCLA_14 | 96.1 | 91.6 | 98.5 | 0.0 |
| Genea47 | 92.9 | 89.5 | 97.9 | 0.0 |
| HUES 44 | 94.5 | 95.3 | 98.1 | 0.0 |
| WA17 | 89.9 | 93.6 | 98.6 | 0.0 |
| CT4 | 93.0 | 94.1 | 99.3 | 0.0 |
| UCSF4 | 92.1 | 91.8 | 98.2 | 0.0 |
| WIBR5 | 86.5 | 73.3 | 88.9 | 0.0 |
| HUES 42 | 95.6 | 95.2 | 99.0 | 0.0 |
| CHB05 | 97.0 | 95.5 | 98.8 | 0.0 |
| UCLA_08 | 94.0 | 94.1 | 95.2 | 0.0 |
| HUES 72 | 86.6 | 82.7 | 95.7 | 0.1 |
| WA18 | 91.5 | 90.6 | 96.1 | 0.0 |
| HS401 | 81.0 | 86.5 | 96.8 | 0.0 |
| ESI017 | 91.9 | 83.8 | 96.2 | 0.1 |
| Genea52 | 96.8 | 68.8 | 90.2 | 0.0 |
| WIBR3 | 91.3 | 85.5 | 95.5 | 0.0 |
| WA22 | 91.6 | 86.4 | 98.0 | 0.0 |
| RUES1 | 90.6 | 85.9 | 95.7 | 0.0 |
| Genea57 | 82.0 | 76.7 | 95.0 | 0.0 |
| HUES 75 | 97.0 | 73.9 | 95.5 | 0.0 |
| HUES 62 | 93.5 | 88.5 | 98.2 | 0.0 |
| CT2 | 89.7 | 84.5 | 95.3 | 0.0 |
| WIBR6 | 97.2 | 97.9 | 97.8 | 0.0 |
| UM77-2 | 95.6 | 98.4 | 98.7 | 0.0 |
| UM 14-2 | 88.1 | 88.8 | 98.6 | 0.0 |
| WA09 | 91.0 | 95.7 | 98.3 | 0.0 |
| Mel4 | 89.3 | 88.0 | 96.9 | 0.0 |
| UCLA_01 | 92.6 | 95.2 | 97.3 | 0.0 |
| CHB9 | 92.5 | 93.1 | 97.0 | 0.0 |
| Genea42 | 95.7 | 96.3 | 93.8 | 0.0 |
| HUES 74 | 65.8 | 91.8 | 90.5 | 0.0 |
| WA7 | 94.9 | 94.9 | 97.0 | 0.0 |
| ESI049 | 93.9 | 90.4 | 92.5 | 0.0 |
| SW.1 | 99.0 | 94.5 | 94.1 | 1.5 |
| 8402 | 93.3 | 99.8 | 93.2 | 0.0 |

### Supplemental Table 2 (Related to Figure 5)

#### SNaP antiviral response to ZIKV: qPCR panel results

| | | ZIKV-Ug (MOI = 10) | | | | ZIKV-PR (MOI = 20) | | | |
|---|---|---|---|---|---|---|---|---|---|
| | | 8 hpi | | 60 hpi | | 8 hpi | | 60 hpi | |
| Gene symbol | Description | Fold Change | p-value | Fold Change | p-value | Fold Change | p-value | Fold Change | p-value |
| AIM2 | Absent in melanoma 2 | 0.98630 | 0.90827 | 0.99330 | 0.92670 | 0.96420 | 0.62723 | 1.01820 | 0.60378 |
| APOBEC3G | Apolipoprotein B mRNA editing enzyme, catalytic polypeptide-like 3G | 0.85060 | 0.77077 | 0.79890 | 0.29963 | 1.21150 | 0.68159 | 1.33290 | 0.28982 |
| ATG5 | ATG5 autophagy related 5 homolog (S. cerevisiae) | 1.13190 | 0.00278 | 1.07800 | 0.34757 | 1.09450 | 0.06466 | 1.12120 | 0.02497 |
| AZI2 | 5-azacytidine induced 2 | 1.06340 | 0.04259 | 1.06610 | 0.17585 | 0.98960 | 0.77546 | 1.04570 | 0.34391 |
| CARD9 | Caspase recruitment domain family, member 9 | 0.85360 | 0.80124 | 0.82400 | 0.85112 | 1.21420 | 0.57164 | 2.38630 | 0.10316 |
| CASP1 | Caspase 1, apoptosis-related cysteine peptidase (interleukin 1, beta, convertase) | 1.13890 | 0.50442 | ND | ND | 0.87150 | 0.92049 | 1.50580 | 0.37734 |
| CASP10 | Caspase 10, apoptosis-related cysteine peptidase | 0.79810 | 0.75110 | **7.11760** | **0.00550** | 1.61740 | 0.68174 | -1.25180 | 0.98894 |
| CASP8 | Caspase 8, apoptosis-related cysteine peptidase | 1.15830 | 0.01550 | 1.00050 | 0.98855 | 0.85190 | 0.05646 | 1.50140 | 0.01397 |
| CCL3 | Chemokine (C-C motif) ligand 3 | 1.03020 | 0.84279 | 1.42230 | 0.70049 | 0.49020 | 0.33131 | 1.67500 | 0.35030 |
| CCL5 | Chemokine (C-C motif) ligand 5 | 0.54160 | 0.48988 | **265.71830** | **0.00004** | 1.65500 | 0.87168 | **93.18700** | **0.00003** |
| CD40 | CD40 molecule, TNF receptor superfamily member 5 | 1.09870 | 0.83565 | 0.83430 | 0.33168 | 1.22440 | 0.60065 | 1.22350 | 0.42106 |
| CD80 | CD80 molecule | 0.35750 | 0.26994 | 0.92160 | 0.73558 | 1.74670 | 0.45163 | 1.08680 | 0.66654 |
| CD86 | CD86 molecule | 0.61380 | 0.55531 | 1.48390 | 0.39389 | 0.11680 | 0.03942 | 1.56470 | 0.36610 |
| CHUK | Conserved helix-loop-helix ubiquitous kinase | 0.85080 | 0.01536 | 0.91440 | 0.31636 | 0.88880 | 0.06848 | -1.01260 | 0.88886 |
| CTSB | Cathepsin B | 0.91170 | 0.04423 | 1.10010 | 0.11074 | 0.82050 | 0.00322 | -1.10190 | 0.14323 |
| CTSL | Cathepsin L1 | 0.97630 | 0.26276 | 1.00400 | 0.92799 | 0.93420 | 0.03280 | 1.02350 | 0.62460 |
| CTSS | Cathepsin S | 0.30660 | 0.14935 | 0.95700 | 0.33343 | 0.72260 | 0.84707 | -1.05380 | 0.80554 |
| CXCL10 | Chemokine (C-X-C motif) ligand 10 | 0.67920 | 0.74629 | **695.73250** | **0.00004** | 0.78240 | 0.62525 | **525.47860** | **0.00001** |
| CXCL11 | Chemokine (C-X-C motif) ligand 11 | 0.07880 | 0.09306 | **28.84610** | **0.00001** | 0.93600 | 0.97847 | **231.15640** | **0.00026** |
| CXCL9 | Chemokine (C-X-C motif) ligand 9 | 0.37950 | 0.36268 | 0.73390 | 0.43729 | 0.67630 | 0.36153 | 1.79990 | 0.15612 |
| CYLD | Cylindromatosis (turban tumor syndrome) | 1.11630 | 0.19462 | 1.28480 | 0.04854 | 1.21940 | 0.01068 | 1.50090 | 0.00616 |
| TKFC | Dihydroxyacetone kinase 2 homolog (S. cerevisiae) | 0.75210 | 0.00164 | 0.74730 | 0.00076 | 0.75280 | 0.00951 | -1.29370 | 0.01447 |
| DDX3X | DEAD (Asp-Glu-Ala-Asp) box polypeptide 3, X-linked | 0.77800 | 0.00012 | 0.90770 | 0.14918 | 0.75470 | 0.00184 | -1.12070 | 0.12710 |
| DDX58 | DEAD (Asp-Glu-Ala-Asp) box polypeptide 58 | 1.01700 | 0.73447 | **2.72650** | **0.00025** | 0.99200 | 0.84692 | 1.72360 | 0.00038 |
| DHX58 | DEXH (Asp-Glu-X-His) box polypeptide 58 | 0.98790 | 0.95892 | **9.80950** | **0.00059** | 0.77940 | 0.36719 | **11.26430** | **0.00032** |
| FADD | Fas (TNFRSF6)-associated via death domain | 1.19450 | 0.00375 | 1.00590 | 0.88151 | 1.10290 | 0.10652 | 1.11740 | 0.14088 |
| FOS | FBJ murine osteosarcoma viral oncogene homolog | 1.14290 | 0.12171 | **3.60400** | **0.00001** | 0.97600 | 0.61568 | **6.10030** | **0.00001** |
| HSP90AA1 | Heat shock protein 90kDa alpha (cytosolic), class A member 1 | 1.00860 | 0.72371 | 1.01140 | 0.81828 | 0.99140 | 0.85791 | -1.17340 | 0.01728 |
| IFIH1 | Interferon induced with helicase C domain 1 | 1.39600 | 0.50103 | **15.89290** | **0.00021** | 0.38260 | 0.22505 | **14.89510** | **0.00002** |
| IFNA1 | Interferon, alpha 1 | 0.22130 | 0.05717 | 0.58270 | 0.60335 | 1.52690 | 0.84363 | 1.03700 | 0.86709 |
| IFNA2 | Interferon, alpha 2 | 0.63610 | 0.58769 | 0.47880 | 0.37837 | 0.33130 | 0.26315 | 1.64200 | 0.36349 |
| IFNAR1 | Interferon (alpha, beta and omega) receptor 1 | 1.00360 | 0.92432 | 1.01040 | 0.85151 | 0.90490 | 0.11943 | -1.09860 | 0.27182 |
| IFNB1 | Interferon, beta 1, fibroblast | 2.61100 | 0.32096 | **13.96280** | **0.00003** | 0.49010 | 0.70114 | **18.65910** | **0.00029** |
| IKBKB | Inhibitor of kappa light polypeptide gene enhancer in B-cells, kinase beta | 0.81520 | 0.00051 | 0.92230 | 0.13555 | 0.81180 | 0.00173 | -1.36660 | 0.01179 |
| IL12A | Interleukin 12A (natural killer cell stimulatory factor 1, cytotoxic lymphocyte maturation factor 1, p35) | 1.03110 | 0.71288 | 1.41270 | 0.16143 | 1.11460 | 0.91976 | -1.25200 | 0.37401 |
| IL12B | Interleukin 12B (natural killer cell stimulatory factor 2, cytotoxic lymphocyte maturation factor 2, p40) | 0.17430 | 0.19072 | 0.43870 | 0.79209 | 3.95290 | 0.11522 | 2.45240 | 0.26001 |
| IL15 | Interleukin 15 | 0.72490 | 0.19785 | 1.18920 | 0.99946 | 0.98530 | 0.79036 | 2.35500 | 0.07339 |
| IL18 | Interleukin 18 (interferon-gamma-inducing factor) | 1.15690 | 0.09340 | 0.60860 | 0.14957 | 1.27860 | 0.18439 | 1.63940 | 0.14347 |
| IL1B | Interleukin 1, beta | 0.58170 | 0.00320 | 0.85690 | 0.53982 | 0.89200 | 0.69326 | **-2.01250** | **0.02775** |
| IL6 | Interleukin 6 (interferon, beta 2) | 0.97440 | 0.89012 | **10.45410** | **0.03899** | 1.25220 | 0.59025 | **24.21060** | **0.00518** |
| CXCL8 | Interleukin 8 | 1.67520 | 0.69002 | 2.35480 | 0.06794 | 0.74700 | 0.66933 | -1.47110 | 0.63466 |
| IRAK1 | Interleukin-1 receptor-associated kinase 1 | 0.94700 | 0.55142 | 0.72660 | 0.09317 | 0.87020 | 0.03464 | -1.07640 | 0.56213 |
| IRF3 | Interferon regulatory factor 3 | 0.95920 | 0.18315 | 1.03360 | 0.29798 | 0.90150 | 0.00520 | -1.31990 | 0.00193 |
| IRF5 | Interferon regulatory factor 5 | 1.84180 | 0.05842 | 1.37260 | 0.39321 | 1.22040 | 0.28539 | 1.36430 | 0.23891 |
| IRF7 | Interferon regulatory factor 7 | 0.99560 | 0.91091 | 1.36870 | 0.03061 | 0.77610 | 0.00844 | 1.05470 | 0.32193 |
| ISG15 | ISG15 ubiquitin-like modifier | 1.16780 | 0.02142 | **7.98570** | **0.00023** | 1.04690 | 0.80801 | **4.49950** | **0.00017** |
| JUN | Jun proto-oncogene | 1.21500 | 0.00541 | 1.69770 | 0.00012 | 0.99990 | 0.98590 | 1.81940 | 0.00046 |
| MAP2K1 | Mitogen-activated protein kinase kinase 1 | 0.84020 | 0.00353 | 0.97230 | 0.39189 | 0.85600 | 0.00149 | 1.08460 | 0.09444 |
| MAP2K3 | Mitogen-activated protein kinase kinase 3 | 0.97080 | 0.59410 | 1.03000 | 0.58397 | 0.90280 | 0.06694 | -1.19410 | 0.00099 |
| MAP3K1 | Mitogen-activated protein kinase kinase kinase 1 | 0.81520 | 0.00294 | 1.03840 | 0.66374 | 0.87130 | 0.00338 | 1.01510 | 0.87214 |
| MAP3K7 | Mitogen-activated protein kinase kinase kinase 7 | 1.03330 | 0.38753 | 1.21940 | 0.00210 | 0.90640 | 0.00761 | -1.03680 | 0.45148 |
| MAPK1 | Mitogen-activated protein kinase 1 | 0.90650 | 0.00528 | 0.72610 | 0.00336 | 0.89500 | 0.00012 | -1.20840 | 0.04485 |
| MAPK14 | Mitogen-activated protein kinase 14 | 0.99620 | 0.84225 | 0.98230 | 0.72838 | 0.98010 | 0.40683 | -1.02920 | 0.57524 |
| MAPK3 | Mitogen-activated protein kinase 3 | 0.89510 | 0.01931 | 0.94500 | 0.41470 | 0.82530 | 0.00152 | 1.07630 | 0.66194 |
| MAPK8 | Mitogen-activated protein kinase 8 | 1.08940 | 0.07299 | 1.06910 | 0.29969 | 1.04640 | 0.23806 | 1.15840 | 0.06795 |
| MAVS | Mitochondrial antiviral signaling protein | 0.77160 | 0.01143 | 0.86720 | 0.00415 | 0.77770 | 0.00040 | -1.47570 | 0.01118 |
| MEFV | Mediterranean fever | 0.48960 | 0.79791 | 0.16400 | 0.14644 | 0.45990 | 0.37317 | -1.56700 | 0.88772 |
| MX1 | Myxovirus (influenza virus) resistance 1, interferon-inducible protein p78 (mouse) | 1.16900 | 0.44744 | **24.27830** | **0.00059** | 0.84390 | 0.26248 | **13.76930** | **0.00018** |
| MYD88 | Myeloid differentiation primary response gene (88) | 0.83350 | 0.01411 | 1.19910 | 0.01744 | 0.77280 | 0.00519 | -1.19560 | 0.00495 |
| NFKB1 | Nuclear factor of kappa light polypeptide gene enhancer in B-cells 1 | 0.95890 | 0.10531 | 1.09290 | 0.17302 | 0.84800 | 0.03612 | 1.03590 | 0.78498 |
| NFKBIA | Nuclear factor of kappa light polypeptide gene enhancer in B-cells inhibitor, alpha | 1.02450 | 0.64858 | 1.31070 | 0.00643 | 0.98600 | 0.68881 | 1.34780 | 0.00121 |
| NLRP3 | NLR family, pyrin domain containing 3 | 1.23980 | 0.88506 | 1.75860 | 0.33719 | 0.67020 | 0.35832 | 1.78970 | 0.15743 |
| NOD2 | Nucleotide-binding oligomerization domain containing 2 | 1.04120 | 0.98433 | 1.26180 | 0.50464 | 1.52680 | 0.89755 | -1.62790 | 0.24144 |
| OAS2 | 2'-5'-oligoadenylate synthetase 2, 69/71kDa | ND | ND | **103.87040** | **0.00078** | 0.46050 | 0.33701 | **332.80030** | **0.00062** |
| PIN1 | Peptidylprolyl cis/trans isomerase, NIMA-interacting 1 | 1.00300 | 0.87246 | 1.04530 | 0.13393 | 0.90680 | 0.01412 | -1.05930 | 0.22480 |
| PSTPIP1 | Proline-serine-threonine phosphatase interacting protein 1 | 0.74480 | 0.01209 | 0.79980 | 0.01826 | 0.82490 | 0.09283 | -1.41670 | 0.01124 |
| PYCARD | PYD and CARD domain containing | 1.02840 | 0.96006 | 0.87770 | 0.50916 | 0.99960 | 0.94976 | 1.35860 | 0.30579 |
| PYDC1 | PYD (pyrin domain) containing 1 | 1.27540 | 0.31309 | 1.45620 | 0.07544 | 1.14270 | 0.91134 | -1.34970 | 0.18460 |
| RELA | V-rel reticuloendotheliosis viral oncogene homolog A (avian) | 1.05060 | 0.15232 | 1.02990 | 0.63473 | 1.02060 | 0.50470 | 1.03120 | 0.75789 |
| RIPK1 | Receptor (TNFRSF)-interacting serine-threonine kinase 1 | 0.90230 | 0.00923 | 1.02870 | 0.65389 | 0.85210 | 0.01761 | 1.05310 | 0.72240 |
| SPP1 | Secreted phosphoprotein 1 | **3.48090** | **0.00005** | 1.28860 | 0.17808 | 1.98890 | 0.00193 | 1.18120 | 0.56360 |
| STAT1 | Signal transducer and activator of transcription 1, 91kDa | 1.07830 | 0.10679 | **2.43820** | **0.00184** | 1.02950 | 0.58303 | 1.56120 | 0.00044 |
| SUGT1 | SGT1, suppressor of G2 allele of SKP1 (S. cerevisiae) | 1.24960 | 0.00160 | 1.30000 | 0.00634 | 1.10370 | 0.04179 | -1.02080 | 0.60879 |
| TBK1 | TANK-binding kinase 1 | 1.03230 | 0.34188 | 0.67420 | 0.62574 | 1.00130 | 0.91961 | -1.39740 | 0.00496 |
| TICAM1 | Toll-like receptor adaptor molecule 1 | 1.14750 | 0.55745 | 1.60220 | 0.01879 | 0.77550 | 0.62728 | -1.44660 | 0.27799 |
| TLR3 | Toll-like receptor 3 | 0.98250 | 0.89037 | 0.74760 | 0.43388 | 0.91710 | 0.77211 | -1.54850 | 0.13554 |
| TLR7 | Toll-like receptor 7 | 1.37170 | 0.78576 | 1.14940 | 0.85552 | 0.29740 | 0.62851 | 6.02710 | 0.13297 |
| TLR8 | Toll-like receptor 8 | 6.99520 | 0.06557 | 0.61130 | 0.85503 | 0.38620 | 0.48272 | 1.34760 | 0.83565 |
| TLR9 | Toll-like receptor 9 | 0.40110 | 0.38712 | 1.25140 | 0.85945 | 0.70530 | 0.25453 | 1.22130 | 0.91049 |
| TNF | Tumor necrosis factor | ND | ND | 5.21570 | 0.11815 | 0.86520 | 0.15761 | **9.28140** | **0.00133** |
| TRADD | TNFRSF1A-associated via death domain | 1.54110 | 0.11676 | 1.22370 | 0.19664 | 0.69600 | 0.10813 | -1.16510 | 0.42798 |
| TRAF3 | TNF receptor-associated factor 3 | 0.92430 | 0.01174 | 0.93080 | 0.22986 | 0.89580 | 0.01796 | -1.03970 | 0.58307 |
| TRAF6 | TNF receptor-associated factor 6 | 1.08920 | 0.12718 | 1.04540 | 0.50932 | 0.98300 | 0.72330 | 1.08750 | 0.51953 |
| TRIM25 | Tripartite motif containing 25 | 0.99890 | 0.96612 | 1.55100 | 0.00508 | 0.95440 | 0.08270 | 1.02060 | 0.77041 |

**BOLD** denotes p<0.05 and Fold change > +/- 2.0 **ND** = Not detected

#### SUPPLEMENTAL EXPERIMENTAL PROCEDURES

*Stem cell culture*: Human ESCs and iPSCs were maintained in mTeSR media (Stem Cell Technologies, 85850) on Geltrex basement membrane matrix (1:100; Life Technologies, A1413301). Cells were split every 4-5 days (when they reached 80-90% confluency) using a 15 minute/37°C incubation in Accutase (Innovative Cell Technologies, AT104) followed by 1:10 dilution in mTeSR. For each passage, media was supplemented with ROCK inhibitor Y-27632 (10 μM; Stemgent, 04-0012) for 24 hours after plating.

*Viral transduction:* TetO-Ngn2-Puromycin and Ubq-rtTA constructs were obtained from the Wernig lab (Stanford) before being packaged as high-titer lentiviruses (Alstem, Richmond, CA). When hPSCs reached 80-100% confluency, they were dissociated with Accutase before being re-suspended in lentivirus-containing mTeSR media supplemented with Y-27632 at a range of MOI = 1 to MOI = 3. Cells were then plated at 100,000-150,000 cells/cm$^2$ on Geltrex-coated plates. After 18-24 hours, cells were fed with mTeSR media and maintained as described above. Transduced cells were maintained for up to 10 passages for inductions.

*Induction of SNaPs from human PSCs*: Human PSCs were dissociated and plated at 75,000 cells/cm$^2$ on Geltrex matrix in mTeSR media supplemented with Y-27632. After 12-24 hours, cells were fed with Induction Media (Day 1): DMEM/F12 (ThermoFisher, 11320082), Glutamax (1:100; ThermoFisher, 10565018), 20% Glucose (1.5% v/v), N2 Supplement (1:100, ThermoFisher, 17502048), Doxycycline (2 μg/mL; Sigma-Aldrich, D9891), LDN-193189 (200 nM; Stemgent, 04-0074), SB431542 (10 μM; Tocris, 1614), and XAV939 (2 μM; Stemgent, 04-00046). After 24 hours in Induction Media, cells were fed with Selection Media (Day 2): DMEM/F12, Glutamax (1:100), 20% Glucose (1.5% v/v), N2 Supplement (1:100), Doxycycline (2 μg/mL), puromycin (5 μg/mL; ThermoFisher, A1113803), LDN-193189 (100 nM), SB431542 (5 μM), and XAV939 (1 μM). After 24 hours in Selection Media, SNaPs were dissociated with Accutase and replated at 120,000 cells/cm$^2$ on Geltrex-coated plates in SNaP maintenance media supplemented with puromycin and Y-27632 (Day 3): DMEM/F12, Glutamax (1:100), MEM-NEAA (1:100; Life Technologies, 10370088), B27 minus Vitamin A (1:50; Life Technologies, 12587010), N2 Supplement (1:100; Life Technologies, 17502048), recombinant human EGF (10 ng/mL; R&D Systems, 236-EG-200), recombinant human basic FGF (10 ng/mL; Life Technologies, 13256029), puromycin (5 μg/mL), and Y-27632 (10 μM). Starting 12-24 hours after passaging, SNaPs were fed daily with SNaP maintenance media lacking Y-27632 and puromycin. SNaPs were passaged every 5-7 days. All SNaP passages throughout this manuscript included overnight Y-27632 treatment.

*Immunostaining*: SNaPs were washed with 1X PBS and then fixed with 4% paraformaldehyde for 15 minutes at room temperature before three more washes with 1X PBS. Cells were permeabilized with 0.1% Triton for 15 minutes and then blocked with 10% normal donkey serum diluted in 1X PBS for 1 hour at room temperature followed by an overnight 4°C incubation in primary antibody diluted in blocking solution: Mouse anti-Nestin (1:1000; Stem Cell Technologies, 60091), Rabbit anti-PAX6 (1:500; Stem Cell Technologies, 60094), Mouse anti-OCT4 (1:1000; Stem Cell Technologies, 60093), Rabbit anti-SOX1 (1:1000; Stem Cell Technologies, 60095), Mouse anti-SOX2 (1:100; R&D Systems, MAB2018), Rabbit anti-ZO1 (1:200; Life Technologies, 617300), Rabbit anti-FOXG1 (1:400; Abcam, 18259), and/or Rabbit anti-KI67 (1:500; ThermoFisher, MA5-14520). After 3 washes in 1X PBS at room temperature, cells were incubated for 2-4 hours at room temperature in secondary antibody diluted in blocking solution: Donkey anti-mouse Alexa647 (1:1000; Life Technologies, A-31571) and/or Donkey anti-rabbit Alexa555 (1:1000; Life Technologies A-31572). Cells were washed once with 1X PBS followed by a 5 minute incubation in 4', 6-Diamidino-2-Phenylindole Dihydrochloride (DAPI, 1:5000; Life Technologies, D1306). Finally, cells were washed twice more with 1X PBS prior to imaging. For each well of a 96-well plate, 4-8 fluorescent images were captured using the Cytation 3 cell imaging multi-mode reader (BioTek Instruments; Winooski, VT). All images were then processed using the CellProfiler imaging analysis software (Carpenter et al., 2006) to quantify the percentage of NPC marker-positive cells.

*Flow cytometry analysis of human pluripotent stem cells and SNaPs:* Human pluripotent stem cells and SNaPs were stained with OCT3/4 antibody (BD Biosciences, 560794) following the manufacturer's instructions contained in the Human Pluripotent Stem Cell Transcription Factor Analysis Kit (BD Biosciences, 560589). Briefly, hPSCs and SNaPs were dissociated with Accutase and fixed using BD Cytofix at a concentration of 1x10$^7$ cells/mL for 20 minutes at room temperature. Fixed cells were washed twice with 1X PBS and permeabilized using 1X BD Perm/Wash at a concentration of 1x10$^7$ cells/mL for 10 minutes at room temperature. 1x10$^6$ fixed/permeabilized cells were stained with OCT3/4 antibody at a concentration of 1x10$^7$ cells/mL for 20 minutes at room temperature in the dark. Stained cells were washed twice with 1X BD Perm/Wash, resuspended in 1X PBS, and kept on ice in the dark until analysis. Cells were passed through a filter-top 12x75 mm polystyrene tube just before analysis on a BD FACSAria II (BD

Biosciences; San Jose, CA). Data was presented with OCT3/4 on the x-axis (PerCP-Cy5.5) and the empty channel mCFP-A on the y-axis.

*Neural progenitor cell culture (LSX-2w)*: When SW.1 hiPS cells reached 80-90% confluency (Day 0), neuroectodermal differentiation media A was added to induce NPCs: DMEM/F12 (47% v/v), Neurobasal media (47% v/v; Life Technologies, 21103049), Glutamax (1:50), MEM-NEAA (1:100), B27 (1:50; Life Technologies, 17504044), N2 Supplement (1:100), SB431542 (10 μM), LDN (100 nM), and XAV-939 (2 μM). Cells underwent complete media exchanges daily. At Day 14, cells were harvested for RNA extraction and subsequent qPCR experiments.

*Neural progenitor cell culture (LSR-3w)*: When SW.1 hiPS cells reached 80-90% confluency (Day 0), neuroectodermal differentiation media A with retinoic acid (1 μM; Sigma, R2625) in place of XAV-939. Cells underwent complete media exchanges daily. Starting on Day 7, cells were fed daily with neuroectodermal differentiation media B: DMEM/F12 (47% v/v), Neurobasal media (47% v/v), Glutamax (1:50), MEM-NEAA (1:100), B27 (1:50), N2 Supplement (1:100), and Retinoic acid (1 μM). At Day 21, cells were harvested for RNA extraction for subsequent qPCR experiments.

*SNaP developmental qPCR:* SW.1 hiPS cells, SNaPs, and NPCs were harvested in 350 μL of RTL Plus (Qiagen, 1053393) per well of a 24-plate. RNA was extracted from the samples using the RNeasy Plus Micro Kit (Qiagen, 74034). Purified RNA was then used as input for the iScript cDNA Synthesis reaction (Bio-Rad, 1708891) and the product was diluted 1:5 in nuclease-free water. For each sample, 1 μL of cDNA was added to iTaq Universal SYBR Green Supermix (Bio-Rad, 1725124) that contained 500 nM of forward and reverse primers in a final volume of 20 μL per well of a 384-well plate. Primers were manufactured by Integrated DNA Technologies (*NESTIN*: 5'-CTG CTA CCC TTG AGA CAC CTG-3' and 5'-GGG CTC TGA TCT CTG CAT CTA C-3'; *PAX6*: 5'-AAC GAT AAC ATA CCA AGC GTG T-3' and 5'-GGT CTG CCC GTT CAA CAT C-3'; *SOX1*: 5'-CCA CAT CCT AAT CTT GAG CCA-3' and 5'-CTG ACG TCC ACT CTC AGT CT-3'; *OCT4*: 5'-CCA AGG AAT AGT CTG TAG AAG TGC-3' and 5'-TGC ATG AGT CAG TGA ACA GG-3'; *FOXG1*: 5'-CGT CCA CCA TAT AGT TCC ATG A-3' and 5'-TGA CTG CTT TGC CAT TTC ATT C-3'; *SOX2*: 5'-CTT GAC CAC CGA ACC CAT-3' and 5'-GTA CAA CTC CAT GAC CAG CTC-3'; *GAPDH*: 5'-TTG TCA AGC TCA TTT CCT GGT ATG-3' and 5'-TCC TCT TGT GCT CTT GCT GG-3'). qPCR reactions were run for 40 cycles on the CFX384 Touch Real-Time PCR Detection System (Bio-Rad; Hercules, CA). All samples were run in triplicate, and results were normalized to a GAPDH control run in duplicate. ΔΔCt values were calculated and plotted to show relative expression.

*Single-cell RNA sequencing:* For monolayer culture, SNaPs were differentiated for 15 days in N2/B27 medium and dissociated with a 15 minute/37°C Accutase treatment followed by 1:1 dilution with N2/B27 medium. For SNaPsphere culture, 10 spheres were pooled and dissociated via Neural Tissue Dissociation Kit (P) (Miltenyi, 130-092-628) on a GentlMACS Octo Dissociator with Heaters (Miltenyi, 130-096-0427). Samples were filtered via 40 μm tip filters (BelArt, H13680-0040) and centrifuged at 400xg for 5 minutes. Cells were resuspended to 1 million cells/mL and run through the 10X Chromium, Version 2 single cell RNA-seq pipeline per vendor's instructions.

Reverse transcription master mix was prepared from 50 μL RT reagent mix (10X, 220089), 3.8μL RT primer (10X, 310354), 2.4 μL additive A (10X, 220074), and 10 μL RT enzyme mix (10X, 220079). Cell solution and master mix were mixed to achieve a target cell number of 3,000 cells per lane via 10X protocol. 90μL sample was loaded onto the 10X Single Cell 3' Chip along with 40μL barcoded gel beads and 270 μL partitioning oil, and the microfluidics system was run to match gel beads with individual cells. The droplet solution was then slowly transferred to an 8-tube strip, which was immediately incubated for 45 minutes at 53°C to perform reverse transcription, then 5 minutes at 85°C. The sample was treated with 125 μL recovery agent (10X, 220016), which was then removed along with the partitioning oil. 200 μL of cleanup solution containing 4 μL DynaBeads MyOne Silane Beads (Thermo Fisher, 37002D), 9 μL water, 182 μL Buffer Sample Clean Up 1 (10X, 220020), and Additive A (10X, 220074) was added to the sample, and the solution was mixed 5 times by pipetting and allowed to incubate at room temperature for 10 minutes. Beads were separated via magnetic separator and supernatant was removed. While still on the magnetic separator, the beads were then washed twice with 80% ethanol. The separator was then removed and the beads were resuspended in 35.5 μL elution solution consisting of 98 μL Buffer EB (Qiagen, 19086), 1 μL 10% Tween 20 (Bio-Rad, 1610781), and 1 μL Additive A (10X, 220074). The solution was then incubated for 1 minute at room temperature, and placed back onto the magnetic separator. 35 μL of eluted sample was transferred to a new tube strip. cDNA amplification reaction mix was prepared from 8 μL water, 50 μL Amplification Master Mix (10X, 220125), 5 μL cDNA Additive (10X, 220067), and 2 μL cDNA Primer Mix (10X, 220106). 65 μL of amplification master mix was added to the sample, mixed 15 times via pipetting, and briefly centrifuged. The sample then underwent 12 amplification cycles (15 seconds at 98°C, 20 seconds at 67°C, 1 minute at 72°C). SPRIselect beads (Beckman Coulter, B23318) were then applied at 0.6X, and solution was mixed 15 times via pipetting. The sample was incubated at room temperature for 5 minutes, placed onto a magnetic separator, and washed twice with 80% ethanol. Sample was air dried for 2 minutes and eluted in 40.5 µL Buffer EB. cDNA yield was measured on a 2100 Bioanalyzer (Agilent, G2943CA) via DNA High Sensitivity Chip (Agilent, 5067-4626).

Fragmentation mix was prepared at 4°C from 10 µL fragmentation enzyme blend (10X, 220107) and 5 µL fragmentation buffer (10X, 220108). 35 µL of sample cDNA was then added to the chilled fragmentation mix. Sample was incubated for 5 minutes at 32°C, then 30 minutes at 65°C to conduct enzymatic fragmentation, end repair, and A-tailing. Sample was then purified using 0.6X SPRIselect reagent (see above). Adaptor ligation mix was prepared from 17.5 µL water, 20µL Ligation Buffer (10X, 220109), 10 µL DNA Ligase (10X, 220110), and 2.5 µL Adaptor Mix (10X, 220026). The ligation mix was added to 50 µL of sample and mixed 15 times via pipetting. Sample was then incubated for 15 minutes at 20°C to conduct the ligation. The sample was purified using 0.8X SPRIselect reagent (see above). Sample index PCR mix was prepared from 8 µL water, 50 µL Amplification Master Mix (10X, 220125), and 2 µL SI-PCR Primer (10X, 220111). 60 µL sample index PCR mix, 30 µL purified sample, and 10 µL of sample index (10X, 220103) were combined and mixed 15 times via pipetting. Indexing was conducted via 9 cycles of 20 seconds at 98°C, 30 seconds at 54°C, then 20 seconds at 72°C. Sample was purified via double-sided SPRI selection at 0.6X and 0.8X, respectively. Sample was then quantified via DNA High Sensitivity Chip. Additional quantification was conducted via KAPA Library Quantification Kit (Illumina, KK4828-07960166001). Sample was diluted at 10-fold increments from 1:100 to 1:1,000,000, and mixed 1:9 with KAPA qPCR mix. qPCR was conducted on a Viia7 qPCR machine (Life Technologies).

Sample was then sequenced on a HiSeq 4000 (Illumina) using 2 x 50-cycle SBS kits (Illumina, FC-410-1001). Sample library was diluted to 2nM in EB buffer with 1% PhiX spike-in. 5 µL nondenatured library was then mixed with 5 µL 0.1N NaOH, then vortexed and briefly centrifuged. Denaturing was conducted at room temperature for exactly 8 minutes, then stopped via addition of 5 µL 200 mM Tris-HCl pH 8.0 (Fluka, 93283). Sample was mixed, briefly centrifuged, and placed on ice. ExAmp reaction mix (Illumina, PE-410-1001) was prepared, added to the sample, and clustering was done on a HiSeq 4000 flow cell via cBot2 (Illumina). The library was then sequenced with paired-end reagents, with 26xRead 1 cycles, 8xi7 index cycles, and 98xRead 2 cycles.

The 10X Cell Ranger 1.3.1 pipeline was utilized to convert raw BCL files to cell-gene matrices. Briefly, the Illumina bcl2fastq script conducted the initial demultiplexing. FASTQ files were then aligned, UMI-filtered, and barcodes were matched via the CellRanger count script. The GRCh37.75 human reference genome was used for alignment. After filtering out barcodes with very few matching transcripts, a total of 3,276 SNaPsphere cells and 2,167 SNaP-derived monolayer cells were adequately sequenced. An average of 144,318 reads were mapped per SNaPsphere cell and 207,402 reads were mapped per SNaP-derived monolayer cell.

*Single cell transcriptional analysis:* Single cell RNAseq datasets were analyzed in R using Seurat2. Cell-gene matrices were log normalized, and cells with >10% mitochondrial reads of >7000 unique genes were filtered out to reduce the number of dead or doublet cells within the dataset. Variable genes were identified and used to determine the top 15 principal components, which were used for the subsequent analysis. Graph-based clustering was used to approximate different cell groups, and t-stochastic neighborhood embedding (TSNE) analysis used for 2-dimensional representation. Differential expression between clusters was determined by the Wilcoxon rank sum test.

*SNaP differentiation immunostaining*: SNaPs were plated on Geltrex at 10,000-15,000 cells/cm$^2$ in base media [DMEM/F12, Glutamax (1:50), MEM-NEAA (1:100), B27 (1:50), N2 Supplement (1:100)] or base media with 10% fetal bovine serum (GE Healthcare, 16777-014) or Astrocyte Media (ScienCell, 1801) in the presence of Y-27632 (10 µM). The media was exchanged the following day to remove Y-27632, and cells were then fed 2-3 times a week for 14 days (base and 10% FBS) or 20-60 days (Astrocyte media). Cells cultured in Astrocyte Media were passaged weekly and re-plated at 15,000 cells/cm$^2$ without Y-27632. To determine cell identity, SNaP-derived cells were immunostained and quantified as described above. The following primary antibodies were used for these experiments: Mouse anti-HuC/D (1:200; Life Technologies, A-21271), Rat anti-CD44 (1:400; eBioScience, 17-0441-82), Rabbit anti-S100β (1:1000; Sigma Aldrich, S2532), Rabbit anti-GFAP (1:100; Millipore, AB5804), Chicken anti-MAP2 (1:500; Abcam, ab5392), Rabbit anti-Synapsin I (1:1000; Millipore, AB1543), Rabbit anti-BRN2 (1:300; Abcam, ab137469), Rabbit anti-CUX2 (1:200; Abcam, ab130395), and/or Rat anti-CTIP2 (1:1000; Abcam, ab18465). Donkey anti-Mouse Alexa555 (1:1000; Life Technologies, A-31570), Donkey anti-Mouse Alexa647 (1:1000; Life Technologies, A-31571), Donkey anti-Rabbit Alexa555 (1:1000; Life Technologies, A-31572), Goat anti-Rat Alexa555 (1:1000, Life Technologies, A-21434), and/or Goat anti-Chicken Alexa647 (1:1000; Life Technologies, A-21449) were used as secondary antibodies.

*Clonal assay for self-renewal:* SNaPs were plated as single cells in SNaP maintenance media plus Y-27632 on Geltrex-coated 96-well plates using a BD FACSAria II. Cells were then fed daily with SNaP complete media for two weeks. At this point, most wells were fixed and stained with Mouse anti-NESTIN and Rabbit anti-PAX6 for quantification of proliferation. The remaining wells were dissociated and re-plated as single cells in SNaP maintenance media plus Y-27632. Media was changed the following day to spontaneous differentiation media (base media plus B27/N2) and fed 2-3 times a week for two weeks. SNaPs were then fixed and processed for immunostaining as previously described.

*Multi-electrode array (MEA)*: SNaPs were plated on Geltrex at 15,000 cells/cm$^2$ in base media supplemented with Y-27632. The media was exchanged the following day to remove Y-27632. After 5-6 days in culture, the cells were fed with base media containing DAPT (5 µM; DNSK International). Two days later, the partially differentiated SNaPs were then dissociated and re-plated at 15,000 cells/cm$^2$ in DAPT-containing base media. One week later, the post-mitotic cells were dissociated and co-cultured with primary mouse glia (23,000 glia + 13,000 neurons per well) on a Geltrex-coated 12-well MEA plate (Axion Biosystems, M768-GL1-30Pt200) in Neurobasal complete media [Neurobasal media (97% v/v; Life Technologies 21103049), Glutamax (1:100), 20% Glucose (1.5% v/v), MEM-NEAA (1:200), B27 (1:50), BDNF (10 ng/mL), CTNF (10 ng/mL), and GDNF (10 ng/mL)]. Cells were fed 2-3 times per week with partial exchanges to reach a final volume consisting of 80% fresh media and 20% conditioned media. Five minutes of neuronal activity was measured weekly using the Maestro 12-well 64 electrodes per well micro-electrode array (MEA) plate system (Axion Biosystems, Atlanta, GA). After approximately 50 days in co-culture, synaptic contents were assessed using pharmacological blockers of neurotransmitter receptors. More specifically, baseline activity was measured for 5 minutes prior to the addition of NBQX (10 µM), D-APV (50 µM), or Picrotoxin (50 µM) directly to the conditioned media. After a brief 5-10 incubation, neuronal activity was again measured for 5 minutes. Data was analyzed using the Axion Integrated Studio 2.4.2 and the Neural Metric Tool (Axion Biosystems).

*SNaPsphere formation*: SNaPs were dissociated in Accutase and plated at 18,000 cells/well of an Ultra-Low Attachment 96-well Round Bottom plate (Corning, 7007) in 150 µL of SNaP maintenance media supplemented with Y-27632 (50 µM). Two days later (Day 2), 75 µL of conditioned media was removed from each well and replaced with 150 µL of fresh SNaP maintenance media supplemented with Y-27632 (50 µM). The following day (Day 3), 125 µL of conditioned media was removed from each well using the "blast" technique and replaced with 150 µL of fresh SNaP maintenance media without Y-27632 (Salick et al., 2017). From Day 4 onward, 150 µL of conditioned media was removed every other day from each well using the "blast" technique and replaced with 150 µL of fresh SNaP maintenance media. SNaPspheres were measured using the Cytation 3 cell imaging multi-mode reader using the 4X bright field objective. All images were then processed using the CellProfiler imaging analysis software to stitch the images and quantify the two-dimensional area of each SNaPsphere.

*Lightsheet imaging:* CUBIC solution was prepared from urea (250 mg/mL; JT Baker, 4204-01) and N,N,N',N'-tetrakis(2-hydroxypropyl) ethylenediamine (THEED) (250 mg/mL; Sigma, 122262) dissolved at 50°C in a 15% triton X-100 (Alfa Aesar, J66624)/85% $H_2O$ mixture. RIMS solution was prepared from 30 mL of .02 M phosphate buffer (diluted from Sigma, P5244), 40g Histodenz (Sigma, D2158), 0.1%v tween-20 (Acros, 23336-2500), and 0.01% sodium azide with pH adjusted to 7.5 by NaOH. Day 7 SNaPspheres were fixed in 4% PFA for 30 minutes at room temperature then washed once with PBS. Samples were immersed in 1.5 mL microtubes filled with CUBIC solution, and placed on a gentle shaker at 37°C overnight. CUBIC was replaced every day until samples had been cleared for 72 hours. CUBIC solution was aspirated, and samples were washed twice with PBS for 30 minutes per wash. PBS was removed and replaced with blocking solution. Samples were blocked for 3 days at 37°C with gentle shaking. Blocking solution was replaced with primary stain consisting of blocking solution with Mouse anti-phospho-Vimentin (1:250; MBL, D076-3) and Chicken anti-MAP2 (1:1500; Novus, NB300-213), and returned to the 37°C shaker for 3 days. Primary stain solution was removed and samples were washed twice in PBS for 30 minutes per wash. Secondary staining solution, consisting of blocking solution plus Goat anti-Mouse Alexa488 (1:1000; Thermo, A28175) and Goat anti-Chicken Alexa647 (1:1000 ; Thermo, A21449) was applied to the samples, which were then given more days at 37°C in the gentle shaker. Samples were then washed twice with PBS for 30 minutes, then submerged in RIMS solution. After approximately 60 minutes, samples were thoroughly transparent, and imaged on a Zeiss Z1 Lightsheet.

*ZIKV propagation:* Vero cells (ATCC, CCL-81) were plated on uncoated 10 cm$^2$ dishes in Vero cell growth media (DMEM + 10% heat-inactivated fetal bovine serum). At 80-90% confluency, cells were exposed to ZIKV-Ug (ATCC, VR-1838) or ZIKV-PR (ATCC, VR-1843) diluted in HyClone Earle's 1X Balanced Salt Solution (EBSS; GE Healthcare Life Sciences, SH30029.02) at a low multiplicity of infection (<0.1; based on ATCC manufacturer's quantification of viral titer) in the minimal amount of media to cover the cells (3 ml). Cells were incubated for 1 hour at 37°C/5% $CO_2$ with gentle rocking every 15 minutes to prevent the cells from drying. After this infection period, the inoculum was removed and replaced with 12 ml of Vero cell maintenance media (DMEM + 2% heat-inactivated fetal bovine serum) pre-heated to 37°C. Two days after infection, the Vero cell conditioned media was collected and centrifuged for 10 minutes at 2000xg at room temperature to remove cell debris. The virus was aliquoted and stored at -80°C prior to quantification.

*ZIKV quantification via focus forming assay*: Vero cells were plated at 150,000 per well on uncoated 24-well plates in Vero cell growth media and incubated at 37°C/5% $CO_2$. One to two days post-plating, cells were rinsed with 1X PBS and then infected with 125 µL of virus diluted in 1X EBSS ($10^{-4}$ to $10^{-7}$) for 1 hour at 37°C/5% $CO_2$ with gentle rocking of the plate every 15 minutes. After the infection period, cells were rinsed with 1X PBS. Then, 1 mL of pre-warmed overlay media [(2.1% carboxymethylcellulose sodium salt (CMC) in DMEM and 2% HI-FBS)] was slowly added onto the monolayer of infected Vero cells. Thirty-six hours later, cells were rinsed several times with 1X PBS to remove the CMC precipitates and then fixed in 4% paraformaldehyde for 15 minutes at room temperature. Post-fixation, cells were rinsed three times with 1X PBS and then permeabilized with 0.1% Triton in 1X PBS for 10 minutes. Cells were blocked with 10% normal donkey serum for 30 minutes at room temperature followed by a 1 hour/37°C incubation in 150 µL of 1:1000 Mouse monoclonal D1-4G2 anti-flavivirus envelope protein (EMD Millipore, MAB10216) antibody diluted in blocking solution. After two washes in 1X PBS at room temperature, cells were incubated for 1 hour/37°C in 1:1000 Goat anti-Mouse HRP-conjugated secondary antibody (Abcam, ab6789) in blocking solution. Cells were once again washed twice with 1X PBS at room temperature followed by the addition of peroxidase substrate (Vector Laboratories, SK-4600) in 1X PBS. The number of foci were counted and then multiplied by the dilution factor to quantify the viral titer. Dilutions were run in quadruplicate and ZIKV-Ug and ZIKV-PR were quantified at the same time. ZIKV-Ug and ZIKV-PR were quantified as $5.5 \times 10^7$ and $4.0 \times 10^7$ focus forming units (FFU) per mL, respectively.

*Infectivity assay:* SNaPs or SNaP-derived glial cells were infected for 1 hour at 37°C/5% $CO_2$ with ZIKV-Ug or ZIKV-PR diluted in 1X EBSS at a MOI of 10, 1, 0.1, or 0.01. Vero cell conditioned media from uninfected cells was diluted in 1x EBSS and used for the mock controls. Mock and ZIKV-infected cells were fixed 54 hours post-infection (hpi) with 4% paraformaldehyde for 15 minutes at room temperature and then washed with 1X PBS. Cells were permeabilized with 0.1% Triton for 15 minutes and then blocked with 10% normal donkey serum diluted in 1X PBS for 1 hour at room temperature followed by an overnight 4°C incubation in primary antibody: Mouse monoclonal D1-4G2 anti-flavivirus envelope protein (1:500) and Rabbit anti-PAX6 (1:500; Stem Cell Technologies, 60094) antibody diluted in blocking solution. After 3 washes in 1X PBS at room temperature, cells were incubated for 2-4 hours at room temperature in secondary antibody: Donkey anti-Mouse Alexa647 (1:1000) and Donkey anti-Rabbit Alexa555 (1:1000). Cells were washed once with 1X PBS followed by a 5 minute incubation in DAPI (1:5000). For Vero cell infections, an additional 20 minute room temperature incubation with F-Actin CytoPainter Phalloidin-iFluor 555 Reagent (1:10000; Abcam, ab176756) was included. Finally, cells were washed twice more with 1X PBS prior to imaging. For each well of a 96-well plate, 4-8 fluorescent images were taken using the Cytation 3 cell imaging multi-mode reader. All images were then processed using the CellProfiler imaging analysis software to quantify the percentage of 4G2-positive PAX6 stained cells.

*Cell viability assay:* At 96 hpi, cell viability was quantified using the CellTiter Glo 2.0 kit (Promega, G9242). In brief, culture media was removed and cells were washed once in 100 µL 1X PBS. The 1X PBS was removed and replaced with 100 µL CellTiter Glo reagent. Plates were rocked gently for 2 minutes to facilitate cell lysis. After a 10 minute incubation at room temperature, luminescence was measured using the Cytation 3 cell imaging multi-mode reader. Data is presented as luminescence as a percentage of mock-infected controls.

*qPCR quantification of ZIKV*: SNaPs and SNaP-derived glial cells were infected with ZIKV-Ug or ZIKV-PR at a MOI of 10 or with mock media for 1 hour at 37°C. At 24 hpi and 72 hpi, 170 µL of conditioned media was removed from each well of a 96-well plate. 140 µL of this supernatant was flash frozen in dry ice and stored at -80°C for subsequent RNA extraction experiments, while 30 µL of the media was added directly to Vero cells for a 1 hour infection at 37°C. Vero cell infectivity was measured at 24 hpi using the infectivity assay. Viral RNA was prepared from the conditioned media using the QIAamp Viral RNA Mini Kit (Qiagen, 52906). cDNA was prepared from 10 µL of viral RNA per sample using the iScript Reverse Transcription Supermix (Bio-Rad, 1708841). At the same time, cDNA from 6 x 1:10 dilutions of the stock viral RNA of known focus-forming units per mL (FFU/mL) was prepared. qRT-PCR was conducted using the CFX96 Touch Real-Time PCR Detection System (Bio-Rad). For each sample, 1 µL of the cDNA or quantified stock cDNA was added to 5 µL iTaq Universal SYBR Green Supermix (Bio-Rad, 1725124), 400 nM primers (Integrated DNA Technologies; Coralville, IA) designed for ZIKV-Ug or ZIKV-PR (ZIKV-Ug primers: TGG GA G GTT TGA AGA GGT TG and TCT CAA CAT GGC AGC AAG ATC T; ZIKV-PR primers: GGG ACA GTC ACA GTG GAG GT and GGT GGA TCA AGT TCC AGC AT), and enough nuclease-free $dH_2O$ for a final reaction volume of 20 μL. Standard curves were established for each strain using the quantified stock dilutions and were used to assign FFU/mL values to tested samples. Due to the high concentration of virus in the supernatant samples, cDNA samples required a 1:100 dilution to fit within the acceptable CT range.

*Human antiviral response:* Mock and ZIKV-infected SNaPs (MOI = 10 for ZIKV-Ug; MOI = 20 for ZIKV-PR) were harvested at 8 hpi and 60 hpi using 350 μL of RLT Plus reagent. Total RNA was then extracted using the RNeasy Plus Micro kit (Qiagen, 74034). The $RT^2$ First Strand kit (Qiagen, 330404) was used to prepare cDNA using 400 ng for each sample. The cDNA from each sample was then diluted in 91 μl of nuclease-free $H_2O$. Then, 102 μL of cDNA was added to 1,248 μL of nuclease-free $H_2O$ and 1,350 μL of 2X $RT^2$ SYBR Green qPCR master mix (Qiagen, 330503). The master mix (10 μL) was then added into each well of a 384-well $RT^2$ Profiler PCR Array Human Antiviral Response plate (Qiagen, PAHS-122ZE-4) that contained primers for 84 genes related to the human antiviral response and five housekeeping genes that were used as internal controls (*ACTB*, *B2M*, *GAPDH*, *HPRT1*, and *RPL13A*). The data was analyzed using the online portal provided by the kit (www.SABiosciences.com/pcrarrayprotocolfiles.php).

*ZIKV infection of SNaPspheres*: Day 7-12 SNaPspheres were infected for 2 hours at 37°C/5% $CO_2$ with ZIKV-Ug or ZIKV-PR diluted in 1X EBSS at a MOI of 10, 1, or 0.1. Vero cell conditioned media from uninfected cells was diluted in 1X EBSS and used for the mock controls. SNaPspheres were imaged using bright-field microscopy with a 4X objective. Images were acquired using the Cytation 3 cell imaging multi-mode reader. All images were then stitched and processed using the CellProfiler imaging analysis software to quantify two-dimensional area.

*SNaPsphere formation assay*: Monoloyer SNaPs in a 24-well plate format were infected with ZIKV-Ug (MOI = 1). Two days later, mock and ZIKV-infected SNaPs were dissociated with Accutase and re-plated into low-attachment 6-well plates. SNaPspheres were imaged using bright-field microscopy with a 4X objective. Images were acquired using the Cytation 3 cell imaging multi-mode reader, and the number of SNaPspheres with a surface area greater than 0.1 $mm^2$ was counted and compared between conditions.

*SNaPsphere outgrowth assay:* Forty-eight hours after infection with ZIKV-Ug (MOI = 1), 10-day-old SNaPspheres were plated on Geltrex-coated 96-well plates. Two days later, SNaPspheres were fixed and immunostained with Mouse monoclonal D1-4G2 anti-flavivirus envelope protein (1:500) and Rabbit anti-TUJ1 (1:1000; Sigma Aldrich, T2200) primary antibodies, before secondary staining with Donkey anti-Mouse Alexa647 (1:1000) and Donkey anti-Rabbit Alexa555 (1:1000) antibodies. DAPI was used to counterstain DNA. SNaPspheres were imaged using bright-field microscopy with a 10X objective. Images were acquired using the Cytation 3 cell imaging multi-mode reader. Images were stitched and then processed for automated detection of neurosphere mean radius (DAPI signal) and mean neurite outgrowth (TUJ1 signal) using the CellProfiler imaging analysis software.

*Genome-wide CRISPR-Cas9 screens:* All sgRNA and lentiviral reagents for the primary and validation screens were generated at Broad Institute Genetic Perturbation Platform. SW.1 SNaPs were generated and transduced with the Brunello barcoded sgRNA library (CP0043 Brunello library containing 77,441 barcoded sgRNAs targeting 19,114 genes and 1,000 not-targeting guides) delivered through the all-in-one LentiCRISPRv2.0 system (pXPR_BRD023 vector) (Doench et al., 2016; Sanjana et al., 2014). One hundred million SNaPs per replicate (3 total replicates) were transduced using the spinoculation method, in which cells were cultured in suspension with LentiCRISPRv2.0 (estimated MOI = 0.4) and centrifuged at room temperature for 2 hours at 1,000 rpm before being plated at 120,000 cells/$cm^2$ on Geltrex coated plates. Transduced SNaPs were then expanded and selected with puromycin (1 μg/mL) for one week, at which point they were passaged onto 15 $cm^2$ Geltrex-coated dishes at 120,000 cells/$cm^2$ (40 million cells were plated per replicate to maintain the 500 cells per sgRNA representation). Two days post-plating (one day post-Y27632 removal), SNaPs were either: (1) harvested using Accutase followed by PBS washes ("Pre-infection/Day 0" samples), (2) infected with mock media (for "Mock" samples), (3) infected with ZIKV-Ug (MOI = 1), or (4) infected with ZIKV-PR (MOI = 5) in minimal media for 1 hour at 37°C/5% $CO_2$ with gentle rocking every 15 minutes to prevent the cells from drying. Cells were then fed every other day starting at 48 hpi by removing all media, washing once with 1X PBS to remove dead cells and debris, and then adding back a 50:50 fresh SNaP maintenance media/conditioned media mixture. On Day 10, all samples ("Mock/Day 10", "ZIKV-Ug", and "ZIKV-PR") were harvested and frozen at -80°C. DNA was then extracted using the QIAmp DNA Blood Maxi kit (Qiagen, 51192). PCR and sequencing were performed as previously described (Doench et al., 2016; Piccioni et al., 2018). Samples were sequenced on a HiSeq2000 (Illumina). Gene-level analysis was executed using RSA, MaGeCK, and BAGEL. RSA analysis was conducted as previously described (König et al., 2007). For RSA analysis, DESeq2 was used to generate

$\log_2$ fold change from sgRNA read counts. Subsequently, z scores were computed in R Studio (version 3.4.2) and RSA scores were generated. For additional significance thresholding, Benjamini Hotchberg correction was performed on RSA values which were plotted against Quantile 3 (Q3) and Quantile 1 (Q1) values. MaGeCK and BAGEL computations were performed on the CRISPRAnalyzer portal (http://crispr-analyzer.dkfz.de/) (Winter et al., 2017) using read counts as the input and the pre-set "Brunello" library. For this analysis, sgRNAs with fewer than 20 reads were eliminated from the analysis pipeline. SNaP fitness screen results (20 days post-library transduction; plt) were compared to previously published datasets: NSC-U5 (Day 21 plt) and NSC-CB660 (Day 23 plt).

*Generation and validation of H1 constitutive Cas9 stem cell line:* A targeting vector with AAVS1 homology arms and a Flag-Cas9-2A-Blast-BGHpA expression cassette was generated and co-electroporated with AAVS1 TALENS (System Biosciences) into H1 hESCs using the Neon Transfection System (Thermo Fisher Scientific; Waltham, MA). Two days post-electroporation, Blasticidin (4ug/mL; Thermo Fisher Scientific, R21001) was added and emerging clones were picked and analyzed by immunocytochemistry for FLAG-Cas9 using Mouse anti-FLAG antibody (1:300; Sigma Aldrich, F1804) and by PCR for proper integration into the locus across the junctions (5' junction: *AAVS1-F2* AACTCTGCCCTCTAACGCTG and *CAG-R2* CTATGAACTAATGACCCCGTAATTG; 3' junction: *BGH-F1* GGAAGACAATAGCAGGCATGC and *AAVS1-R4* CCACGTAACCTGAGAAGGGAAT; Non-targeted allele: *AAVS1-F3* CCTGGCCATTGTCACTTTGC and *AAVS1-R4* CCACGTAACCTGAGAAGGGAAT). The H1-36-23 clone was differentiated into neurons using a dual SMAD inhibition protocol and plated into 96-well plates. sgRNAs were delivered by lentiviral vectors that confer puromycin resistance, and the neurons were selected with puromycin (2 µg/mL) for 2 weeks. Neurons were lysed and next-generation sequencing of the gRNA-targeted sites was performed in order to identify and quantify indels generated.

*Validation of primary CRISPR-Cas9 ZIKV survival screen:* SNaPs were induced from H1-36-23 constitutive Cas9 stem cells before lentiviral transduction via spinoculation with individual sgRNAs (pXPR_003 and pXPR_050 vectors) in a 24-well plate format. Cells were then expanded and selected with puromycin (1 µg/mL) for one week. Cas9 sgRNA-expressing SNaPs were passaged and plated onto Geltrex at 40,000 cells per well of a 96-well plate (120,000 cells/cm$^2$). Two days later, SNaPs were infected with ZIKV-Ug (MOI = 1) before conducting the infectivity assay at 54 hpi and the cell viability assay at 120 hpi, as previously described. Infectivity and cell viability values were compared to Cas9-SNaPs that were transduced with non-targeting sgRNAs.

*Validation of primary CRISPR-Cas9 fitness screen:* To validate the genetic drivers of SNaP proliferation identified in the SNaP fitness screen, SW.1 SNaPs were transduced with Cas9-lentivirus (pLX-311-Cas9 vector) via spinoculation in 24-well plates followed by expansion and selection with blasticidin (10 µg/mL) for one week. SNaPs were then transduced with individual lentivirus sgRNAs (pXPR_003 and pXPR_050 vectors) in the 24-well plate format, before expansion and selection with puromycin (1 µg/mL) for one week. Cas9 sgRNA-expressing SNaPs were passaged and plated onto Geltrex at 1,000 cells per well of a 96-well plate (3,333 cells/cm$^2$). The following day (Day 1), Hoechst-33342 dye (1:2000 in base media) was added to 1/2 of the wells before Cytation 3 cell imaging multi-mode reader (4X objective; 3 x 3 grid). The dye and imaging process was repeated 9 days later (Day 10) for the remaining wells. All images were then processed using the CellProfiler imaging analysis software to quantify the number of Hoechst-positive cells. Data is presented as calculated doubling rate for each sgRNA line using the following equation:

$$Doubling\ time = Duration/log2(\frac{cell\ count\ at\ Day\ 10}{cell\ count\ at\ Day\ 1})$$

where *Duration* refers to time between measurements in hours (216 hours). For all experiments, the average number of cells for a given sgRNA was used as the denominator the $\log_2$ calculation (i.e. *cell count at Day 1*). Doubling rates were compared to Cas9-SNaPs that were transduced with non-targeting sgRNAs.

*Gene Ontology and pathway analysis:* Gene sets were analyzed for GO term statistical overrepresentation using the PANTHER Classification System (http://patherdb.org) with default settings. The *GO biological process complete* annotation dataset was used for the RSA-enriched proliferation and ZIKV-Ug Top 100 gene lists, while *PANTHER GO-Slim biological process* annotation dataset was used for the BAGEL (BF>10) fitness gene set. Fisher's Exact test was used for statistical analysis. Proliferation and fitness gene sets were analyzed for KEGG pathway enrichment using the Gene Set Enrichment Analysis (http://software.broadinstitute.org/gsea).

*Data analysis*: All data was analyzed and plotted using the Prism version 7.03 software (GraphPad; La Jolla, CA). Tests for multiple comparisons (i.e. Dunnett, Sidak) were chosen based on software recommendations.

**SUPPLEMENTAL REFERENCES**

Carpenter, A.E., Jones, T.R., Lamprecht, M.R., Clarke, C., Kang, I.H., Friman, O., Guertin, D.A., Chang, J.H., Lindquist, R.A., Moffat, J., et al. (2006). CellProfiler: image analysis software for identifying and quantifying cell phenotypes. Genome Biol. *7*, R100.

Piccioni, F., Younger, S.T., and Root, D.E. (2018). Pooled Lentiviral-Delivery Genetic Screens. Curr. Protoc. Mol. Biol. *121*, 32.1.1-32.1.21.

Sanjana, N.E., Shalem, O., and Zhang, F. (2014). Improved vectors and genome-wide libraries for CRISPR screening. Nat. Methods *11*, 783–784.
